# Unravelling the Graded Millisecond Allosteric Activation Mechanism of Imidazole Glycerol Phosphate Synthase

**DOI:** 10.1101/2021.10.21.464958

**Authors:** Carla Calvó-Tusell, Miguel A. Maria-Solano, Sílvia Osuna, Ferran Feixas

**Author notes:** Corresponding authors. / /.

## Abstract

Deciphering the molecular mechanisms of enzymatic allosteric regulation requires the structural characterization of key functional states and also their time evolution toward the formation of the allosterically activated ternary complex. The transient nature and usually slow millisecond timescale interconversion between these functional states hamper their detailed experimental and computational characterization. Here, we design a computational strategy tailored to reconstruct millisecond timescale events to describe the graded allosteric activation of imidazole glycerol phosphate synthase (IGPS) in the ternary complex. IGPS is a heterodimeric bienzyme complex responsible for the hydrolysis of glutamine to glutamate in the HisH subunit and delivering ammonia for the cyclase activity in HisF. Despite significant advances in understanding the underlying allosteric mechanism, essential molecular details of the long-range millisecond allosteric activation pathway of wild-type IGPS remain hidden. Without using *a priori* information of the active state, our simulations uncover how IGPS, with the allosteric effector bound in HisF, spontaneously captures glutamine in a catalytically inactive HisH conformation, subsequently attains a closed HisF:HisH interface, and finally forms the oxyanion hole in HisH for efficient glutamine hydrolysis. We show that effector binding in HisF dramatically decreases the conformational barrier associated with the oxyanion hole formation in HisH, in line with the experimentally observed 4500-fold activity increase in glutamine production. The formation of the allosterically active state is controlled by time-evolving dynamic communication networks connecting the effector and substrate binding sites. This computational strategy can be generalized to study other unrelated enzymes undergoing millisecond timescale allosteric transitions.

## INTRODUCTION

Proteins reshape their function in response to environmental changes through allosteric regulation.(1) Allostery is the process in which two distinct sites within a protein or protein complex are functionally coupled.(2) In allosterically regulated enzymes, effector binding at a distal site alters the thermodynamic and/or kinetic parameters of the catalytic reaction at the active site.(3) The transfer of chemical information between the two energetically coupled sites is mediated by structural(4) and/or dynamical(5) changes that generally make accessible the pre-organized active site conformation characteristic of the enzyme active state.(6, 7) To attain such catalytically competent state, effector binding finely tunes the enzyme dynamic conformational ensemble by reshaping the relative populations of the conformational states and/or the timescales of structural fluctuations and conformational transitions.(8) Complete bidirectional communication between distal sites occurs at the ternary complex, i.e. when both the effector and substrate are bound at their respective sites, and propagates through dynamic networks of inter- and intramolecular interactions.(9, 10) Thus studying the ternary complex conformational ensemble is essential for the complete characterization of enzymatic allosteric activations. Capturing the time evolution of the allosteric activation of enzymes toward the formation of the ternary complex involves deciphering the interplay of fast and slow conformational dynamics coupled to effector and substrate binding.(11) The transient nature of the ternary complex and the allosteric transition of enzymes undergoing turnover hampers the structural and dynamic characterization of allosteric mechanisms and the identification of functionally relevant states.(12–17) It is therefore not surprising that the allosterically active state remains hidden for several enzymes.

Allosteric regulation operating in the model *Thermotoga maritima* Imidazole Glycerol Phosphate Synthase (IGPS) has been investigated from structural and dynamical perspectives.(18-29) IGPS is a heterodimeric enzyme belonging to class I glutamine amidotransferases (GATase) that encompasses the catalytic interplay between HisH and HisF subunits (Figure 1). HisH catalyzes glutamine hydrolysis producing glutamate and ammonia. The HisF cyclase monomer couples the ammonia produced by HisH, that migrates through an internal tunnel, with N’-[(5’-phosphoribulosyl)formimino]-5-aminoimidazole-4-carboxamide ribonucleotide (PRFAR). The later also acts as allosteric effector for the reaction occurring in HisH. The binding of PRFAR, ca. 30 Å far away from the HisH active site, enhances 4500-fold the basal glutaminase activity of IGPS, while the substrate affinity is only moderately altered.(25) This is consistent with a prevalent V-type allosteric coupling between HisF and HisH subunits.(29)

**Figure 1.**
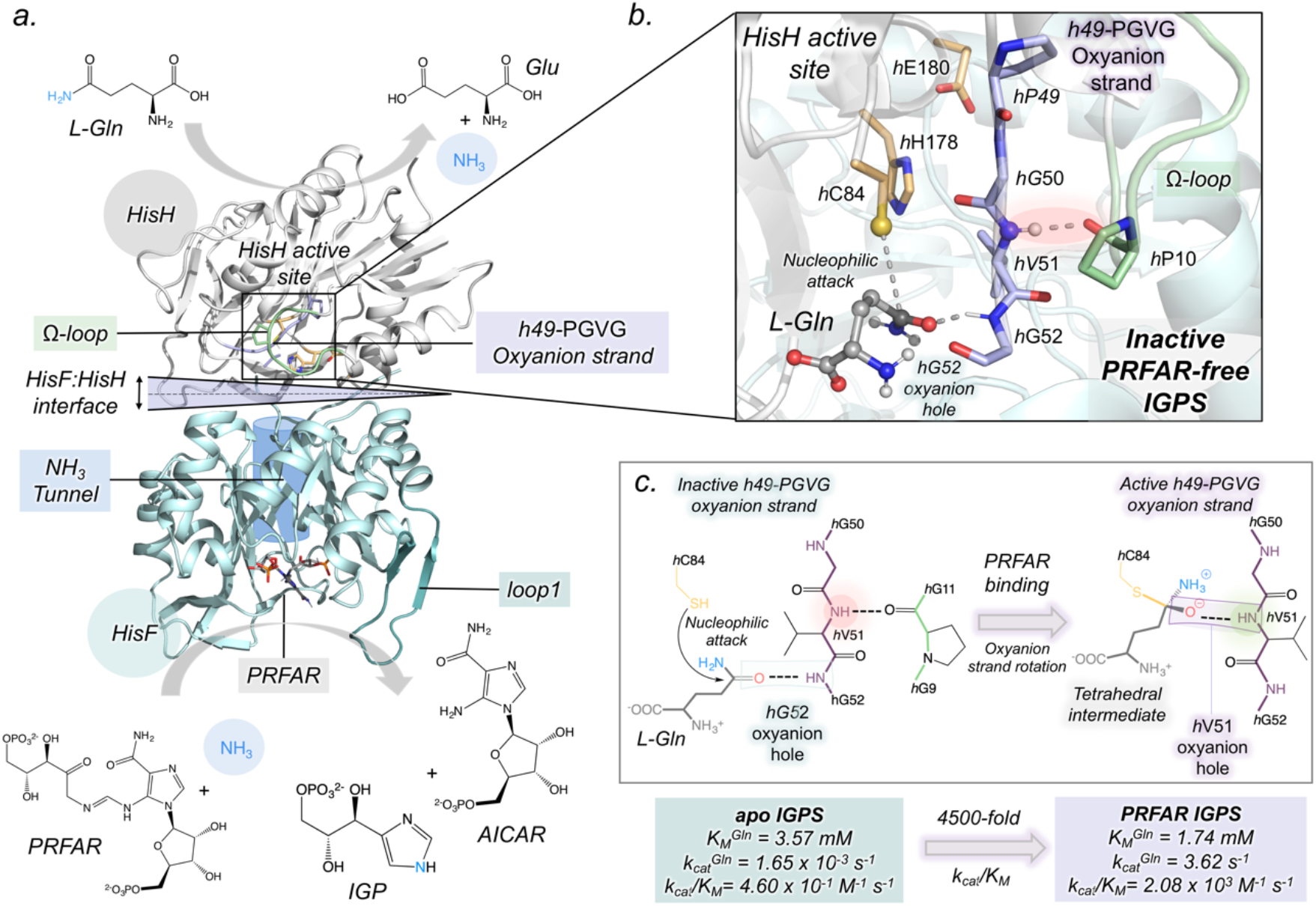
Overview of IGPS structure and global mechanism. (a) The enzyme is a heterodimeric complex formed by two subunits (PDB: 1GPW): HisH (white) and HisF (cyan). (b) HisH active site (PDB: 3ZR4) with substrate glutamine (L-Gln, gray) bound in inactive *h*49-PGVG oxyanion strand (purple). The catalytic and Ω-loop residues are highlighted in orange and green, respectively. The NH backbone of *h*V51 is shown in spheres. (c) Hypothesis of *h*V51 oxyanion hole formation and kinetic parameters for glutamine hydrolysis in *apo* and PRFAR-bound IGPS, extracted from reference (29).

From the structural perspective, it was initially hypothesized that PRFAR binding (HisF) allosterically activates IGPS through the formation of an oxyanion hole composed by the amide HN backbone of *h*V51 that pre-organizes the HisH active site for glutamine hydrolysis (*h* and *f* labels are used to highlight HisH or HisF residues, respectively).(22, 26) *h*V51 is located in the oxyanion strand, which consists of four residues (*h*49-PGVG) situated in the proximity of the catalytic triad formed by *h*C84, *h*H178, and *h*E180 (Figure 1b). Based on mechanistic observations of other GATases, the oxyanion hole is required to stabilize the transient negative charge of the tetrahedral intermediate formed during the glutaminase reaction (Figure 1c).(30) However, for the wild-type IGPS (*wt*IGPS) none of the available X-ray structures present the amide HN *h*V51 pointing toward the HisH active site, suggesting that the pre-organized HisH active site is not prevalent in IGPS conformational ensemble. An alternative hypothesis is that the tetrahedral intermediate can be stabilized by the HN of adjacent *h*G52, and that the closure of the HisF:HisH interface upon PRFAR binding are key for the allosterically-triggered enhanced catalytic activity.(27) However, the rapid glutamine turnover of the allosterically activated enzyme prevented the structural characterization of *wt*IGPS, whose active conformation remains hidden. There is also a lack of structural details of the allosteric communication pathway connecting the substrate-free IGPS with the postulated allosterically active ternary complex, either with the *h*V51 oxyanion hole formed and/or with the HisF:HisH interface productively closed.

From the dynamic perspective, PRFAR binding enhances IGPS conformational flexibility through the activation of millisecond motions that stimulate allosteric communication between the two subunits.(25) NMR experiments suggest different patterns of millisecond motions when comparing PRFAR-free, PRFAR-bound, and ternary complexes. Upon the formation of the ternary complex, an allosteric network of residues displaying correlated millisecond motions connecting HisF and HisH binding sites arises. Recent NMR experiments of the *h*C84S mutant, displaying drastically reduced glutaminase activity, suggested that inactive and active states are in dynamic equilibrium.(31) However, the active state was not detected in *wt*IGPS under turnover conditions, being below the detection limit of NMR experiments, thus concluding that the *h*C84S mutant stabilizes the active conformation. Nanosecond molecular dynamics (MD) simulations indicated that the effect of PRFAR binding propagates from *f*loop1 through a network of salt bridges connecting HisF and HisH subunits.(26) This results in altered dynamics of the HisF:HisH interface and the weakening of a hydrogen bond between *h*P10 located at the Ω-loop and the HN of the oxyanion strand *h*V51 (Figure 1b). However, the predicted millisecond time-scale rotation of the oxyanion strand was not captured by the sub-microsecond MD simulations performed in all previous studies. Different studies using dynamical network models revealed the enhancement of HisF:HisH interdomain communication in the presence of PRFAR.(26, 32–38) Given the difficulties to computationally study slow millisecond time-scale events, any of the previous dynamical network studies was carried out in the active IGPS state. Despite significant advances in the understanding of the underlying allosteric mechanism operating in IGPS, the sequence of molecular events of how substrate binding couples with allosteric activation and HisF:HisH interdomain motions toward the formation of the active ternary complex of IGPS remain unknown. The elucidation of the complete millisecond time-evolution of the allosteric activation of IGPS at atomic resolution is crucial as it harbors essential information for the enzyme function and engineering.

In this study, we characterize the molecular details of the graded allosteric activation of *wt*IGPS and identify hidden states relevant for IGPS catalytic activity with a computational strategy tailored to explore millisecond timescale events (Figure S1). Our approach focused on long timescale molecular dynamics simulations, enhanced sampling techniques, and dynamical networks captures, without using *a priori* information of the active state, the time-evolution of the allosterically-driven conformational ensemble toward the presumed active state of PRFAR-bound IGPS. This study thus uncovers the HisH active site pre-organized with the *h*V51 oxyanion hole properly oriented to stabilize the substrate glutamine, in both substrate-free and ternary complex. Spontaneous substrate binding simulations and dynamical network-analysis reveal a delicate coupling between substrate binding and IGPS conformational dynamics that fine tunes correlated motions through the allosteric activation of IGPS ternary complex. We find that the productive closure of the HisF:HisH interface is a prerequisite to effectively populate the pre-organized HisH active site upon the formation of the ternary complex.(19, 27) During the elaboration of the present manuscript, Wurm and coworkers successfully captured through X-ray crystallography the allosterically activated conformation of a catalytically inactive *h*C84A IGPS mutant bound to PRFAR precursor and L-Gln substrate.(31) This *h*C84A IGPS structure presents a closed HisF:HisH interface and the *h*V51 oxyanion hole formed, as predicted by our simulations, thus providing experimental evidence to the allosteric activation observed here through an *in silico* approach. This computational strategy can be generalized to decipher allosteric mechanisms of unrelated enzymes, which is of interest for enzyme design and drug discovery.(39, 40)

## RESULTS

### Effect of PRFAR binding in IGPS: structural characterization of transient hV51 oxyanion hole formation in HisH

PRFAR binding in HisF is postulated to influence the HisH *h*49-PGVG oxyanion strand conformational dynamics, making accessible the pre-organized active site with the *h*V51 oxyanion hole formed. To elucidate the impact of PRFAR, we performed microsecond conventional molecular dynamics (cMD) simulations of IGPS in both *apo* (neither L-Gln substrate nor PRFAR are bound) and PRFAR-bound states, starting all simulations from an inactive IGPS conformation (SI Methods). MD simulations reveal that PRFAR induces significant changes in the orientation and conformational dynamism of HisH oxyanion strand motif, even in the absence of substrate.

The conformational landscape that illustrates the different orientations of the oxyanion strand is reconstructed from ten replicas of 1.5 μs cMD simulations for both *apo* and PRFAR-IGPS. The coordinates selected to capture relevant *h*49-PGVG conformations are the *ϕ* dihedral angles of *h*V51 and *h*G50 (Figures 2a and S2-S3). Far from the uniformity observed in available X-ray structures, our simulations show that the oxyanion strand presents the ability to sample two major orientations in the *apo* state: the inactive-OxH (given that the oxyanion strand is not properly oriented for catalysis), and another conformation with the oxyanion strand unblocked not previously observed in any X-ray (Figure 2b). In the inactive-OxH state, the carbonyl group of *h*G50 is pointing toward the catalytic *h*C84 while the amide HN backbone of *h*V51 is oriented toward the carbonyl backbone of *h*P10 at the Ω-loop, establishing a stable *h*Pro10-*h*V51 hydrogen bond that blocks the rotation of the *h*V51 backbone. As in most IGPS X-ray structures, the side chain of *h*V51 points toward the HisH active site. The unblocked-OxH state is characterized by a partial rotation of *ϕ*-*h*G50, while *ϕ*-*h*V51 gains some flexibility. This enhanced dynamism of the oxyanion strand is triggered by the complete breaking of the *h*Pro10-*h*V51 hydrogen bond that induces the separation of the oxyanion strand from the Ω-loop (Figure S4). Because of these motions, the side chain of *h*V51 is displaced from the HisH active site (Figure 2b). The analysis of individual cMD trajectories show that the inactive-OxH and unblocked-OxH states can interconvert in the microsecond timescale (Figures S2). However, the formation of the *h*V51 oxyanion hole is not observed in these *apo* μs-cMD simulations, thus suggesting a rather high energy barrier associated with the oxyanion strand transition in the absence of PRFAR.

**Figure 2.**
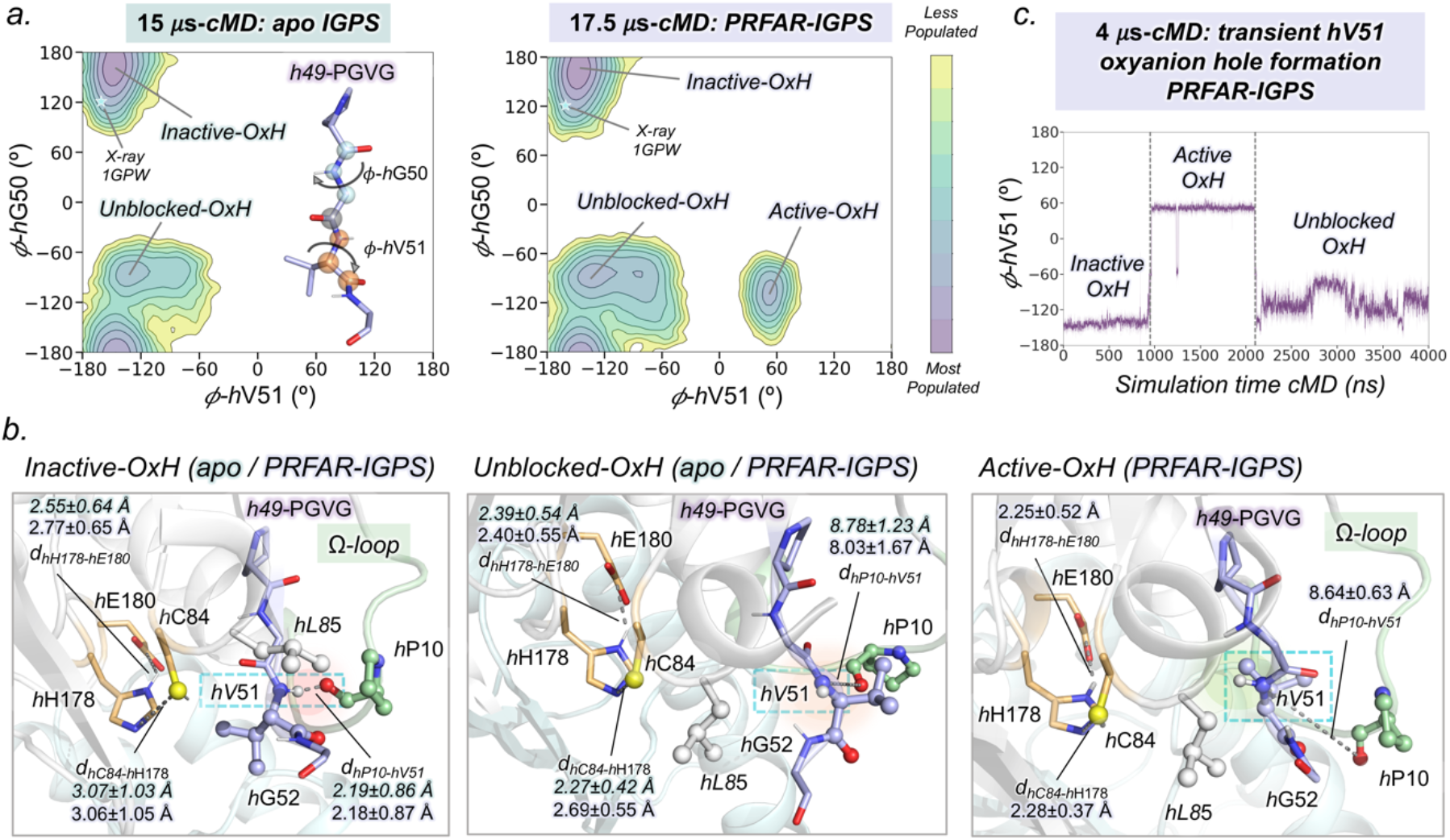
Conformational landscape of *h*49-PGVG oxyanion strand obtained from conventional molecular dynamics (cMD) simulations. (a) Conformational landscape of *apo* and PRFAR-IGPS constructed using the *ϕ* dihedral angles of *h*V51 and *h*G50. The cyan star symbol indicates the *h*V51 and *h*G50 of X-ray IGPS structure (1GPW chain A/B) used as starting point in cMD simulations. (b) Representative HisH active site structures of most populated states in the PRFAR-IGPS conformational landscape (2a). The NH backbone of *h*V51 is highlighted inside a cyan dashed box. Average distances (in Å) are depicted in green and purple for *apo* and PRFAR-bound states, respectively. (c) Transient *h*V51 oxyanion hole formation observed in a cMD trajectory of 4 μs.

Interestingly, the presence of PRFAR favors the exploration of an extra state in the *h*49-PGVG conformational landscape (Figure 2a). This state is characterized by the complete rotation of *ϕ*-*h*V51 with respect to the inactive IGPS X-ray structure used as starting point (Figure 2b), thus demonstrating that the oxyanion hole formed conformation (named active-OxH) is accessible even in the absence of L-Gln substrate, as observed in other GATases.(41) MD simulations therefore reveal the ability of PRFAR-IGPS to form the *h*V51 oxyanion hole, which is consistent with the hypothesis that PRFAR stimulates changes in the HisH active site despite being located 30 Å away. This hidden active-OxH state of the oxyanion strand shows remarkable similarities with X-ray structures of L-Gln-bound *h*C84A IGPS variant and other GATases presenting the equivalent oxyanion hole formed (Figure S5).(30, 31) In particular, the HN backbone of *h*V51 is pointing towards the catalytic *h*C84 and the *h*Pro10-*h*V51 hydrogen bond observed in the previous inactive-OxH conformation is clearly broken (see Figure 2b). Moreover, the catalytic triad is significantly more stable than in inactive-OxH and unblocked-OxH conformations (Figures 2b and S4). In both active-OxH and unblocked-OxH states, the bulky side chain of *h*L85 occupies the space left in the HisH active site by the *h*V51 side chain, blocking the access to the catalytic *h*C84 (Figure 2b and S6). As we will discuss later, these HisH active site rearrangements have direct implications in substrate binding and IGPS allosteric activation.

The *h*V51 oxyanion hole formation occurs only in 1/10 replicas of 1.5 μs MD simulations, indicating that it is a rare event in the microsecond timescale. The cMD trajectory where the oxyanion strand completely rotates was extended up to 4 μs (Figures 2c and S7 and Movie S1). In this trajectory, a transient *h*V51 oxyanion hole formation occurs after 1 μs of simulation time, remaining formed for around 1 μs, and subsequently evolving to unblocked-OxH conformation. Based on μs-cMD, the active-OxH conformation represents a high energy state in the conformational ensemble of PRFAR-IGPS. The transient μs-formation of the *h*V51 oxyanion hole, induced by PRFAR, suggests a lower barrier for the interconversion between the inactive-OxH and active-OxH states with respect to *apo* IGPS. Most importantly, these results demonstrate that the postulated pre-organized HisH active site pre-exists in solution for *wt*IGPS in the presence of PRFAR and in the absence of L-Gln substrate.

### IGPS captured with a closed HisF:HisH interface

The μs-cMD simulations performed in the previous section successfully caught the pre-organized HisH active site and associated IGPS conformational changes (SI Extended Text and Figures S2-S10), still millisecond-timescale events dominating IGPS allosteric communication are not completely captured. To unravel the effects of PRFAR beyond microsecond timescales, we performed substrate-free accelerated molecular dynamics (aMD) simulations for *apo* and PRFAR-bound states (10 replicas of 1 μs performed, SI Methods).(42, 43) In general, microsecond aMD simulations provide sufficient unconstrained enhanced conformational sampling to make accessible millisecond timescale events typical of allosterically regulated systems.(44) Indeed, our aMD simulations show multiple infrequent and short-lived formations of the *h*V51 oxyanion hole in the presence of PRFAR, i.e. the active-OxH state is explored, and any transition in the *apo* state (Figures S11-S12). These results again suggest that, within the substrate-free IGPS conformational ensemble, the active-OxH state presenting the *h*V51 oxyanion hole formed is substantially higher in energy than the inactive-OxH conformation.

Additionally, aMD simulations show significant global conformational changes of IGPS. PRFAR releases tension in the interdomain region facilitating both the rotation and closure of the HisF:HisH interface (Figure 3 and S13). In fact, these simulations unveil a displacement of the conformational ensemble toward closed states of the IGPS interface (in most inactive IGPS X-ray structures the HisF:HisH interface angle (θ) is *ca*. 25º). Interestingly, a closed metastable state is captured displaying an average interface angle of around 11º (in substrate-bound *h*C84A IGPS, HisF:HisH(θ) is *ca*. 10º).(30) This productive closure is stabilized by the formation of a hydrogen bond between the backbones of *h*H53 and *f*T119 that restrains the HisF:HisH opening-closing motion. This results in the Ω-loop collapsing over *f*α3 as *h*α1 and *f*α3 become perfectly aligned, enhancing HisF:HisH communication suggested to be key for productive catalysis.(45, 46). Importantly, aMD simulations indicate that, without the L-Gln substrate, the productive closure of the HisF:HisH interface is not correlated with the formation of the catalytically relevant *h*V51 oxyanion hole (Figure S14). Similar closed states were also sampled in the *apo* state simulations, although less frequently, showing that IGPS interface closure is not an exclusive allosteric effect elicited by PRFAR (Figure S13).

**Figure 3.**
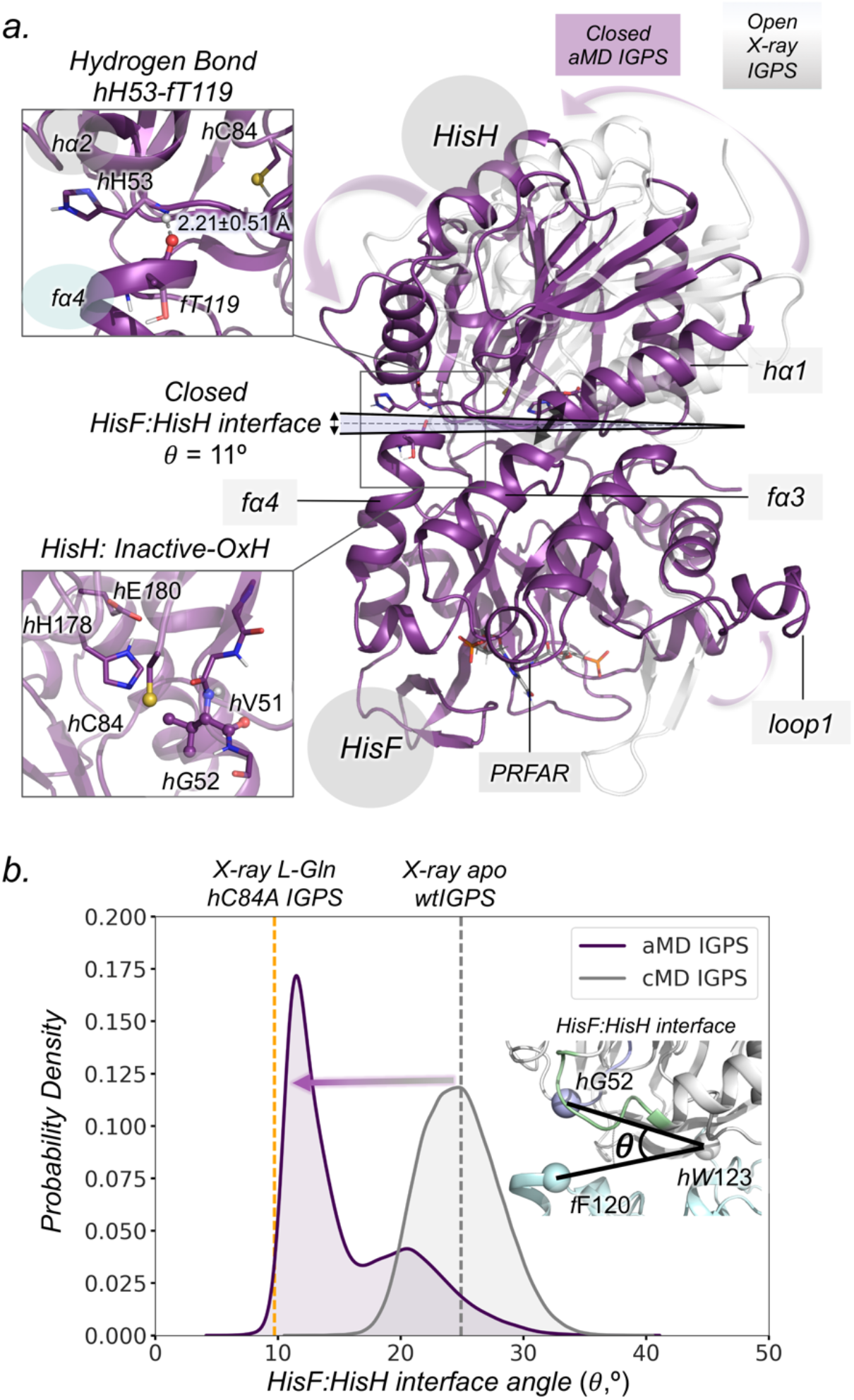
Accelerated Molecular Dynamics (aMD) simulations: Identification of IGPS productive closure. (a) Structural comparison of open (in gray, PDB: 1GPW (chains A/B)) and closed HisF:HisH interfaces obtained from aMD simulations (in purple). The hydrogen bond between the backbones of *h*H53 and *f*T119 that stabilizes the closed HisF:HisH interface and the conformation of the HisH active site are depicted. (b) Probability density distribution for HisF:HisH interface angle obtained in cMD and aMD simulations (Figure S13 for more details). The angle (8) of the HisF:HisH interface is calculated from the alpha-carbons of *f*F120, *h*W123 and *h*G52. The vertical dashed orange and gray line corresponds to the to the *h*C84A IGPS (PDB: 7AC8 (chains E/F)) and *wt*IGPS (PDB:1GPW (chains A/B)) X-ray HisF:HisH interface angles, respectively.

### Molecular basis of substrate binding in IGPS: glutamine binding occurs in the inactive-OxH state in both PRFAR-free and PRFAR-bound

The next open question is how the intrinsic μs-ms conformational dynamics of IGPS described in previous sections is coupled to L-Gln binding toward the formation of the allosterically activated ternary complex. The comparable *K*M^L-Gln^ mM values for both PRFAR-free and PRFAR-bound IGPS suggests infrequent binding events at low concentration.(25) To reconstruct the spontaneous substrate binding pathways coupled with IGPS conformational dynamics, we devised a strategy that consists in positioning a single L-Gln molecule *ca*. 25 Å away from the HisH active site and subsequently running multiple replicas of aMD simulations starting from the three different IGPS conformations (active-OxH, inactive-OxH, and unblocked-OxH) sampled in the substrate-free cMD simulations (Figures 4a and S15). We find that unconstrained aMD simulations with an accumulated time of 36 μs (60 replicas of 600 ns of aMD performed, SI Methods) provide enough conformational sampling to capture the spontaneous binding of L-Gln from the solvent to the HisH active site in both PRFAR-free and PRFAR-IGPS.

**Figure 4.**
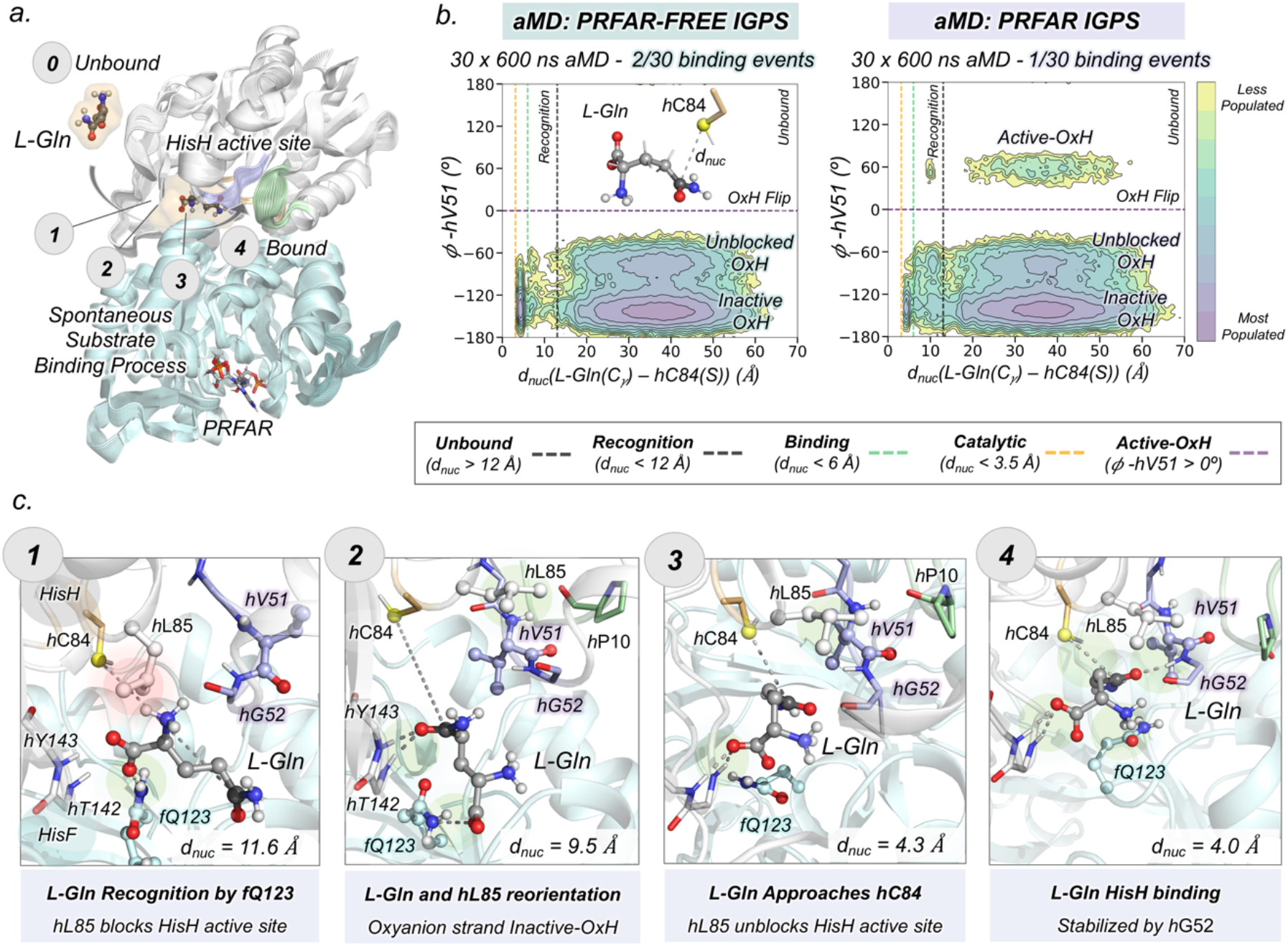
Molecular basis of L-Gln binding in IGPS. (a) General scheme of spontaneous substrate binding process in the PRFAR-free and PRFAR-IGPS states. The numbers indicate the most relevant steps of the binding process. (b) Conformational landscape obtained from the nucleophilic attack distance (*d*_*nuc*_) between the thiol group of catalytic *h*C84 (in yellow) and the amide carbon of L-Gln (in black), and the *ϕ* dihedral angle of *h*V51. The purple horizontal dashed line indicates the oxyanion strand flip, and the gray, green and orange vertical dashed lines indicate the distance where recognition, binding, and catalysis take place. (c) Molecular representation of selected key conformational states of the L-Gln binding pathway. The substrate is shown in gray, the oxyanion strand residues in purple, the catalytic residues in orange, the Ω-loop in green and the other HisH and HisF residues in white and cyan, respectively.

In line with experimentally reported *K*M values, L-Gln binding only occurs in 2/30 and 1/30 replicas in PRFAR-free and PRFAR-bound, respectively, thus indicating that substrate binding is indeed an infrequent event (Figure S16). To evaluate the binding process coupled with the oxyanion strand conformational dynamics, we collectively represented all aMD simulations using two coordinates: the nucleophilic attack distance (*d*_*nuc*_) between the thiol group of catalytic *h*C84 and the amide carbon of L-Gln, and the *ϕ* dihedral angle of *h*V51 (Figure 4b). We consider that L-Gln is captured by HisH active site when *d*_*nuc*_ is below 6 Å, while catalytically productive distances will be only sampled when both the *d*_*nuc*_ is shorter than 3.5 Å and the active-OxH state (*ϕ*-*h*V51 *ca*. 60º) is attained. The binding conformational landscape shows that the bottleneck of the binding process is the recognition of the substrate at the HisF:HisH entrance channel, which corresponds to a long *d*_*nuc*_ (above 12 Å). The difficulties to capture the substrate may be associated with the polarity of L-Gln and intrinsic HisF:HisH interface fluctuations. In PRFAR-free simulations, binding readily occurs upon recognition while, in PRFAR-IGPS, L-Gln binding is controlled by the oxyanion strand conformation. Interestingly, L-Gln binding (*d*_*nuc*_ below 6 Å), only occurs when the oxyanion strand attains the inactive-OxH conformation (*ϕ*-*h*V51 within [-180º,-100º]), irrespective of the starting orientation. The ability of L-Gln to bind only the inactive-OxH conformation in the presence of PRFAR is in line with recent solution NMR experiments of the *h*C84S IGPS variant.(31)

From the independent aMD trajectories, we can reconstruct the sequence of events of L-Gln binding at the molecular level, and also identify the associated key conformational states of IGPS (Movies S2-S3). The detailed analysis of the L-Gln binding pathway along a representative PRFAR-IGPS aMD simulation (starting from an IGPS conformation with the active-OxH oxyanion-strand and open HisF:HisH interface) is shown in Figures 4c and S17-S19. L-Gln is first recognized by *h*α2 and *f*α4 interface residues. Upon initial recognition, the HisF:HisH interface expands up to 30º, subsequently capturing L-Gln. The carboxylate group of L-Gln is then rapidly recognized by the side chain of HisF residue *f*Q123 (step 1). At this point, the active-OxH conformation of the oxyanion strand prevents the access of L-Gln close to the catalytic *h*C84 as the side chain of *h*L85 blocks its entrance. This blockage by *h*L85 leads to rather long nucleophilic attack *h*C84-Gln distances of near 10 Å. Subsequently, the oxyanion strand readily transitions from the catalytically-relevant active-OxH to the unblocked-OxH and inactive-OxH orientations. The population of inactive-OxH displaces the *h*L85 side chain from the active site, allowing the reorientation of the substrate in the HisH active site entrance: the carboxylate of L-Gln is stabilized by *h*T142 and *h*Y143 backbones and the side chain of *h*Q88, while the L-Gln amino group establishes a salt bridge with the side chain of *h*E96 (step 2 in Figure 4c and S18). This reorientation is rapidly followed by the extension of the side chain of L-Gln closer to the *h*C84 catalytic residue (steps 3 and 4).

When L-Gln eventually binds the HisH active site in the inactive-OxH state (step 4), the carbonyl of L-Gln is stabilized by the HN backbone of *h*G52 (2.47 ± 1.07 Å) and the nucleophilic attack Gln-*h*C84 distance is still rather long (i.e. 4.72 ± 0.67 Å). At the same time, the side chains of *h*L85 and *f*Q123 stabilize the side chain of L-Gln. At this point, catalysis cannot readily occur as the oxyanion strand is not in the catalytically active-OxH conformation for exploring short Gln-*h*C84 distances required for efficient glutamine hydrolysis (see next section below). Interestingly, the substrate-bound pose predicted from the spontaneous binding aMD simulations perfectly overlays with the X-ray structure of L-Gln-bound PRFAR-free IGPS (Figure S20). It should be mentioned that a similar L-Gln binding pose is obtained in PRFAR-free simulations, which again occurs in the inactive-OxH conformation of the oxyanion strand (Figures S20-S22). In all cases, unbinding events are not observed when the hydrogen bond between L-Gln and *h*G52 is established. Upon L-Gln binding in the inactive-OxH state, the reorientation of the oxyanion strand to form the active-OxH state is not observed within the 600 ns aMD simulation, thus indicating that the complete allosteric activation has still not taken place, in line with the millisecond timescale associated with this transition. Still, these aMD simulations indicate that PRFAR is not significantly altering the rates of the initial steps toward the formation of the active ternary complex, as we observe L-Gln binding both in the presence and absence of PRFAR with similar probabilities. At this point, one question remained unanswered: How does the coupled effect of L-Gln and PRFAR binding trigger the sequence of events that allosterically activate IGPS for glutamine hydrolysis?

### Time evolution toward the active ternary complex: IGPS caught in the allosterically active state

Next, we explored how the formation of the IGPS ternary complex alters the oxyanion strand conformational dynamics and the IGPS conformational ensemble with respect to the substrate-free form. The intriguing question that is still open is whether the presence of L-Gln in the HisH active site facilitates the *h*V51 oxyanion hole formation in the IGPS ternary complex. NMR studies indicate that millisecond motions associated with the allosteric activation are significantly perturbed in the ternary complex in comparison with PRFAR-IGPS.(25) To properly capture the complete millisecond allosteric activation of *wt*IGPS, we extended the aMD simulations from the previously obtained substrate bound pose (L-Gln bound in the inactive-OxH conformation), for both PRFAR-free and PRFAR-bound. The formation of the *h*V51 oxyanion hole in the ternary complex was evaluated by monitoring the orientation of the *h*49-PGVG oxyanion strand from five replicas of aMD simulations that accumulated a total of 30 μs (Figures 5 and S23-S24). Our aMD simulations of the PRFAR-IGPS ternary complex reveal that the formation of *h*V51 oxyanion hole is accessible in the presence of L-Gln in the HisH active site (Movie S4). Long-lived formations of the active-OxH state are observed without unbinding of L-Gln in 2/5 replicas of aMD, thus indicating that the oxyanion strand transition is a frequent event in the L-Gln bound form.

**Figure 5.**
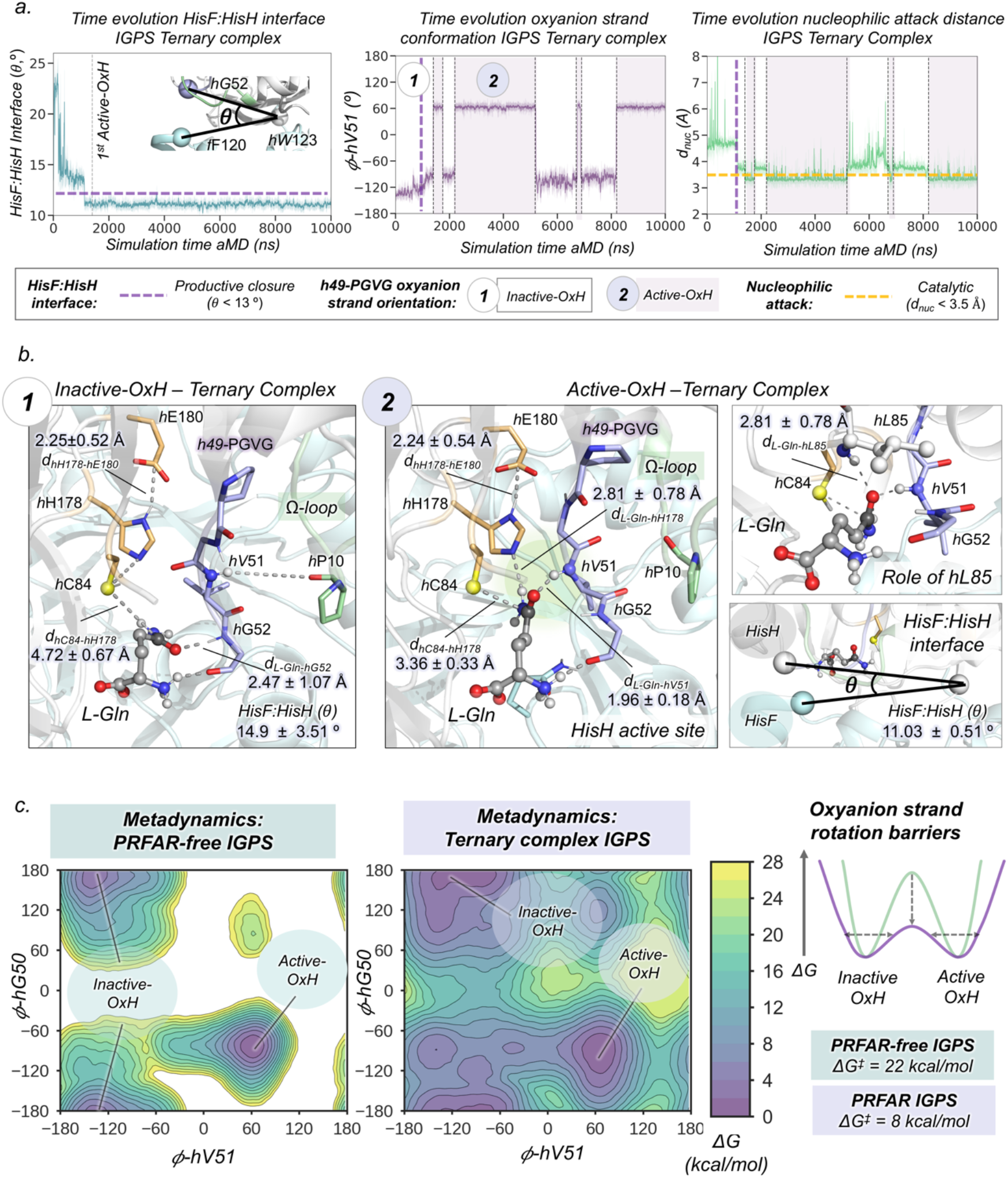
Allosteric activation of IGPS in the ternary complex. (a) Plot of the HisF:HisH interface angle along the 10 μs-aMD simulations. Plot of the *h*V51 dihedral angle along the 10 μs-aMD simulations. Plot of the distance corresponding to the nucleophilic attack along the 10 μs-aMD simulations. The purple dashed line indicates when productive closure takes place, purple regions indicate when Active-OxH state is populated, and white regions indicate when Inactive-OxH state is populated. (b) Representative structures of the Inactive-OxH and Active-OxH states sampled in the aMD simulations of the IGPS ternary complex. The role of *h*L85 is shown for Active-OxH. (c) Free energy landscape of the *h*49-PGVG in the PRFAR-free and PRFAR-IGPS states obtained from well-tempered metadynamics simulations. Representative scheme of the oxyanion strand interconversion barrier in the PRFAR-free (green) and PRFAR (purple) states. The presence of PRFAR decreases the interconversion barrier and broadens the energy minima.

Analyzing the independent aMD trajectories, we can determine the sequence of events that occur upon L-Gln binding to identify the molecular basis of the allosteric activation mechanism in the ternary complex (Figure 5a and S25-26). In all cases, the *h*V51 oxyanion hole formation is preceded by significant changes in the HisF:HisH interface. After the substrate is captured in the inactive-OxH, the HisF:HisH(θ) angle decreases from 20º to 15º. This partial closure is, however, not directly affecting the nucleophilic attack Gln-*h*C84 distance that remains around 4.5 Å. After 1 μs of aMD simulation time, the productive closure of the HisF:HisH interface takes place (HisF:HisH(θ) below 13º). Coupled to this collective motion of both IGPS subunits, the amide backbone of *h*H53 establishes a hydrogen bond with the carbonyl backbone of *f*T119, the Ω-loop collapses over *f*α3, and the *h*α1 and *f*α3 become perfectly aligned. The productive closure brings the nucleophilic attack distance down to 4 Å, while the *h*V51 oxyanion hole remains unformed. Subsequently, *h*Pro10-*h*V51 hydrogen bond completely breaks and the HisF *f*Q123 residue positions near the substrate, enhancing the communication between the HisF:HisH subunits through the L-Gln substrate. The closure of the interface enhances the flexibility of *ϕ*-*h*V51 triggering the rotation of the oxyanion strand. When the *h*V51 oxyanion hole is formed, the nucleophilic attack *h*C84-Gln catalytic distance decreases down to 3.4 Å (Figure 5a). The closed HisF:HisH interface is significantly stable eventually adopting an open conformation beyond 10 μs of aMD, thus pointing out a slow transition between open and closed states of IGPS in the presence of L-Gln (Figure S28). This indicates that the substrate helps stabilizing the closed conformation of IGPS, which might be important to retain L-Gln in the active site during catalysis and favor ammonia transfer through the HisF tunnel.

The structural characterization of the HisH active site pre-organized with the *h*V51 oxyanion hole formed with L-Gln bound displays a similar conformation of the oxyanion strand as the one identified previously in the substrate-free form (Figure 5b). In the active-OxH ternary complex, the substrate links catalytic and oxyanion strand residues through an extensive network of non-covalent interactions. The formation of the active-OxH is coupled to the reorientation of the substrate amide group, which now presents the carbonyl oxygen stabilized by the HN backbone of *h*V51 (1.96 ± 0.18 Å) instead of *h*G52 (now at 3.87 ± 0.55 Å) and the amino group of L-Gln pointing toward the catalytic *h*H178 (2.81 ± 0.78 Å). Simultaneously, the electrophilic carbon of L-Gln moves closer to the nucleophilic thiol group of *h*C84, which now explores much shorter catalytically competent distances of 3.36 ± 0.33 Å. The carbonyl group of L-Gln is further stabilized by the HN backbone of *h*L85 (2.47 ± 0.39 Å) that completes the oxyanion hole together with HN *h*V51. Overall, non-covalent interactions between L-Gln and active site residues are enhanced when transitioning from inactive-OxH to active-OxH states (Figure S25). Moreover, a more preorganized arrangement of catalytic residues is also observed. All these rearrangements can facilitate the nucleophilic attack, proton transfer, and subsequent stabilization of the tetrahedral intermediate required for efficient glutamine hydrolysis. More importantly, this active-OxH conformation of *wt*IGPS revealed by means of extensive aMD simulations presents significant similarities with the recently obtained allosterically activated *h*C84A IGPS X-ray structure (Figure S29).(31) Altogether, μs-aMD simulations unraveled, without using *a priori* information of the active state, the catalytically competent pose corresponding to the allosterically active ternary complex of *wt*IGPS.

The striking allosteric event observed (i.e. *h*V51 oxyanion hole formation) occurs up to four times within the 10 μs aMD simulation (Figure 5a). Thus, IGPS in the ternary complex has the ability to transition between the two dominant states of the oxyanion strand upon productive HisF:HisH closure (inactive-OxH and active-OxH) indicating a lower energy barrier for the interconversion once IGPS is allosterically activated by PRFAR. The presence of L-Gln in the active site clearly stabilizes the *h*V51 oxyanion hole conformation in comparison to the infrequent transient formations observed in the substrate free-form, thus inducing a population shift towards the active-OxH state. However, the inactive-OxH remains quite stable in the conformational ensemble because when populated strong networks of interactions are established with *h*G52 and other active site residues. Moreover, we observed only one *h*V51 oxyanion hole formation in 1/5 replicas of PRFAR-free aMD simulations (Figure S23). However, in this particular case, the *h*V51 oxyanion hole remains formed for the rest of the simulation time (up to 7 μs of aMD), thus indicating a higher barrier for the oxyanion strand interconversion.

To carefully estimate the energy profile of the oxyanion strand reorientation in the presence of L-Gln in both PRFAR-free and PRFAR-bound states, we performed well-tempered metadynamics (WT-MetaD) simulations using *ϕ*-*h*V51 and *ϕ-h*G50 as collective variables (Figure 5c and SI Methods). We relied on the multiple-walkers approach using ten conformations (walkers) as starting points taken from the aMD simulations that encompass global and local features of the allosterically inactive-OxH and active-OxH states (Figure S27). The output information from all walkers was used to completely reconstruct the free energy landscape (FEL) of the *h*49-PGVG oxyanion strand conformational dynamics (Figure 5c). The FEL shows remarkable differences in the PRFAR-free and PRFAR-bound states. In PRFAR-bound, the *h*V51 oxyanion hole formation presents a barrier of *ca*. 8 kcal/mol, while in PRFAR-free this value rises to 22 kcal/mol. These results clearly indicate that the formation of the active-OxH state is a much slower step in the absence of PRFAR. It is also worth mentioning that the relative stability between the two oxyanion strand orientations, i.e. inactive-OxH and active-OxH, is preserved, which indicates that both states are similarly populated in the ternary complex, thus playing an important role along the IGPS catalytic cycle.

### Activation of correlated motions: unraveling the allosteric activation mechanism of IGPS

After capturing IGPS in the allosterically active state, the next question that we aimed to address is how PRFAR and L-Gln activate correlated motions in the ternary complex that control the HisF:HisH interface and oxyanion strand conformational dynamics. To trace down the allosteric communication pathways that interconnect HisF and HisH active sites, we dissected the allosteric activation process analyzing the time evolution of dynamic-networks of residues displaying correlated motions with the shortest-path map (SPM) tool.(47) To capture the changes on the residue-correlations along the relevant steps of the allosteric activation, we split the SPM analysis of aMD trajectories in concatenated time spans of 600 ns (i.e. from 0-600 ns, from 300-900 ns, from 600-1200 ns, etc) in what we call time-dependent SPM (td-SPM). The td-SPM analysis of the 5 μs aMD simulation that captured substrate binding, HisF:HisH productive closure, and subsequent active-OxH formation reveals a fine-tuning of correlated motions and dynamic-networks toward the allosteric activation of IGPS in the ternary complex (Figures 6 and S30-31).

**Figure 6.**
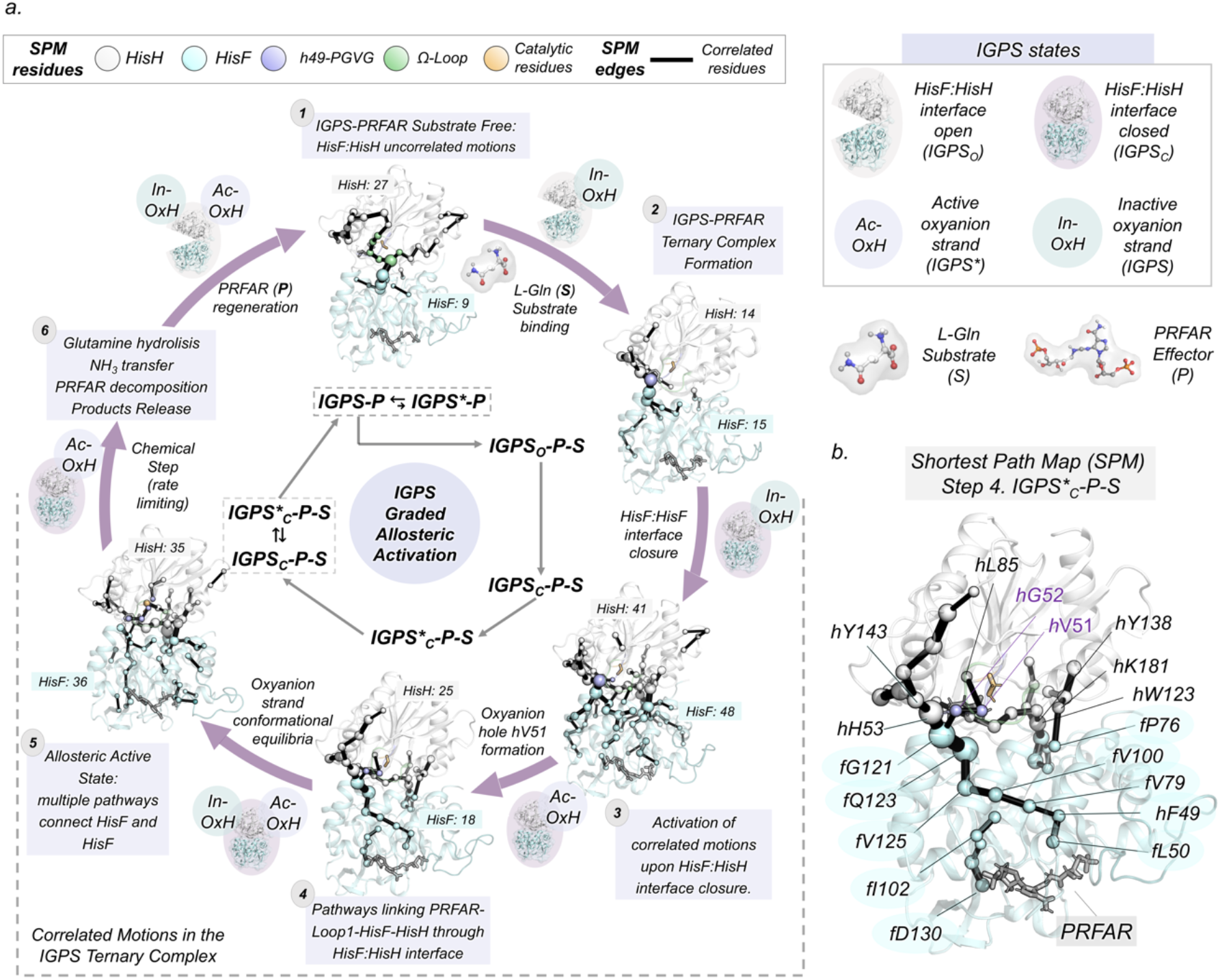
General scheme of the IGPS graded allosteric activation. (a) Time-dependent Shortest-Path Map analysis along the key states of the activation pathway. The sizes of the spheres and black edges are indicative of the importance of the position for the IGPS conformational dynamics. HisF (cyan), HisH (white), oxyanion strand (purple), Ω-loop (green), and catalytic (orange) residues are depicted in different colors. PRFAR, L-Gln, and catalytic *h*C84 are represented in sticks. (b) SPM map of the IGPS active ternary complex corresponding to step 4.

In the substrate-free form (step 1 in Figure 6), most of the correlated motions are located in the HisH subunit and HisF:HisH interface, involving connections with key residues for allosteric activation: *f*K99, *f*D98, *f*I93, *h*N15, or *h*P10. When L-Gln is captured in the HisH active site and the HisF:HisH(θ) reaches *ca*. 15º (step 2), the communication between the oxyanion strand (*h*G52 and *h*H53) and *f*α4 residues (*f*G121 and *f*S122) is enhanced. After productive HisF:HisH closure occurs (step3, *ca*. 1.1 μs), concerted motions between both subunits are significantly enhanced and multiple pathways connecting the interface of HisF and HisH arise. This includes connections from *h*H53 to *f*L153, catalytic *h*E180-*h*K181-*f*P76-*f*I75, or the anchor *h*W123-*f*A3. Interestingly, when active-OxH formations start taking place (step 4, *ca*. 1.4 μs), multiple allosteric pathways through HisF and HisH active sites arise. The connection between both sites points out the existence of functional correlated motions at the ternary complex. This long-range HisF:HisH communication dynamically couples HisH residues of the oxyanion strand including *h*V51, catalytic residues *h*C84 and *h*H178, with HisF PRFAR binding site residues: *f*V12, *f*L50, *f*I102, *f*L222. Several relevant allosteric paths are identified connecting the PRFAR site with the glutaminase HisH site, including extensive networks of hydrophobic residues at HisF: *f*L50, *f*F49, *f*V79, *f*V100, *f*V125, *f*Q123, *f*G121 and HisH residues: *h*G52, and *h*V51 (Figure 6b). When the oxyanion hole is no longer formed (step5, *ca*. 5 μs), a new network of residues appears that connect the PRFAR active site with the interdomain HisF:HisH region. These involve HisF residues: *f*D130, *f*L169, *f*I199, *f*A220, *f*R5, *f*D45, *f*D98 and *h*N12 at HisH, thus pointing out communication in both directions between substrate and effector sites. The communication between both active sites evolves over time through multiple dynamic pathways. The td-SPM analysis on the activation aMD simulation captures several residues involved in millisecond motions in the ternary complex.(25) Additional complementary insights are gained by tracing the changes in the dynamic network of interactions in the HisF and HisH subunits (SI Extended Text). This analysis paves the way towards a deeper analysis of the molecular basis of the activation process.

## DISCUSSION

Unravelling the molecular mechanisms of allosteric regulation in enzymes involves the structural characterization of functionally relevant states and also monitoring the time evolution of the dynamic conformational ensemble toward the formation of the ternary complex.(2, 12) Millisecond motions, activated by effector binding and finely tuned by the substrate, trigger the allosteric activation of imidazole glycerol phosphate synthase (IGPS), significantly stimulating glutamine hydrolysis in the HisH subunit.(25, 28, 29) However, the rapid turnover observed in *wt*IGPS and the instability of the effector PRFAR prevented the experimental detection of the allosterically active state in the wild-type enzyme, which still remains elusive.(31) The millisecond allosteric activation of IGPS challenges the computational elucidation of these functionally relevant states and the characterization of the time evolution of the conformational ensemble upon ternary complex formation. In this work, a computational strategy tailored to reconstruct millisecond timescale events was devised to describe, step by step, the graded allosteric activation of IGPS at the molecular level, from the inactive substrate-free form to the active ternary complex. Our results reveal a delicate coupling between effector and substrate binding, as well as with the HisF:HisH interface conformational dynamics, which all together regulate the allosteric activation of IGPS ternary complex. Without using *a priori* information of the IGPS active state, the simulations spontaneously uncovered a closed HisF:HisH interface of IGPS with the HisH active site preorganized with the *h*V51 oxyanion hole properly oriented to stabilize the substrate glutamine in a catalytically productive pose. The computational insights provided in this study tie up the loose ends of many of the existing knowns and unknowns in IGPS function and allosteric regulation mechanism.

We explored the effect of PRFAR binding in the substrate-free conformational ensemble of *wt*IGPS with microsecond conventional molecular dynamics (cMD) and accelerated molecular dynamics (aMD) simulations. Our μs-cMD simulations revealed that a hidden conformation of the *h*49-PGVG oxyanion strand with the *h*V51 oxyanion hole formed (active-OxH) can exist in solution when only PRFAR is bound. Although the simulations started from the inactive IGPS X-ray structure with the *h*V51 unformed, we observed that the effect of PRFAR is two-fold: first, it enhances the flexibility of the oxyanion strand triggered by the disruption of *h*Pro10-*h*V51 hydrogen bond and, second, it alters the oxyanion strand conformational ensemble making accessible the catalytically-relevant active-OxH state. In this active-OxH conformation, the HN backbone of *h*V51 is pointing towards the catalytic *h*C84, thus providing a HisH active site properly pre-organized to stabilize the tetrahedral intermediate formed in the glutaminase reaction. This is consistent with the hypothesis that active-OxH can assemble in IGPS as a result of allosteric activation by PRFAR.(22, 25, 26) Our computational insights bridge the structural gap with NMR experiments that indicated the broadening beyond detection of *h*G50 and *h*G52 NH signals upon PRFAR stimulation, suggesting the activation of μs-ms motions in the HisH active site.(25, 28) The active-OxH conformation is infrequently and transiently populated in these μs-cMD simulations. The scarce population of the active-OxH state together with the instability of PRFAR can contribute to explain why the crystallization of wild-type IGPS with PRFAR bound and the oxyanion hole formed remains elusive.(19, 22, 25, 27)

The study of μs-ms motions of IGPS in the substrate-free form with aMD simulations show multiple *h*V51 oxyanion hole formations and also different degrees of closure of the HisF:HisH interface, including the identification of a metastable closed state of IGPS. This conformation, characterized by the alignment of *h*α1 and *f*α3 helices, is transiently populated in the substrate-free simulations and resembles the substrate-bound X-ray closed state of the catalytically-inactive *h*C84A IGPS.(31) Our simulations indicate that a closed state of the HisF:HisH interface can be attained in solution, even in the absence of substrate or PRFAR. These results are consistent with the idea that productive HisF:HisH closure is key for efficient catalysis and for retaining the substrate during glutamine hydrolysis and preventing the loss of the produced ammonia to the media.(46, 48) In line with these observations, Kneuttinger et al. related the partial closure of IGPS with an increase in catalytic activity by introducing a light-switchable non-natural amino acid at position *h*W123.(45)

The reconstruction of the PRFAR-IGPS conformational ensemble in the absence of L-Gln substrate revealed that the *h*V51 oxyanion hole formation and the closed HisF:HisH interface states are transiently populated. Both events occur infrequently and are uncoupled from each other in line with independent μs-ms motions identified with NMR in the presence of PRFAR.(25) At this point, the molecular basis of L-Gln binding into the HisH active site and the subsequent time of evolution toward the formation of the allosterically active ternary IGPS complex were still missing. The reconstruction of the spontaneous binding of glutamine with unconstrained enhanced sampling aMD simulations showed L-Gln binding to the HisH active site in both PRFAR-free and PRFAR-bound IGPS. In both cases, similar binding pathways are followed: initial recognition by *f*Q123, L-Gln captured in the active site when the oxyanion strand attains the inactive-OxH state and the HisF:HisH interface is open, and ultimately stabilization of L-Gln in the HisH active site by the HN *h*G52 and the side chains of a number of HisH and HisF residues. These results are in line with IGPS being predominantly a V-type enzyme, i.e. the substrate affinity is similar in PRFAR-free and PRFAR bound.(19) It should be emphasized that aMD simulations suggest that the inactive-OxH state of the oxyanion strand, is a prerequisite for substrate binding to the HisH active site. When the *h*V51 oxyanion hole is formed, the binding of L-Gln is impeded by the side chain of *h*L85, which lies between *h*V51 and *h*C84. These results provide the molecular explanation to recent NMR studies with the inactivating *h*C84S IGPS mutant that confirm that substrate binding only occurs when IGPS attains the inactive state.(31) Our findings suggest that in the presence of PRFAR, access to both the open state of the HisF:HisH interface and the inactive-OxH state of the oxyanion strand is a pre-requisite for facilitating the recognition and the accommodation of the substrate in the HisH active site.

The binding of glutamine in the HisH active site, in tight coupling with PRFAR, gates a sequence of conformational rearrangements that unravel the time-evolution toward the allosterically active state of IGPS. Extensive μs-aMD simulations indicated that the coupled effect of substrate and PRFAR binding significantly perturbs both the dynamism of *h*49-PGVG oxyanion strand and the HisF:HisH interface conformational ensemble. After L-Gln is captured in the inactive-OxH conformation of the oxyanion strand, the HisF:HisH interface attains the productively closed state. Therefore, substrate binding shifts the conformational ensemble toward closed states of IGPS. In tight coupling with the interdomain closure, multiple long-lived formations of the *h*V51 oxyanion hole are observed along the aMD simulations suggesting similar relative stabilities of active-OxH and inactive-OxH states and low interconversion barriers between them. Indeed, well-tempered metadynamics simulations indicate that the presence of PRFAR decreases the energy barrier of the oxyanion strand rotation (8 kcal/mol) while keeping the equilibrium between inactive-OxH and active-OxH populations unaltered. This is in line with NMR experiments of Lisi et al. that upon PRFAR binding observed a broadening of the ensemble of IGPS conformations without significant changes on the average solution conformations.(28) It also fits with their mutagenesis and NMR experiments indicating that glutamine hydrolysis and associated chemical steps are rate limiting in the presence of PRFAR.(28)

The formation of active-OxH state is coupled to a reorientation of the substrate providing a nucleophilic attack catalytic distance of around 3.4 Å, as opposed to the ca. 4.5 Å in the inactive-OxH conformation. In the active-OxH state, L-Gln is therefore finally properly oriented for proton abstraction and subsequent stabilization of the tetrahedral intermediate. This change in the pre-organization of the HisH active site and the closure of the interdomain HisF:HisH region can be key to trigger the catalytic activity. In line with previous hypotheses based on NMR and kinetic experiments,(25, 26, 46) we suggest this conformation as the allosterically active state of wild type IGPS. Our aMD simulations also point out that both the active-OxH and inactive-OxH states are important for IGPS function, thus, the facile interconversion between both states may be required for the different steps along the catalytic cycle. The fast-dynamic equilibrium between active-OxH and inactive-OxH could be the reason why the active state was not detected in NMR experiments of wild-type IGPS. In the PRFAR-free state, the oxyanion strand interconversion barrier is three-fold higher (i. e. with a barrier of 22 kcal/mol, as compared to 8 kcal/mol in the presence of PRFAR), while keeping the equilibrium population of both oxyanion strand conformations equivalent. Although we have not quantified the corresponding activation barrier for glutamine hydrolysis, the rather high energy barrier associated to the conformational rearrangement of the HisH active site suggests that conformational change is practically unattainable in PRFAR-free IGPS at room temperature. This is indeed consistent with the observed 4500-fold enhancement of basal glutaminase activity of IGPS in the presence of PRFAR.(18) It should be also noted that based on solution NMR experiments on some IGPS mutants a direct correlation between the population of the active conformation and the activity of the IGPS complex (in terms of *k*cat) was proposed.(31) Our results strictly focused on the wild-type IGPS enzyme, however, indicate that PRFAR binding does not alter the relative populations of the active-OxH and inactive-OxH states, but instead impacts the associated active-to-inactive conformational transition barrier.

The analysis of the allosteric communications pathways carried out in this work indicated that the communication between both active sites evolves through multiple dynamic pathways as allosteric activation progresses. The time dependent shortest-path map (td-SPM) analysis revealed the existence of concerted motions that are activated upon productive HisF:HisH interface closure and expand throughout the whole HisF subunit, the interdomain region and HisH active site, resulting in the V51 oxyanion hole formation. Of special interest is the td-SPM analysis along the aMD simulation that captured the complete allosteric activation, which successfully identified several residues involved in millisecond motions in the ternary complex.(25, 29) This contrasts with previous studies based on the application of correlation-based tools focused on short nanosecond timescale MD simulations unable to capture the allosteric activation process.(26) The coupled HisF:HisH interdomain closure with OxH-V51 formation, and the activation of correlated motions preceding allosteric activation are in line with concerted μs-ms motions identified with NMR in the ternary complex.(25) The existence of multiple communication pathways and the activation of millisecond motions are landmark features of dynamic allostery.(8) Based on these multiple communication pathways in the td-SPM analysis and the observed broadening of the conformational ensemble of IGPS in presence of PRFAR, we suggest that IGPS allosteric activation resembles the violin model of dynamics-based allostery suggested in protein kinases.(49, 50)

Through the development of a computational strategy tailored to reconstruct millisecond timescale events, we captured the essential molecular details of the time evolution of the millisecond allosteric activation of IGPS in the ternary complex. Based on these extensive conformational sampling simulations, we suggest a general scheme for describing the IGPS allosteric activation pathway taking place prior to the chemical step (Figure 6). First, oxyanion hole formation and closure of the HisF:HisH interface pre-exist in solution in the substrate free-form, although both are high energy states in the IGPS-PRFAR conformational ensemble. Second, substrate recognition occurs in the IGPS open HisF:HisH interface state, while the oxyanion strand attains an inactive-OxH conformation. Third, the interdomain region productively closes to retain the substrate in the HisH active site. Finally, formation of the *h*V51 oxyanion hole couples with the repositioning of the substrate in a catalytically productive pose to finally form the allosterically active state. The formation of this allosterically active state is controlled by fine-tuned correlated motions connecting the PRFAR effector and HisH binding sites that are activated throughout the whole process.

The proposed model of the allosteric activation pathway of IGPS based on the tailored millisecond timescale computational strategy developed provides multiple new molecular insights not previously identified by means of X-ray crystallography, solution NMR experiments, and short timescale MD simulations. Most importantly, it also answers many of the open questions existing in IGPS allosteric regulation and function. Our computational strategy can be generalized to decipher the molecular basis of allosteric mechanisms in enzyme catalysis, signal transduction, and disease, which is key for developing new therapeutics and engineering novel enzymatic functions.

## MATERIALS AND METHODS

Extended details of all simulations protocols including system set up and simulation analysis can be found in SI *Methods*. Section A of SI *Methods* describes the details of conventional molecular dynamics simulations (cMD) of PRFAR-free and PRFAR bound states. All cMD simulations were performed starting from the IGPS inactive oxyanion strand conformation (PDB: 1GPW). Section B of SI *Methods* provides information about the calibration of parameters for accelerated molecular dynamics simulations (aMD) in PRFAR-free and PRFAR bound states. Section C of SI *Methods* describes the details of the strategy used to study spontaneous glutamine binding and subsequent allosteric activation in the ternary complex with aMD simulations. The starting structures of IGPS for substrate binding simulations were obtained from the most representative conformations of the oxyanion strand sampled in cMD simulations. Section D of SI *Methods* describes the details of well-tempered metadynamics (WT-MetaD) simulations used to estimate the conformational barrier associated to the oxyanion hole formation in PRFAR-free and PRFAR-bound states. Ten representative conformations that encompass global and local features of allosterically inactive-OxH and active-OxH states extracted from aMD simulations were used as starting point for WT-MetaD simulations. Section E of SI *Methods* provides the information on the calculation of dynamical-networks using the shortest-path Map (SPM) tool. SPM is calculated in time spans of 600 ns along the aMD trajectory that describes the complete allosteric activation of IGPS considering all alpha carbons of the protein.

## Supporting information

S1 Movie - cMD IGPS-PRFAR

S2 Movie - aMD IGPS-PRFAR L-Gln binding

S3 Movie - aMD IGPS-PRFAR L-Gln binding (global view)

S4 Movie - aMD ternary complex - allosteric activation

## DATA AVAILABILITY

Relevant structures of key functional states and molecular dynamics trajectories are available at https://github.com/ccalvotusell/igps

## ACKNOWLEDGEMENTS

We thank the Generalitat de Catalunya for the emerging group CompBioLab (2017 SGR-1707) and Spanish MINECO for projects PGC2018-102192-B-I00 (S.O) and RTI2018-101032-J100 (F.F). S.O. is grateful to the funding from the European Research Council (ERC) under the European Union’s Horizon 2020 research and innovation program (ERC-2015-StG-679001). F.F. thanks the Spanish Super-computing Network (RES) for access to supercomputing resources (project BCV-2021-1-0015). M. A. M. S. was supported by the Spanish MINECO for a PhD fellowship (BES-2015-074964) and by the National Research Foundation of Korea (NRF) under the Brain Pool Program (NRF-2021H1D3A2A02038434).

## Notes

https://github.com/ccalvotusell/igps

### Competing Interest Statement

The authors have declared no competing interest.

## REFERENCES

1. S. J. Wodak, et al., Allostery in Its Many Disguises: From Theory to Applications. Structure 27, 566–578 (2019).

2. C. J. Tsai, R. Nussinov, A Unified View of “How Allostery Works.” PLoS Comput. Biol. 10, e1003394 (2014).

3. G. P. Lisi, J. P. Loria, Allostery in enzyme catalysis. Curr. Opin. Struct. Biol. 47, 123–130 (2017).

4. J. Monod, J. Wyman, J. P. Changeux, On the nature of allosteric transitions: A plausible model. J. Mol. Biol. 12, 88–118 (1965).

5. A. Cooper, D. T. F. Dryden, Allostery without conformational change - A plausible model Eur. Biophys. J. 11, 103–109 (1984).

6. R. Nussinov, C. J. Tsai, Allostery without a conformational change? Revisiting the paradigm. Curr. Opin. Struct. Biol. 30, 17–24 (2015).

7. G. Jiménez-Osés, et al., The role of distant mutations and allosteric regulation on LovD active site dynamics. Nat. Chem. Biol. 10, 431–436 (2014).

8. J. Guo, H. X. Zhou, Protein Allostery and Conformational Dynamics. Chem. Rev. 116, 6503–6515 (2016).

9. A. W. Fenton, Allostery: an illustrated definition for the “second secret of life.” Trends Biochem. Sci. 33, 420–425 (2008).

10. N. V. Dokholyan, Controlling Allosteric Networks in Proteins. Chem. Rev. 116, 6463–6487 (2016).

11. O. Bozovic, et al., Real-time observation of ligand-induced allosteric transitions in a PDZ domain. Proc. Natl. Acad. Sci. U. S. A. 117, 26031–26039 (2020).

12. P. Mehrabi, et al., Time-resolved crystallography reveals allosteric communication aligned with molecular breathing. Science (80-.). 365, 1167–1170 (2019).

13. J. S. Fraser, et al., Hidden alternative structures of proline isomerase essential for catalysis. Nature 462, 669–673 (2009).

14. H. Y. Aviram, et al., Direct observation of ultrafast large-scale dynamics of an enzyme under turnover conditions. Proc. Natl. Acad. Sci. U. S. A. 115, 3243–3248 (2018).

15. M. Kovermann, C. Grundström, A. Elisabeth Sauer-Eriksson, U. H. Sauer, M. Wolf-Watz, Structural basis for ligand binding to an enzyme by a conformational selection pathway. Proc. Natl. Acad. Sci. U. S. A. 114, 6298–6303 (2017).

16. M. Dasgupta, et al., Mix-and-inject XFEL crystallography reveals gated conformational dynamics during enzyme catalysis. Proc. Natl. Acad. Sci. U. S. A. 116, 25634–25640 (2019).

17. K. W. East, et al., Allosteric Motions of the CRISPR–Cas9 HNH Nuclease Probed by NMR and Molecular Dynamics. J. Am. Chem. Soc. 142, 1348–1358 (2019).

18. S. Beismann-Driemeyer, R. Sterner, Imidazole glycerol phosphate synthase from Thermotoga maritima. Quaternary structure, steady-state kinetics, and reaction mechanism of the bienzyme complex. J. Biol. Chem. 276, 20387–20396 (2001).

19. B. N. Chaudhuri, S. C. Lange, R. S. Myers, V. J. Davisson, J. L. Smith, Toward understanding the mechanism of the complex cyclization reaction catalyzed by imidazole glycerolphosphate synthase: Crystal structures of a ternary complex and the free enzyme. Biochemistry 42, 7003–7012 (2003).

20. I. Rivalta, et al., Allosteric Communication Disrupted by a Small Molecule Binding to the Imidazole Glycerol Phosphate Synthase Protein-Protein Interface. Biochemistry 55, 6484– 6494 (2016).

21. G. P. Lisi, A. A. Currier, J. P. Loria, Glutamine Hydrolysis by Imidazole Glycerol Phosphate Synthase Displays Temperature Dependent Allosteric Activation. Front. Mol. Biosci. 5, 4 (2018).

22. B. N. Chaudhuri, et al., Crystal structure of imidazole glycerol phosphate synthase: A tunnel through a (β/α)8 barrel joins two active sites. Structure 9, 987–997 (2001).

23. R. E. Amaro, R. S. Myers, V. J. Davisson, Z. A. Luthey-Schulten, Structural elements in IGP synthase exclude water to optimize ammonia transfer. Biophys. J. 89, 475–487 (2005).

24. T. J. Klem, V. J. Davisson, Imidazole Glycerol Phosphate Synthase: The Glutamine Amidotransferase in Histidine Biosynthesis. Biochemistry 32, 5177–5186 (1993).

25. J. M. Lipchock, J. P. Loria, Nanometer propagation of millisecond motions in V-type allostery. Structure 18, 1596–1607 (2010).

26. I. Rivalta, et al., Allosteric pathways in imidazole glycerol phosphate synthase. Proc. Natl. Acad. Sci. U. S. A. 109, 8366 (2012).

27. F. List, et al., Catalysis uncoupling in a glutamine amidotransferase bienzyme by unblocking the glutaminase active site. Chem. Biol. 19, 1589–1599 (2012).

28. G. P. P. Lisi, et al., Dissecting Dynamic Allosteric Pathways Using Chemically Related Small-Molecule Activators. Structure 24, 1155–1166 (2016).

29. G. P. Lisi, K. W. East, V. S. Batista, J. P. Loria, Altering the allosteric pathway in IGPS suppresses millisecond motions and catalytic activity. Proc. Natl. Acad. Sci. U. S. A. 114, E3414–E3423 (2017).

30. J. B. Thoden, et al., Carbamoyl phosphate synthetase: Caught in the act of glutamine hydrolysis. Biochemistry 37, 8825–8831 (1998).

31. J. P. Wurm, et al., Molecular basis for the allosteric activation mechanism of the heterodimeric imidazole glycerol phosphate synthase complex. Nat. Commun. 2021 121 12, 1–13 (2021).

32. A. T. Vanwart, J. Eargle, Z. Luthey-Schulten, R. E. Amaro, Exploring residue component contributions to dynamical network models of allostery. J. Chem. Theory Comput. 8, 2949–2961 (2012).

33. A. A. S. T. Ribeiro, V. Ortiz, Determination of signaling pathways in proteins through network theory: Importance of the topology. J. Chem. Theory Comput. 10, 1762–1769 (2014).

34. A. T. Van Wart, J. Durrant, L. Votapka, R. E. Amaro, Weighted implementation of suboptimal paths (WISP): An optimized algorithm and tool for dynamical network analysis. J. Chem. Theory Comput. 10, 511–517 (2014).

35. C. F. A. Negre, et al., Eigenvector centrality for characterization of protein allosteric pathways. Proc. Natl. Acad. Sci. U. S. A. 115, E12201–E12208 (2018).

36. W. M. Botello-Smith, Y. Luo, Robust Determination of Protein Allosteric Signaling Pathways. J. Chem. Theory Comput. 15, 2116–2126 (2019).

37. A. Gheeraert, et al., Exploring Allosteric Pathways of a V-Type Enzyme with Dynamical Perturbation Networks. J. Phys. Chem. B 123, 3452–3461 (2019).

38. P. T. Lake, R. B. Davidson, H. Klem, G. M. Hocky, M. McCullagh, Residue-Level Allostery Propagates through the Effective Coarse-Grained Hessian. J. Chem. Theory Comput. 16, 3385–3395 (2020).

39. P. Campitelli, T. Modi, S. Kumar, S. B. Ozkan, The Role of Conformational Dynamics and Allostery in Modulating Protein Evolution. https://doi.org/10.1146/annurev-biophys-052118-115517 49, 267–288 (2020).

40. J. R. Wagner, et al., Emerging Computational Methods for the Rational Discovery of Allosteric Drugs. Chem. Rev. 116, 6370–6390 (2016).

41. S. Mouilleron, B. Golinelli-Pimpaneau, Conformational changes in ammonia-channeling glutamine amidotransferases. Curr. Opin. Struct. Biol. 17, 653–664 (2007).

42. D. Hamelberg, J. Mongan, J. A. McCammon, Accelerated molecular dynamics: A promising and efficient simulation method for biomolecules. J. Chem. Phys. 120, 11919– 11929 (2004).

43. D. Hamelberg, C. A. F. De Oliveira, J. A. McCammon, Sampling of slow diffusive conformational transitions with accelerated molecular dynamics. J. Chem. Phys. 127, 155102 (2007).

44. Y. Miao, J. A. McCammon, Graded activation and free energy landscapes of a muscarinic G-protein–coupled receptor. Proc. Natl. Acad. Sci. 113, 12162–12167 (2016).

45. A. C. Kneuttinger, et al., Significance of the Protein Interface Configuration for Allostery in Imidazole Glycerol Phosphate Synthase. Biochemistry 59, 2729–2742 (2020).

46. R. S. Myers, R. E. Amaro, Z. A. Luthey-Schulten, V. J. Davisson, Reaction coupling through interdomain contacts in imidazole glycerol phosphate synthase. Biochemistry 44, 11974–11985 (2005).

47. A. Romero-Rivera, M. Garcia-Borràs, S. Osuna, Role of Conformational Dynamics in the Evolution of Retro-Aldolase Activity. ACS Catal. 7, 8524–8532 (2017).

48. A. Douangamath, et al., Structural evidence for ammonia tunneling across the (βα)8 barrel of the imidazole glycerol phosphate synthase bienzyme complex. Structure 10, 185–193 (2002).

49. L. G. Ahuja, S. S. Taylor, A. P. Kornev, Tuning the “violin” of protein kinases: The role of dynamics-based allostery. IUBMB Life 71, 685–696 (2019).

50. L. G. Ahuja, A. P. Kornev, C. L. Mcclendon, G. Veglia, S. S. Taylor, Mutation of a kinase allosteric node uncouples dynamics linked to phosphotransfer. Proc. Natl. Acad. Sci. U. S. A. 114, E931–E940 (2017).

