## Supplementary Information for "Unravelling the Graded Millisecond Allosteric Activation Mechanism of Imidazole Glycerol Phosphate Synthase"

### Table of contents

1. SI Methods
2. SI Extended Text
3. Conventional Molecular Dynamics Simulations IGPS: substrate-free (Figures S2-S10)
4. Accelerated Molecular Dynamics Simulations IGPS: substrate-free (Figures S11-S14)
5. Accelerated Molecular Dynamics Simulations IGPS: spontaneous substrate binding (Figures S15-S22)
6. Accelerated Molecular Dynamics Simulations IGPS: ternary complex (Figures S23-S26 and S28-S29)
7. Metadynamics Simulations IGPS: ternary complex (Figure S27)
8. Dynamical-network analysis IGPS (Shortest-Path Map): ternary complex (Figures S30-S31)

### *1. SI Methods.*

#### Computational Strategy

In this work, we use molecular dynamics (MD) simulations, enhanced sampling techniques, and dynamical networks to characterize the molecular details of the millisecond graded allosteric activation of IGPS and identify hidden states relevant for IGPS catalytic activity. IGPS is prepared for MD simulations in both the absence (apo) and presence of PRFAR (PRFAR-IGPS). The starting point for the computational sampling is an inactive IGPS conformation (PDB:1GPW), that is, the *h*49-PGVG oxyanion strand is found in an inactive conformation (the $H^N$ *h*V51 is not pointing toward HisH active site) and the HisF:HisH is found in a partially open state (HisF:HisH interface angle around 25º). From this initial structure, the following computational strategy is applied to characterize the allosterically active state of IGPS.

First, we explored the effect of PRFAR binding on the oxyanion strand conformational dynamics through long-time scale conventional MD (cMD) simulations to characterize μs time-scale motions. These cMD simulations provided the conformational ensemble of the oxyanion strand in both, the absence and presence of PRFAR. Second, to capture the millisecond motions characteristic of IGPS allosteric activation, we resorted to accelerated molecular dynamics (aMD). These aMD simulations provided information of both, the oxyanion strand dynamics and global IGPS dynamics beyond the microsecond time scale. Third, the most relevant conformational states of the oxyanion strand sampled in cMD simulations were used as a starting point for spontaneous substrate (L-Gln) binding aMD simulations. From these simulations, we explored the spontaneous formation of the ternary complex initiated by the substrate binding process and subsequent allosteric activation. aMD simulations reveal the spontaneous binding of glutamine in the HisH active site and the formation of the *h*V51 oxyanion hole in the ternary complex, pointing out a tight coupling between interdomain changes, substrate binding, and oxyanion hole formation. Fourth, the states identified with aMD are used as starting points for well-tempered metadynamics simulations to reconstruct the free energy landscape of the oxyanion strand conformational dynamics. Finally, we explore the existence of correlated motions with the shortest-path map tool along the allosteric activation process that highlight an enhancement of communication between the two subunits upon activation. This computational strategy can be generalized to decipher allosteric mechanisms and identify hidden states along the allosteric activation pathway. The different steps of the computational protocol are summarized in Fig. S1.

**System preparation**

*Protein preparation.* The computational structural models of IGPS were based on the crystal structure of the *apo* complex from *Thermotoga maritima* at 2.4 Å resolution (PDB:1GPW) reported by Douangamath and coworkers.(1) To generate the structural model of IGPS, chains A and B of PDB 1GPW were used. In chain B of 1GPW, the *h*49-PGVG oxyanion strand is found in an inactive conformation (Inactive-OxH). It is postulated that the C-terminal *loop* of chains A and B is found in a closed (assumed active) conformation. The original bacterial crystal structure presents an active site mutation (*f*D11N) that is mutated back to its original residue using PyMOL. The crystal structure of the PRFAR-bound complex from *Saccharomyces cerevisiae* at 2.5 Å resolution (PDB:1OX5), crystallized with the effector PRFAR, was used to generate the PRFAR-bound state. The coordinates of the effector PRFAR were aligned to two phosphate groups from the chains A and B of the PDB 1GPW. These phosphate groups were suggested to belong to an unresolved PRFAR molecule since the effector was present in the solution but not in the crystal during the crystallization procedure. Following the same system preparation described in previous works, the crystallographic waters of 1GPW were kept for the molecular dynamics simulations.(2) According to previous works,(2–4) a δ-nitrogen (HID) protonation state was assigned to residues *f*H84, *f*H209, *f*H244 of HisF subunit and *h*H73, *h*H120, *h*H141, and *h*H178 in HisH subunit; ε-nitrogen (HIE) protonation state for residues *f*H228 and *h*H53 and both δ-nitrogen and ε-nitrogen (HIP) of residue *f*H151 were protonated. The catalytic residues *h*C84, *h*H178, and *h*E180 were treated as protonated thiol group (-SH), δ-nitrogen protonated (HID), and deprotonated carboxylate group ($COO^-$).

*Ligand parametrization.* PRFAR initial structure has been obtained from PDB 1OX5. Parameters for MD simulations for PRFAR and L-Gln were generated with antechamber module of AMBER16(5) using the generalized AMBER force field (GAFF),(6) with partial charges set to fit the electrostatic potential generated at HF/6-31G* level of theory by restrained electrostatic potential (RESP) model.(7) The atomic charges were calculated according to the Merz−Singh−Kollman(7) scheme using Gaussian 09.(8)

**Conventional Molecular Dynamics simulation protocols**

Molecular Dynamics simulations of all IGPS complexes (apo and PRFAR effector-bound) were performed in explicit water using AMBER16 package. AMBER-ff14SB force field(9) was used to describe the protein, GAFF for PRFAR and L-Gln and TIP3P for water molecules.(10) Each system was solvated in a pre-equilibrated cubic box with a 10 Å buffer of TIP3P water molecules and was neutralized by addition of explicit sodium and chloride counterions ($Na^+$ or $Cl^-$). Subsequently, a two-stage geometry optimization approach was performed. First, a short minimization of the water molecules positions, with positional restraints on solute by a harmonic potential with a force constant of 500 kcal $mol^{-1}$ $Å^{-2}$ was done. The second stage was an unrestrained minimization of all the atoms in the simulation cell. Then, the systems were gently heated using six 50 ps steps, incrementing the temperature 50 K each step (0-300 K) under constant-volume, periodic-boundary conditions and the particle-mesh Ewald approach to introduce long-range electrostatic effects.(11) For these steps, an 11 Å cut-off was applied to Lennard-Jones and electrostatic interactions. Bonds involving hydrogen were constrained with the SHAKE algorithm. Harmonic restraints of 10 kcal $mol^{-1}$ were applied to the solute, and the Langevin equilibration scheme is used to control and equalize the temperature. The time step was kept at 2 fs during the heating stages, allowing potential inhomogeneities to self-adjust. Each system was then equilibrated for 4 ns with a 2 fs timestep at a constant pressure of 1 atm to relax the density of the system. After the systems were equilibrated in the NPT ensemble, MD simulations were performed under the NVT ensemble and periodic-boundary conditions using the Galatea cluster of the university of Girona (composed by 178 GTX1080 GPUs). PRFAR-bound state simulations were carried out applying distance restrains between the effector phosphate groups and two residues (*f*T104/*f*A224) of PRFAR binding site. The cMD simulations data used for the following analyses consist of 10 replicas of 1.5 μs for the *apo* state; and 10 replicas of 1.5 μs for the PRFAR-IGPS state. The analysis of the distances, angles, and dihedral angles is performed using the *cpptraj* MD analysis MD program. The PyEMMA 2.5 software was used for constructing the conformational landscape and for clustering analysis.

**Accelerated Molecular Dynamics simulation protocols**

Accelerated Molecular Dynamics simulations (aMD) have been used to explore the conformational dynamics of *apo* and PRFAR-IGPS.(12, 13) Starting from the inactive IGPS structures (see above), we performed unrestrained conventional MD simulations (100 ns) as described above from which the acceleration parameters were determined. Then, ten replicas of 1 μs of dual-boost accelerated Molecular Dynamics (aMD) simulations were carried out in both apo and PRFAR-IGPS states.

aMD enhances the conformational sampling of biomolecules, by adding a non-negative boost potential to the system when the system potential is lower than a reference energy:

$$V^*(r) = V(r), \qquad V(r) \geq E,$$
$$V^*(r) = V(r) + \Delta V(r), \quad V(r) < E,$$

where $V(r)$ is the original potential, *E* is the reference energy, and $V^*(r)$ is the modified potential. In the simplest form, the boost potential, $\Delta V(r)$ is given by:

$$\Delta V(r) = \frac{(E - V(r))^2}{\alpha + E - V(r)}, \qquad (2)$$

where $\alpha$ is the acceleration factor. As the acceleration factor $\alpha$ decreases, the energy surface is flattened more and biomolecular transitions between the low-energy states are increased.

Here a total boost potential is applied to all atoms in the system in addition to a more aggressive dihedral boost, *i.e.*, ($E_{\text{dihed}}$, $\alpha_{\text{dihed}}$; $E_{\text{total}}$, $\alpha_{\text{total}}$), within the dual-boost aMD approach. The acceleration parameters used in this work for exploring IGPS conformational dynamics, are the following:

$$E_{\text{dihed}} = V_{\text{dihed\_avg}} + a_1 \times N_{\text{res}}, \ \alpha_{\text{dihed}} = a_2 \times N_{\text{res}}/5;$$
$$E_{\text{total}} = V_{\text{total\_avg}} + b_1 \times N_{\text{atoms}}, \ \alpha_{\text{total}} = b_2 \times N_{\text{atoms}}, \qquad (3)$$

where $N_{\text{res}}$ is the number of protein residues, $N_{\text{atoms}}$ is the total number of atoms, and $V_{\text{dihed\_avg}}$ and $V_{\text{total\_avg}}$ are the average dihedral and total potential energies calculated from 100 ns cMD simulations, respectively. aMD simulations were performed for each system after 100 ns of cMD. Two different levels of acceleration were tested for *apo* and PRFAR-IGPS. The parameters used for aMD simulations are: High acceleration ($a_1$=3.5, $a_2$=3.5; $b_1$=0.175, $b_2$=0.175) and moderate accelerated ($a_1$=2, $a_2$=3.5; $b_1$=0.16, $b_2$=0.16). The high acceleration parameters were selected for exploring the conformational dynamics of IGPS in both apo and PRFAR-bound states. The moderate accelerated parameters were selected for exploring the spontaneous substrate binding process of L-Gln and subsequent allosteric activation of IGPS.

*Substrate Binding Simulations*. Accelerated Molecular Dynamics simulations (aMD) have been used to study the spontaneous binding of glutamine (L-Gln) in the HisH active site of both *apo* and PRFAR states. In both cases, we placed one substrate in the solvent with a minimum distance of 25 Å from the *h*C84 active site residue. As starting IGPS structures, we selected the most representative structures of the oxyanion strand conformational ensemble sampled in the cMD simulations. In *apo*-IGPS, 15 replicas of 600 ns of aMD were run from the each Inactive-OxH and Unblocked-OxH states. In PRFAR-IGPS, 10 replicas of 600 ns of aMD were run from each Inactive-OxH, Unblocked-OxH, and Active-OxH conformations. From these coordinates, we first performed unrestrained conventional MD simulations (100 ns) from which the acceleration parameters were determined. Then, a total of 30 replicas of 600 ns of dual-boost accelerated Molecular Dynamics (aMD) for the *apo* and PRFAR-IGPS simulations were performed to allow the substrate to diffuse freely until it spontaneously associates with the surface of the protein, and finally targets the active site. The simulations that substrate binding was observed were extended up to 5 microseconds (one to 10 microseconds) using the moderated acceleration parameters following Equation 3. These long timescale unconstrained aMD simulations were performed with the aim of capturing the complete allosteric activation of IGPS.

**Well-tempered Metadynamics Simulations**

The PLUMED2(14) software package together with the GROMACS 5.1.2 code (M.J. Abraham et al. GROMACS User Manual version 5.1.2) were used to carry out the metadynamics simulations. Metadynamics enhances the sampling of the conformational space by adding external energy potentials to a selected set of degrees of freedom, namely collective variables (CVs). The CVs chosen were the φ dihedral angle of *h*G50 and the φ dihedral angle of *h*V51.This bias potential gradually overcomes energy barriers allowing for efficient exploration of different conformational states. After a certain simulation time, the biasing potential corresponds to the negative of the free energy surface (FES) and, therefore, all possible states are equally sampled. More exhaustive discussions of the method can be found elsewhere. Here, the well-tempered version of metadynamics algorithm was used to improve the convergence of the FES reconstruction.(15) Initial Gaussian potentials of height 0.5 kcal $mol^{-1}$, deposited every 2 ps of MD simulations at 300 K, were gradually decreased over time proportional to the potential deposited in the currently visited point of the CV space. A bias factor parameter of 10 was selected to control how quick the Gaussian height is decreased.

The WT-Metadynamics simulations were performed in combination with multiple-walkers approach.(16) We used 10 conformations sampled in aMD simulations as starting points for the WT-metadynamics simulations. In this case, we started five walker replicas in the active and five walker replicas in the inactive states in order to increase the sampling of the oxyanion hole conformational space. The ten walker replicas were run in parallel reading the external energy potentials deposited by the others, thus reconstructing the same metadynamics bias simultaneously. Each walker replica was run for 20 ns, giving a total of 200 ns simulation time for each system computed. Finally, the FEL of the oxyanion strand conformational dynamics was completely reconstructed by summing the Gaussian potentials deposited by all walker replicas as a function of the CVs.

**Shortest Path Map analysis.**

The first step of the Shortest Path Map (SPM) calculation relies on the construction of a graph based on the computed mean distances and correlation values observed along the MD simulations. For each residue of the protein a node is created and centered on the Cα atom if both residues display a mean distance less than 6 Å along the simulation time. The length of the line connecting both residues is drawn according to their correlation value ($d_{ij}$=-log $|C_{ij}|$). Larger correlation values (closer to 1 or -1) will have shorter edge distances, whereas less correlated residue pairs (values closer to 0) will have edges with long distances. At this point, we make use of Dijkstra algorithm to identify the shortest path lengths. The algorithm goes through all nodes of the graph and identifies which is the shortest path to go from the first until the last protein residue. The method therefore identifies which are the edges of the graph that are shorter, i.e. more correlated, and that are more frequently used for going through all residues of the protein, i.e. they are more central for the communication pathway. More details about our SPM tool can be found in our recent publication in ACS Catalysis.(17) To capture the changes on the residue-correlations during the allosteric activation, we decided to split the analysis of aMD trajectories in concatenated time spans of 600 ns (i.e. from 0-600 ns, from 300-900 ns, from 600-1200 ns, …) in what we call time-dependent SPM (td-SPM). The td-SPM analysis is performed considering all alpha carbons of all residues of the protein.

### 2. SI Extended Text

#### Analysis of Conventional Molecular Dynamics Simulations.

*Oxyanion strand conformational dynamics.* Additional unblocked-OxH states of the oxyanion strand presenting similar characteristics are sampled in cMD simulations. These conformations are represented by the rotation of other dihedral angles of the oxyanion strand (see Fig. S2 and S3) such as $\phi$ hG52. The analysis of individual cMD trajectories show that the inactive-OxH and unblocked-OxH states can interconvert in the microsecond time scale (2 out 10 replicas for $\phi$ hG50, while $\phi$ hG52 transition occurs more frequently). The conformation of the unblocked state associated with the rotation of $\phi$ hG52 resembles the one identified by Kneuttinger and coworkers by means of MD simulations.(18)

*Loop1 conformational dynamics and hydrophobic cluster*. To assess the changes in global backbone flexibility, the Root-mean-square fluctuations (RMSF) of all Cα atoms is computed for *apo* and PRFAR bound states. RMSF analysis show no signficant global backbone rearrangements, both *apo* and PRFAR display similar patterns (see Fig. S8). The main conformational differences are located in Loop 1 of the HisF subunit (*f*R16-*f*D31), which presents enhanced flexibility in the presence of PRFAR. In the x-ray (PDB 1GPW), Loop1 is formed by two small β-sheets strands that are stabilized by a hydrogen bond network of conserved residues in this enzyme family. Interestingly significant differences arise after one microsecond of simulation time when Loop1 loses its secondary structure and moves away from the cyclase active site (see Fig. S8). Loop1 adopts a conformational structure similar to some IGPS x-ray structures (e.g. chain E of PDB 1GPW) where the loop is disordered and partially unsolved. This conformational transition was not described previously and occurs in three out of ten PRFAR bound replicas indicating that PRFAR binding enhances microsecond Loop 1 motions compared to *apo*. In the PRFAR bound replica where the formation of the oxyanion hole is observed, the conformational change of Loop1 is correlated with the rotation of the oxyanion strand and preceded by the disruption of the hydrophobic cluster (*f*F23 and *f*I52 hydrophobic interaction, see Fig. S9). Upon the conformational change, both *f*F23 and *f*R27 are pointing towards the solvent while *f*K19 establishes transient interactions with the glycerol phosphate group (gP), in contrast to the *apo* state simulations where *f*K19 is exposed to the bulk during the whole simulation time. The hydrophobic cluster remains unformed and flexible when IGPS presents the oxyanion hole formed. The inner flexibility of Loop1 is related to facile proteolysis by trypsin at *f*R27 position.(19) Upon hV51 oxyanion hole formation, Loop 1 remains relatively stable for 1 μs of MD simulation time while subtle changes occur upon oxyanion hole deactivation. However, in other replicas the Loop1 conformational rearrangements are uncoupled to changes on the oxyanion strand dynamics indicating the existence of uncorrelated motions. Our hypothesis is that the ordered x-ray conformation of Loop1 (used as a starting point in most IGPS MD simulations) is not the most stable in solution. These observations corroborate previous NMR and computational studies that described that PRFAR binding alters Loop 1 dynamics.

*Salt Bridge network between fα2, fα3 and hα1 and Heterodimer Interface network*. The binding of PRFAR gated a series of conformational rearrangements on the HisF and HisH subunits besides the formation of the oxyanion hole and motions in Loop1 (see Fig. S9). In particular, the salt bridge network between *f*α2, *f*α3 and *h*α1 helices presents some alterations in the presence of PRFAR. In the early steps of the allosteric activation, the salt-bridge interactions between *f*E67 and both *f*R95 and *h*R18 are strengthened compared to *apo* while the interaction between *f*E71 and *h*R18 is weakened (see Fig. S9). *h*Arg18 can adopt two major conformations, one pointing towards the dimer interface and another pointing towards the solvent. The reshape of these interactions enhances the communication between the *f*α2, *f*α3 and *h*α1 structural motifs and are key to unlocked changes on the interdomain region that precede the oxyanion strand formation. These rearrangements impact orientation of *h*N12 and *h*N15 in *h*α1. In particular, *h*N15 backbone establishes a transient hydrogen bond with the *h*Arg18 side chain that helps orient the side chain of *h*N15 towards the interdomain region and HisH active site. These rearrangements precede the rotation of *h*P10, leads to the breaking of *h*P10-*h*V51 hydrogen bond and the displacement of the Ω-loop (see Fig. S4). The breaking of *h*P10-*h*V51 interaction is, thus, a prerequisite for oxyanion hole formation and for the activation of HisH for catalysis. However, in the *apo* state simulations the hydrogen bond between *h*N15 and *h*Arg18 is not established and hN15 leaves move away from the interdomain region while keeping the Ω-loop stable in the inactive form. Finally, at the same time, the formation of the oxyanion hole alters the interdomain salt bridge between the side chains of *f*D98 and *h*K181 that has been shown to be key for allosteric communication. Upon oxyanion hole formation the salt bridge is weakened compared to *apo* and PRFAR inactive states (see Fig. S9). *h*K181 gains mobility establishing interactions with catalytic *h*E180 when the breathing motion is more compressed while *f*D98 alters between *h*Y138, *h*N15, *h*K181 residues. Therefore, PFRAR alters the electrostatic environment of the interdomain region. However, the sequence of these events differ among replicas indicating the predominance of uncorrelated motions in these non-equilibrium MD simulations.

*HisF:HisH interface.* All these local rearrangements impact the global dynamics of IGPS. One of the unanswered questions is whether IGPS is able to attain a closed state of the HisF:HisH interface to retain ammonia. When analyzed globally, the HisF:HisF interface conformational dynamics is not showing significant differences in the *apo* and PRFAR bound states (see Fig. S10). Angles between 15º and 35º are sampled and similar distributions are observed in both cases. However, displacement towards shorter angles is observed in the replica where the oxyanion hole is formed. A deeper analysis of the MD simulation where the oxyanion hole formation is observed reveals that the HisF:HisH conformational dynamics becomes slightly restrained when the active state is sampled (values below 20 Å are frequently sampled and stabilized). This is consistent with the idea of a population shift towards a closed state of the interdomain region when PRFAR is present. However, the values sampled in the cMD simulations are still far from the values observed in the x-ray *h*C84A IGPS structure.

**Analysis of Accelerated Molecular Dynamics Simulations.**

To unravel the effect of PRFAR on the conformational dynamics of IGPS beyond microsecond time scale, we run ten replicas of 1 μs accelerated molecular dynamics (aMD) simulations in the *apo* and PRFAR bound states. To explore global conformational motions in deeper detail, we performed PCA analysis on the aMD simulations (see Fig. S13). Interestingly, PC1 describes the counter rotation of HisH and HisF subunits involving most of the residues on the sideR of IGPS pointing out an enhanced communication between subunits. On the other hand, PC2 captures motions in Loop1 and changes on the HisF:HisH interface. Overall, a number of metastable states are identified that present different degrees of HisF:HisH rotation and interdomain closure/occlusion. In more detail, PRFAR unlocks the rotation of HisF:HisH subunits compared to *apo*, flexibilizing the interdomain region. These results are in line with the enhanced millisecond motions upon PRFAR binding observed in NMR studies. Several orientations of the HisF:HisH subunits are found to be relatively stable along PC1 that highlight different closures of IGPS interdomain regions. In the most populated one ($\mathbf{P^C}$, see Fig. S13), the oxyanion strand loop interacts with the top of *f*α4 residues (*f*T119) while the Ω-loop and the bottom of *h*α1 establish interactions with the top of *f*α3. In this state, the HisF:HisH interface angle decreases to average values of 14.0 ± 3.5 º. The two additional states along PC1 correspond to different degrees of rotation of the HisF:HisH subunits. In $\mathbf{R^L}$, the Ω-loop interacts with the top of *f*α4 residues and represents the displacement of HisH towards the left with respect to HisF. In **R**$^{R}$, the HisH subunit rotates towards the right with respect to HisF, with the oxyanion strand residues topping the *f*α3. This degree of rotation is not captured in *apo* IGPS. PC2 captures transitions in Loop 1 and the closure of the HisF:HisH interface. We identified a state that displays productive closure (**C**, HisF:HisH interface angle of 11.3 ± 0.6 º), as observed in the x-ray *h*C84A IGPS structure, PDB 7AC8 (interface angle of 9.7 º). In this state, the amide backbone of *h*H53 establishes a hydrogen bond with the carbonyl backbone of *f*T119, the Ω-loop collapses over *f*α3 and the *h*α1 and *f*α3 are perfectly aligned. We have identified a potential productively closed state of IGPS that can be key for catalytic activity. In general, the conformational ensemble is displaced towards shorter angles of the HisF:HisH interface (see Fig. S13e). However the closure of the subunit is not stabilized through the simulation time and is not correlated with other motions. Similar closed states are sampled in the *apo* state simulations indicating that the efficient closure of IGPS is not limited to PRFAR simulations. PRFAR releases tension in the interdomain region facilitating the rotation of HisH and HisF subunits and the closure of the interdomain region.

**Analysis of WT-Metadynamics Simulations**

To estimate the energy barrier of the oxyanion strand reorientation, we performed well-tempered metadynamics (WT-Metadynamics) simulations using the multiple-walkers approach. Defining a reaction coordinate without knowing the end state can be difficult. However, the inactive and active states of IGPS identified in the aMD simulations can be used as a reference for accurate metadynamics simulations. Therefore, we use the aMD simulations to extract relevant conformations to seed the starting conformations (i.e. walker replicas staring points) for metadynamics simulations (see Fig. S27 for more details). The selected states encompass global and local features of inactive and active states respectively. In this case, we started five walkers in the Active-OxH and five walkers in the Inactive-OxH states to completely reconstruct the free energy landscape of the oxyanion strand conformational dynamics. The FEL obtained from metadynamics simulations show remarkable differences in the PRFAR-free and PRFAR bound states. In the PRFAR bound state, the formation of the oxyanion hole presents a surmountable energy barrier of 8 kcal/mol while in the PRFAR-free state this value rises to 22 kcal/mol. These results are in line with experimental $k_{cat}$ values. Further, the relative stability of the inactive and active forms is maintained which indicates that both states are accessible in PRFAR and may be important for enzyme catalysis (binding and chemical step). These transitions take place in the IGPS closed state without changes in the HisF:HisH interface. Most importantly, while in the presence of PRFAR the oxyanion hole can easily arrange and disarrange in the closed state, the oxyanion hole cannot form in the PRFAR-free state hampering the catalytic activity

**Ternary complex HisF Conformational Dynamics**

Additional complementary insights are gained by further tracing the changes in the dynamic network of interactions in the HisF and HisH subunits, including alterations in the open-to-closed exchange of the interdomain region (see Fig. S30 and S31). In the presence of PRFAR, *f*K19 side chain makes permanent contacts with the glycerol phosphate group (gP) of PRFAR (distance of *ca.* 3.87 ± 0.63 Å). This interaction remains formed most of the simulation time and helps orienting PRFAR in the cyclase active site. Among the residues that form the hydrophobic cluster, the interactions between *f*L50-*f*I52 and *f*V48-*f*L50 are persistent along the simulation (see Fig. S30). When IGPS attains the HisF:HisH closed state, a series of dynamic interactions propagate through the two active sites of HisH and HisF and interdomain regions. In particular, we observed in the aMD simulation that the productive closure is preceded by the breaking of *f*E67-*h*R18 salt bridge in the presence of PRFAR. This interaction occurs in the interdomain HisF:HisH region and seems to play a key role in controlling the dynamics of interface residues, which exhibit a higher flexibility in the presence of PRFAR. On the other hand, the *f*E67-*f*R95 salt bridge remains quite formed and stable as well as the *f*E91-*f*R95 and *f*R59-*f*E91. Simultaneously, in the interdomain region, the new orientation of *h*R18 points towards the side chain of *h*N15 at the Ω loop in the productive closed state of HisF:HisH altering the interface dynamics (see Fig. S30). As a consequence, the side chain of *h*N15 alternates between hK181 and hR18 residues, while *h*N12 side chain establishes a relatively stable hydrogen bond with amide backbone of *h*N15 (3.1 ± 0.71 Å). Eventually the *h*K181 and *f*D98 salt bridge is strengthened upon productive HisF:HisH closure enhancing the communication between HisF subunit and HisH active site. This pair of residues is key for allosteric communication and, in fact, the mutation of *f*D98 disrupts the millisecond motions in IGPS.

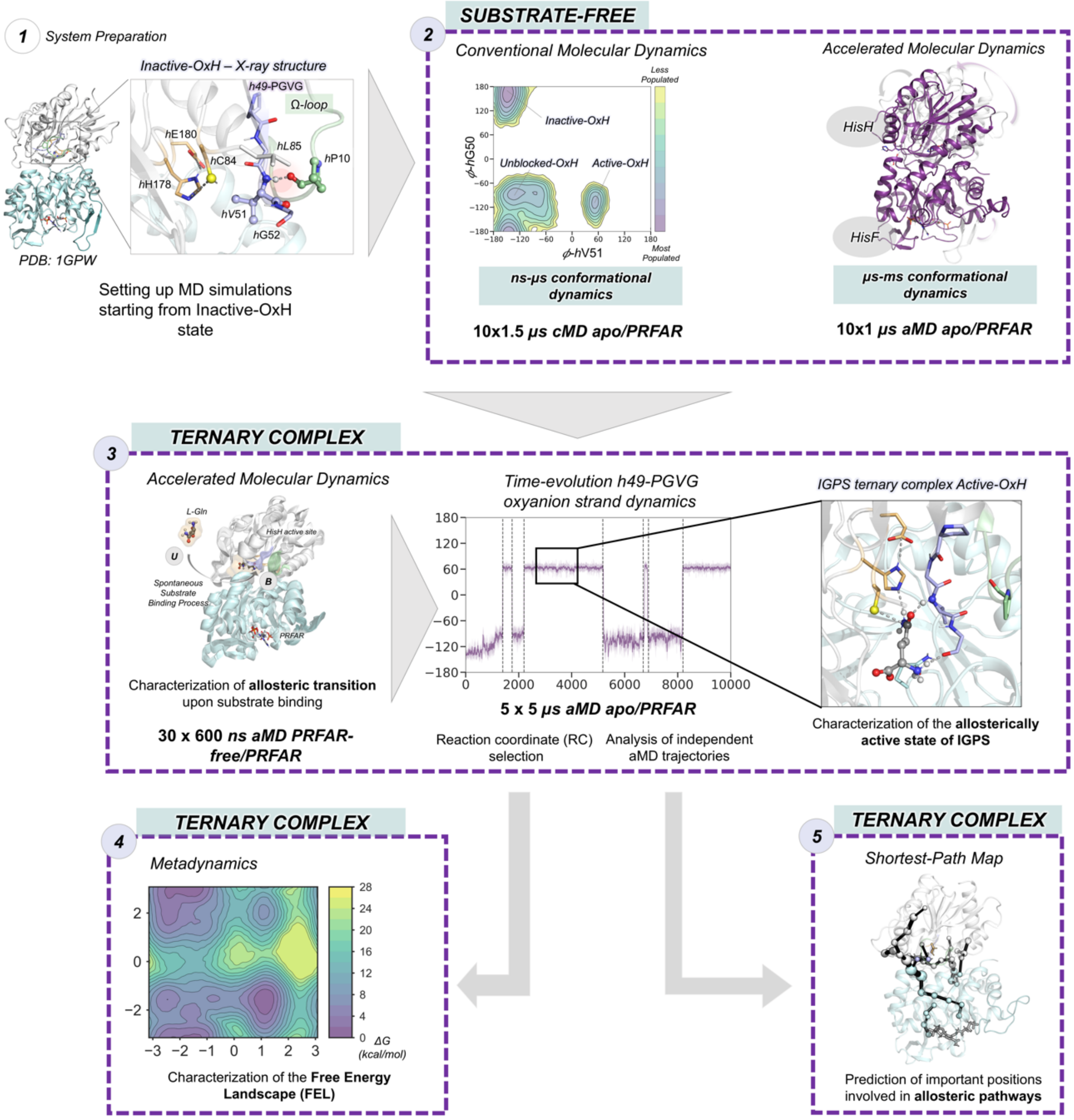


**Fig. S1.** Summary of the computational strategy used to characterize the molecular basis of the graded millisecond allosteric activation of *wt*IGPS. All simulations were performed starting from x-ray structure of IGPS in the inactive state (PDB ID 1GPW (chains A and B)).

*Conventional MD: HisH h49-PGVG oxyanion strand conformational dynamics along cMD simulations*

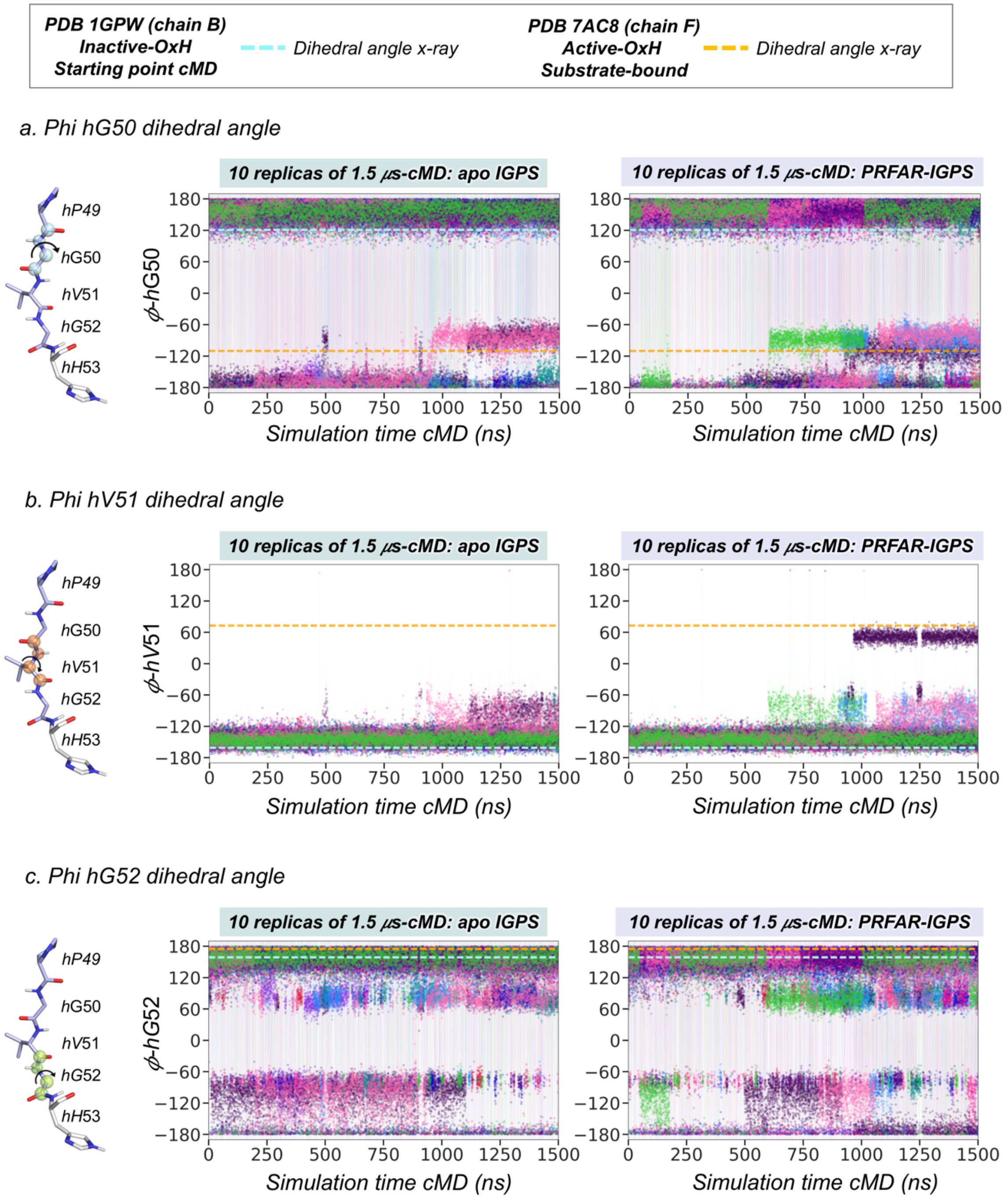

*Conventional MD: HisH oxyanion strand conformational dynamics along cMD simulations (continuation)*

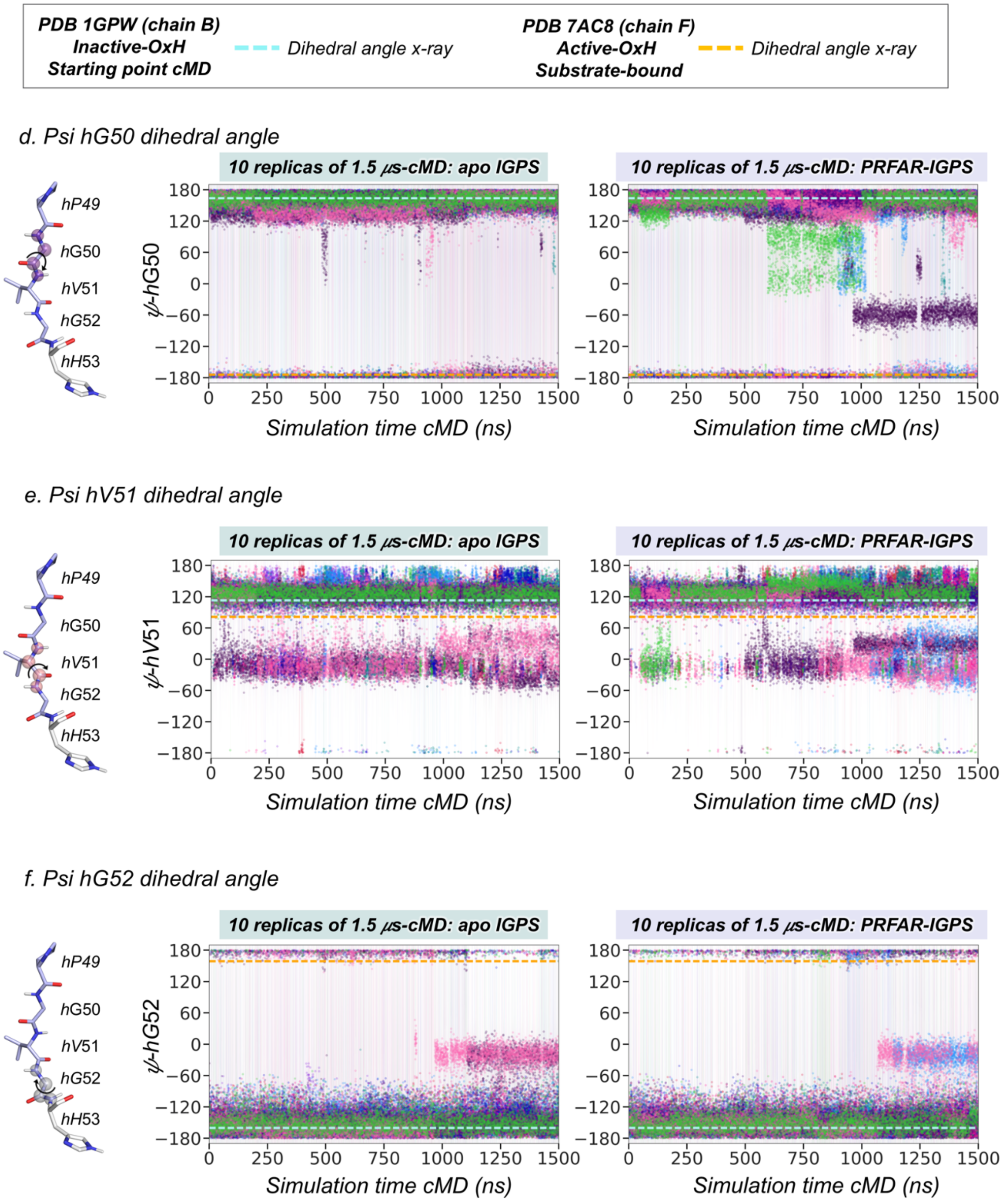


**Fig. S2. HisH *h*49-PVGV conformational dynamics along cMD simulations.** Plot of the most relevant dihedral angles of the *h*49-PGVG oxyanion strand for ten replicas of 1.5 μs conventional molecular dynamics (cMD) simulations in the apo-IGPS and PRFAR-IGPS states. Each replica is depicted in a different color. Horizontal cyan dashed lines indicate the value of the dihedral angle corresponding to the x-ray structure (PDB 1GPW (chain B)) used as starting point for cMD simulations. Horizontal orange dashed lines indicate the value of the dihedral angle corresponding to the x-ray structure of substrate-bound *h*C84A IGPS (PDB 7AC8 (chain F)) that displays an active conformation of the h49-PVGV oxyanion strand. The oxyanion strand residues are shown in light purple and the atoms involved in each dihedral angle are represented as spheres of different color. The cMD trajectory of PRFAR-IGPS where the Active-OxH state is sampled is represented in deep purple in all plots. (a) $\phi$ dihedral angle of *h*G50; (b) $\phi$ dihedral angle of *h*V51; (c) $\phi$ dihedral angle of *h*V51; (d) $\psi$ dihedral angle of *h*G50; (e) $\psi$ dihedral angle of *h*V51; (f) $\psi$ dihedral angle of *h*G52.

All dihedrals display a certain degree of flexibility along the 1.5 $\mu$s cMD simulations. Multiple orientations with respect to the x-ray dihedral angle are observed in most cases, being $\phi$-*h*G52 and $\psi$-*h*V51 the ones displaying more transitions in the nanosecond timescale. The dihedral angles $\phi$-*h*G50 and $\phi$-*h*V51 were selected as the most relevant ones for monitoring the formation of the Active-OxH state and the allosteric activation of IGPS. See Fig. S3 for a molecular representation of the most relevant states of the *h*49-PGVG oxyanion strand.

### *Conventional MD: HisH oxyanion strand conformational landscape ϕ-hV51 vs ϕ-hG50*

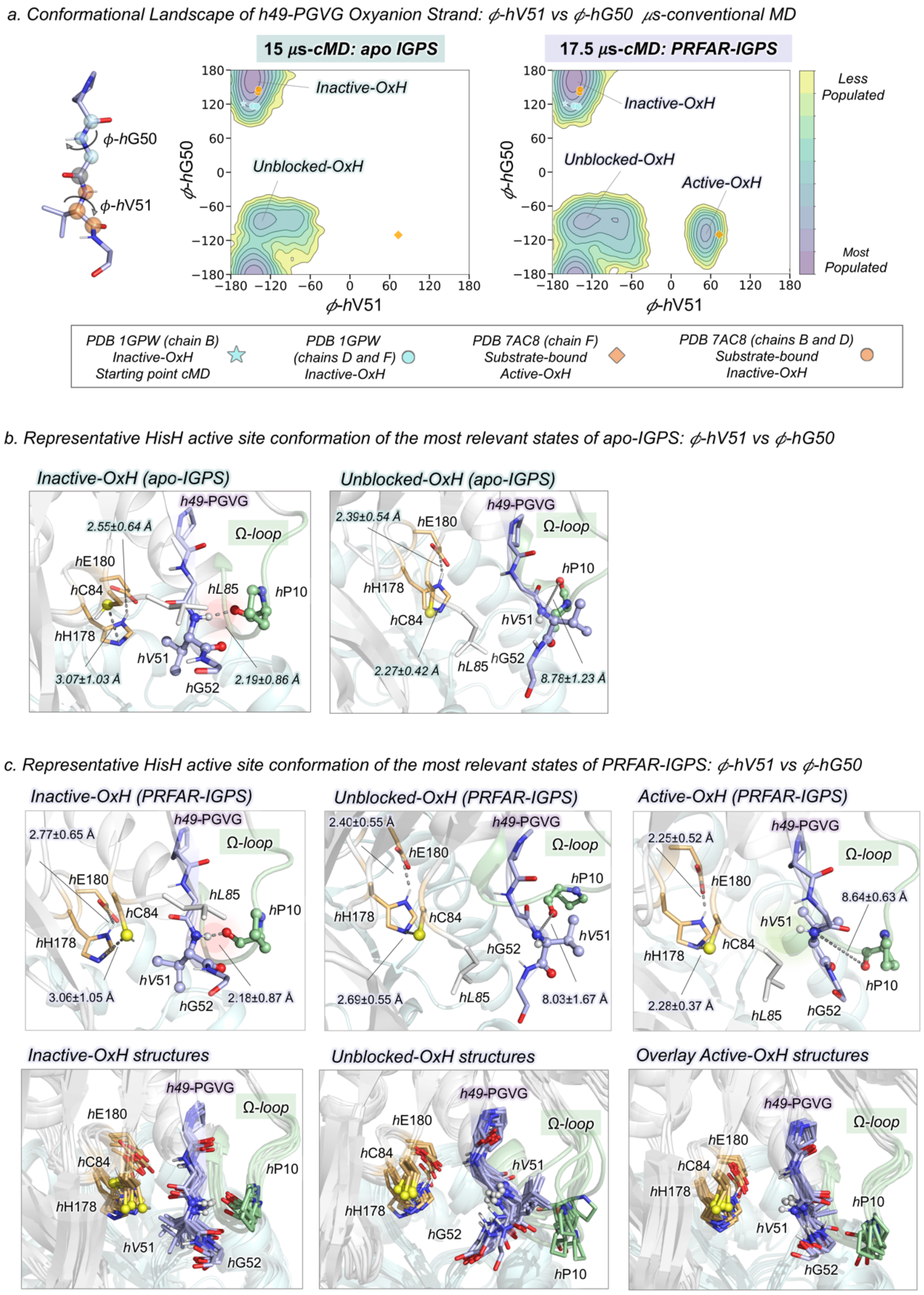

*Conventional MD: HisH oxyanion strand conformational landscape $\phi$-hV51 vs $\phi$-hG52*

*d. Conformational Landscape of h49-PGVG Oxyanion Strand: $\phi$-hV51 vs $\phi$-hG52 μs-conventional MD*

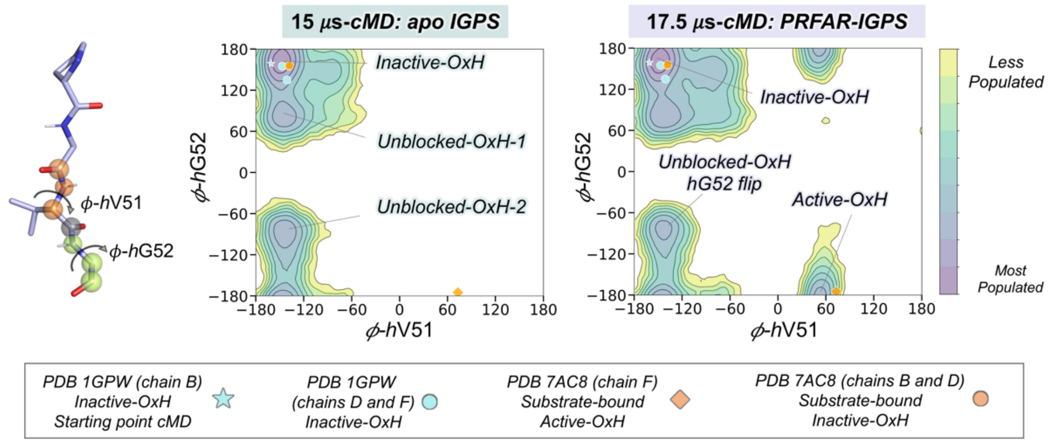


*e. Representative HisH active site conformations of the most relevant states of apo-IGPS: $\phi$-hV51 vs $\phi$-hG52*

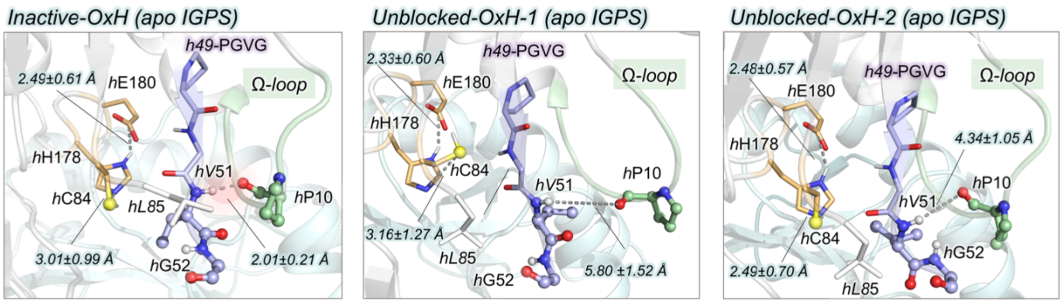


*f. Representative HisH active site conformations of the most relevant states of PRFAR-IGPS: $\phi$-hV51 vs $\phi$-hG52*

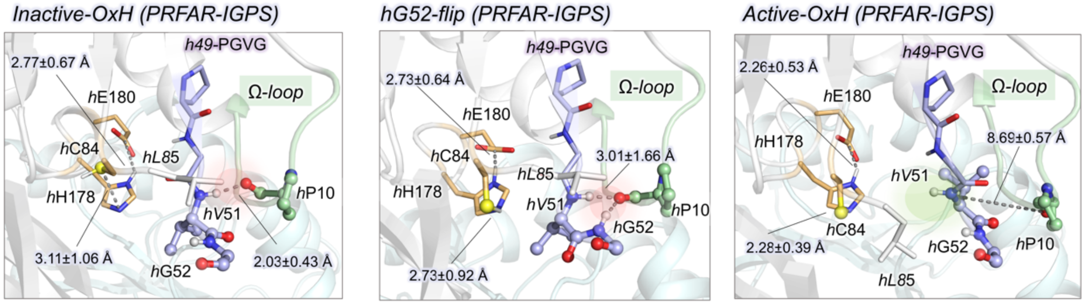


**Fig. S3. Conformational landscape of *h*49-PGVG oxyanion strand obtained from μs-conventional Molecular Dynamics (cMD) simulations**. The conformational landscape of the *h*49-PGVG oxyanion strand of apo-IGPS is constructed from an accumulated time of 15 μs of cMD simulations (10 replicas of 1.5 μs) while the oxyanion strand conformational landscape of PRFAR-IGPS is built from 17.5 μs of cMD simulation time (9 replicas of 1.5 μs and one replica of 4 μs). The conformational landscape of each state is clusterized into 20 different clusters. The clusters corresponding to the most populated regions are selected for further analysis (e.g. calculation of average distances). The *h*49-PGVG conformational landscape and the most representative conformations of each of the major conformational states have been obtained for both $\phi$-*h*V51/$\phi$-

*h*G50 and $\phi$-*h*V51/$\phi$-*h*G52 pairs of dihedral angles. (a) Conformational landscape of apo and PRFAR-IGPS constructed using the $\phi$ dihedral angles of *h*V51 and *h*G50. The values of the $\phi$ dihedral angles of *h*V51 and *h*G50 found in the x-ray structures corresponding to the three chains of PDB 1GPW are depicted in cyan and the three chains of PDB 7AC8 are represented in orange, respectively. The conformation used as starting point for cMD simulations is shown using the cyan star symbol. The conformation corresponding to the active oxyanion strand (Active-OxH) observed in *h*C84A IGPS is depicted using the orange diamond symbol. (b) Representative HisH active site structures of most populated states in apo-IGPS conformational landscape: inactive-OxH and unblocked-OxH. (c) Representative HisH active site structures of most populated states in PRFAR-IGPS conformational landscape: inactive-OxH, unblocked-OxH, and active-OxH. Overlay of eight representative structures corresponding to each conformational state of the oxyanion strand in PRFAR-IGPS. (d) Conformational landscape of apo and PRFAR-IGPS constructed using the $\phi$ dihedral angles of *h*V51 and *h*G52. The values of the $\phi$ dihedral angles of *h*V51 and *h*G52 found in the x-ray structures corresponding to the three chains of PDB 1GPW are depicted in cyan and the three chains of PDB 7AC8 are represented in orange, respectively. The conformation used as starting point for cMD simulations is shown using the cyan star symbol. The conformation corresponding to the active oxyanion strand (Active-OxH) observed in *h*C84A IGPS is depicted using the orange diamond symbol. (e) Representative HisH active site structures of most populated states in apo-IGPS conformational landscape: inactive-OxH, unblocked-OxH-1, unblocked-OxH-2. (f) Representative HisH active site structures of most populated states in PRFAR-IGPS conformational landscape: inactive-OxH, *h*G52-flip-OxH, and active-OxH. Relevant average distances (in Å) of each conformational state are depicted in green and purple for apo and PRFAR-bound states, respectively. The average distances are calculated considering all the structures included in each cluster. The HisH catalytic residues are highlighted in orange, Ω-loop residues in green, and the residues of the *h*49-PGVG oxyanion strand in purple. Other relevant HisF and HisH residues are shown in cyan and white, respectively. The atoms of *h*V51 and *h*P10 are shown as spheres. The red surface is used to show when the *h*P10-*h*V51 hydrogen bond is shown, blocking the formation of the *h*V51 oxyanion hole. The green surface is used to show when $H^N$ of *h*V51 is pointing toward the active site (Active-OxH with *h*V51 formed).

The active-OxH state of the *h*49-PGVG oxyanion strand is only sampled in PRFAR-IGPS. In the active-OxH state, the average values of the $\phi$-*h*V51 and $\phi$-*h*G50 are 51.3 ± 8.2 º and -107.5 ± 5.5 º. These values slightly deviate from the x-ray $\phi$-*h*V51 and $\phi$-*h*G50 dihedral angles of substrate-bound *h*C84A IGPS (PDB 7AC8 (chain F)) which are 73.2 º and -110.5 º, respectively. This displacement is associated with the presence of the substrate (see Fig. SX and SY). The three catalytic HisH active site residues display more flexibility in the Inactive-OxH state than in the Unblocked-OxH and Active-OxH states of the oxyanion-strand (overlay Fig. S3c). The distances between catalytic residues are monitored in Fig. S4.

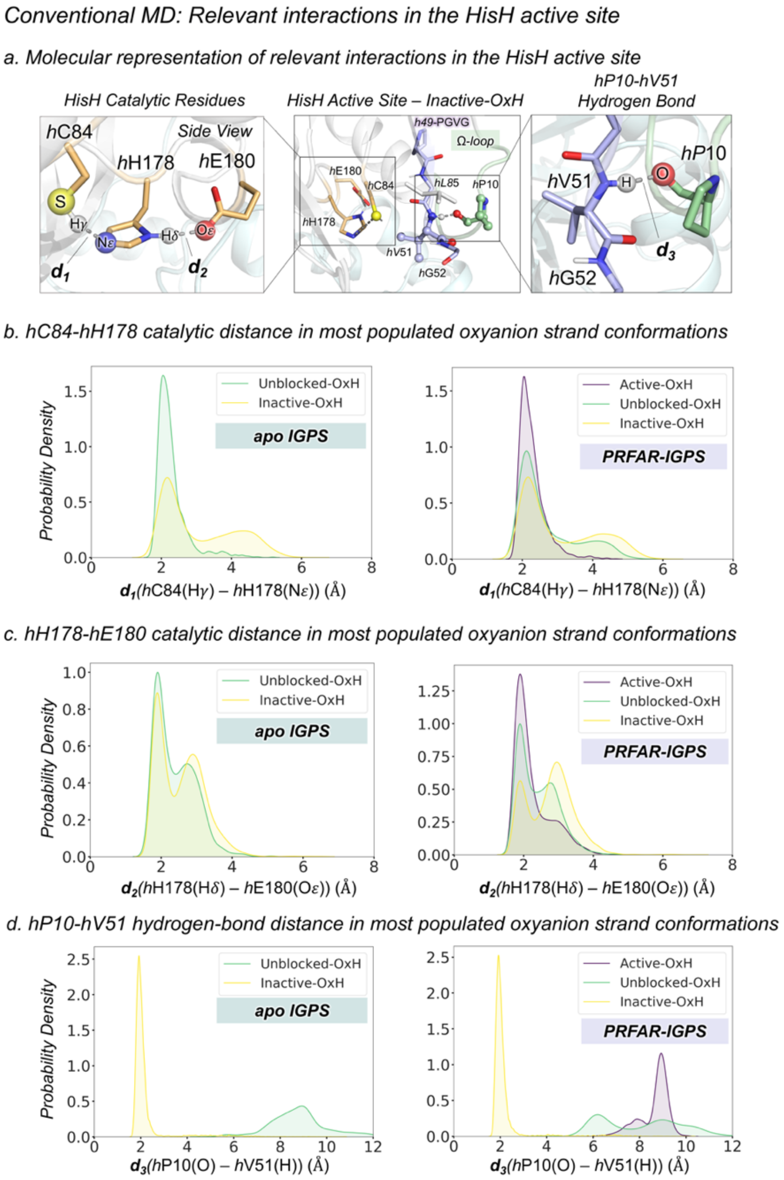


**Fig. S4. Relevant interactions in the HisH active site in cMD simulations**. (a) Structural representation of the relevant interactions in the HisH active site: interaction between catalytic residues *h*C84-*h*H178 ($d_1$) and *h*H178-*h*E180 ($d_2$), and the hydrogen between the Ω-loop residue *h*P10 and the oxyanion strand residue *h*V51 ($d_3$). (b) Probability density distribution of the *h*C84-*h*H178 distance in the most populated states of the oxyanion strand conformational landscape (see S3) of apo and PRFAR-IGPS. The distances are calculated considering all the structures included in the representative cluster of each state. The distance is monitored between the thiol hydrogen (Hγ) of *h*C84 and the epsilon nitrogen (Nε) of *h*H178. (c) Probability density distribution of the *h*H178-*h*E180 distance. The distance is monitored between the delta hydrogen (Hδ) of *h*H178 and the oxygen of the carboxylate group (Oε) of *h*E180. (d) Probability density distribution of the *h*P10-*h*V51 distance. The distance is monitored between the amide backbone hydrogen ($H^N$) of *h*V51 and the backbone oxygen (O) of *h*P10. The distances corresponding to the Inactive-OxH, Unblocked-OxH, and Active-OxH are shown in yellow, green, and purple respectively. All distances are in Å. The average distances of each state are shown in Fig. S3.

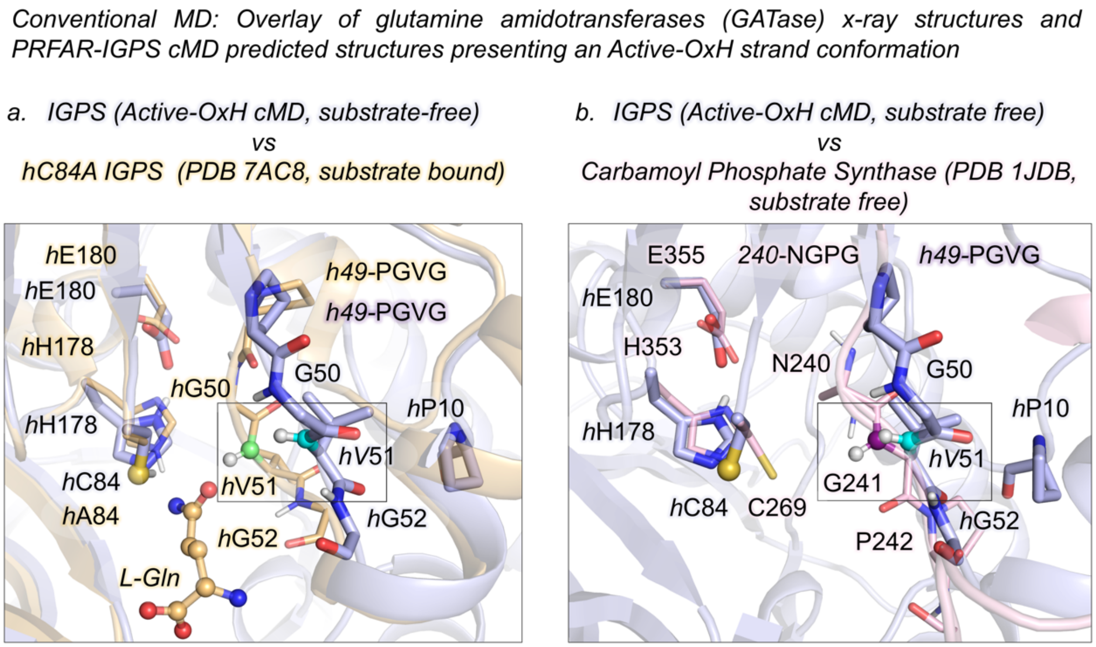


**Fig. S5. Overlay of class I glutamine amidotransferase (GATase) x-ray structures and PRFAR-IGPS cMD structures**. (a) Overlay of a representative substrate-free active-OxH PRFAR-IGPS (in purple) structure extracted from cMD simulations with the x-ray structure of the postulated active state of substrate-bound *h*C84A IGPS (PDB 7AC8 (chain F), in orange). The NH backbone of *h*V51 in the active-OxH PRFAR-IGPS (cMD) is highlighted in cyan while the NH backbone of *h*V51 corresponding to *h*C84A IGPS is highlighted in green. The glutamine (L-Gln) substrate present in the x-ray structure is highlighted with spheres. In both cases, the NH backbone is pointing toward the HisH catalytic residues (*h*C84/*h*A84). The conformation of the *h*49-PGVG oxyanion strand in the HisH active site show some differences due to the presence of the substrate in PDB 7AC8 (chain F). (b) Overlay of a representative substrate-free active-OxH PRFAR-IGPS (in purple) structure extracted from cMD simulations with the x-ray structure of the active state substrate-free carbamoyl phosphate synthase (PDB 1JDB (chain B), in light pink). In carbamoyl phosphate synthase the oxyanion strand is formed by 240-NGPG residues being G241 the residue responsible of forming the oxyanion hole (equivalent to *h*V51 in IGPS). The NH backbone of *h*V51 in the active-OxH PRFAR-IGPS is highlighted in cyan while the NH backbone of G241 corresponding to carbamoyl phosphate synthase is highlighted in magenta. In both cases, the NH backbone of *h*V51 is pointing toward the catalytic residues.

*a. Conventional MD: hL85 orientation in the HisH active site*

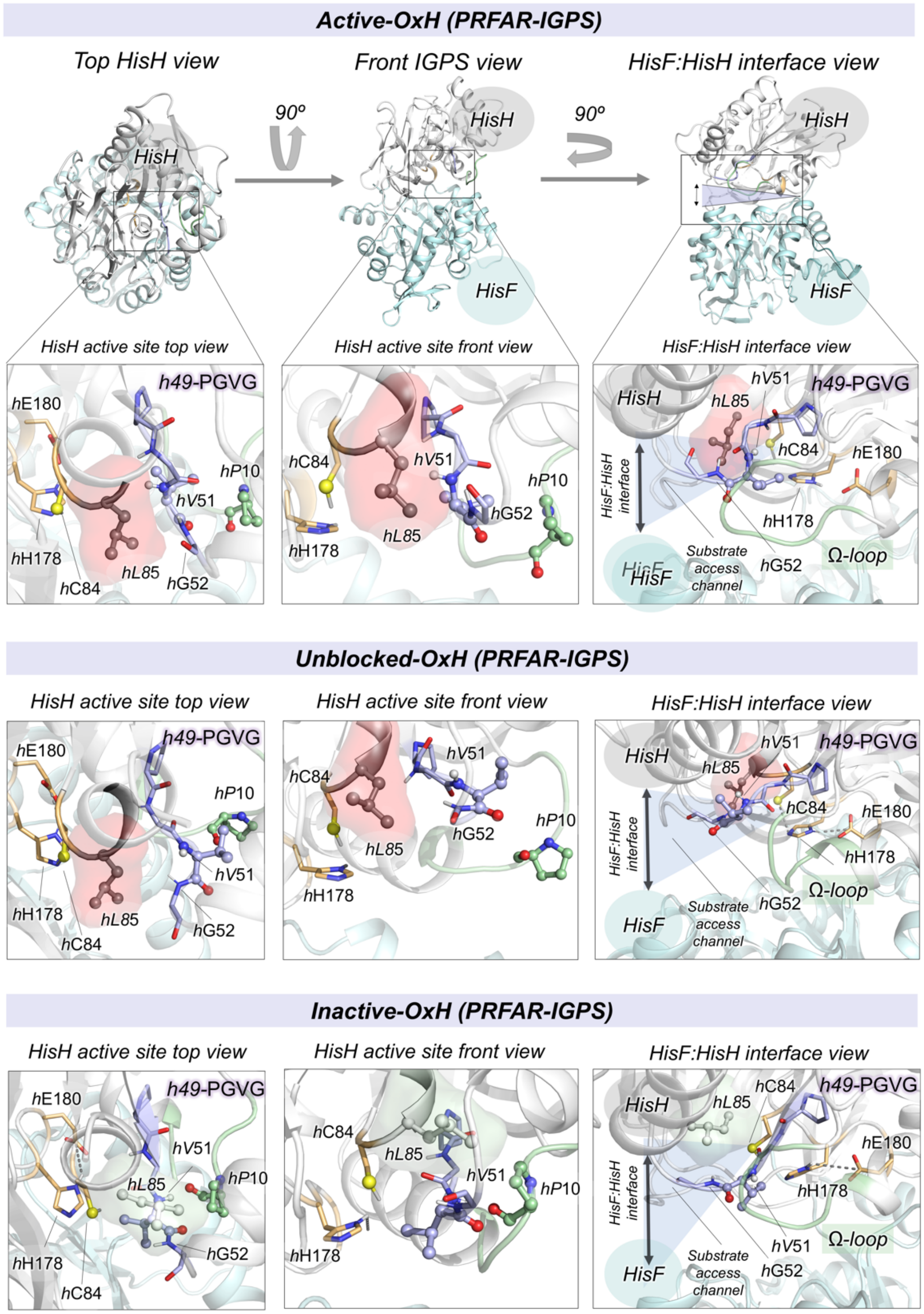


**Fig. S6. Orientation of *h*L85 in the HisH active site**. (a) Three different points of view of IGPS and HisH active site: top view (HisH is located above HisF), front view, and HisF:HisH interface view. Representative HisH active site conformations for the active-OxH, unblocked-OxH, and inactive-OxH *h*49-PGVG oxyanion strand states of PRFAR-IGPS sampled in cMD simulations. *h*L85 is highlighted as a red surface when it is blocking the access to the HisH active site (active-

OxH and unblocked-OxH states) and as a green surface when is not oriented toward the HisH active site (inactive-OxH). In the active-OxH and unblocked-OxH states, the *h*L85 side chain is positioned between the catalytic and oxyanion strand residues blocking the substrate access. In the inactive-OxH conformation is the side chain of *h*V51 that occupies the active site while *h*L85 is placed above the oxyanion strand residues (in the selected HisH active site views). The HisH catalytic residues are highlighted in orange, Ω-loop residues in green, and the residues of the *h*49-PGVG oxyanion strand in purple. Other relevant HisF and HisH residues are shown in cyan and white, respectively. The atoms of *h*L85, *h*V51, and *h*P10 are shown as spheres. In the HisF:HisH interface view, the substrate access channel is shown in blue.

*Conventional MD: Analysis of a 4 μs-cMD simulation (PRFAR-IGPS) displaying the hV51 oxyanion hole formation*

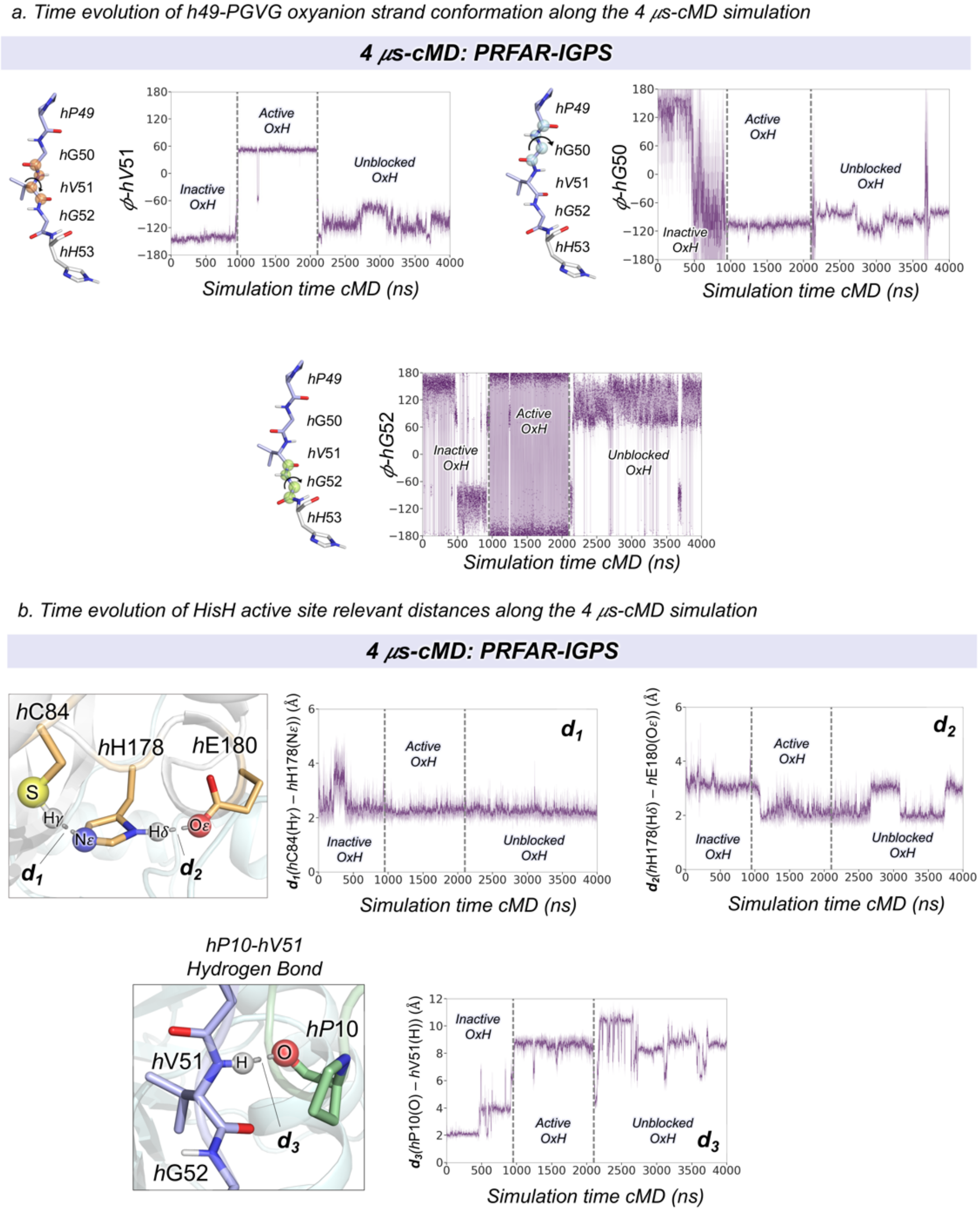


**Fig. S7. Analysis of a representative 4 μs-cMD simulation displaying the *h*V51 oxyanion hole formation**. (a) Plot of the most relevant dihedral angles of the *h*49-PGVG oxyanion strand along a 4 μs-cMD simulation: ϕ dihedral angle of *h*G50; ϕ dihedral angle of *h*V51; ϕ dihedral angle of *h*V51. Vertical gray dashed lines indicate the *h*V51 oxyanion hole formation. (b) Plot of the most relevant HisH active site distances (see Fig. S4) along the 4 μs-cMD simulation. Interaction between catalytic residues *h*C84-*h*H178 ($d_1$) and *h*H178-*h*E180 ($d_2$), and the hydrogen between the Ω-loop residue *h*P10 and the oxyanion strand residue *h*V51 ($d_3$). The formation of the Active-OxH is preceded by the disruption of the *h*P10 and *h*V51 hydrogen bond. All distances are in Å.

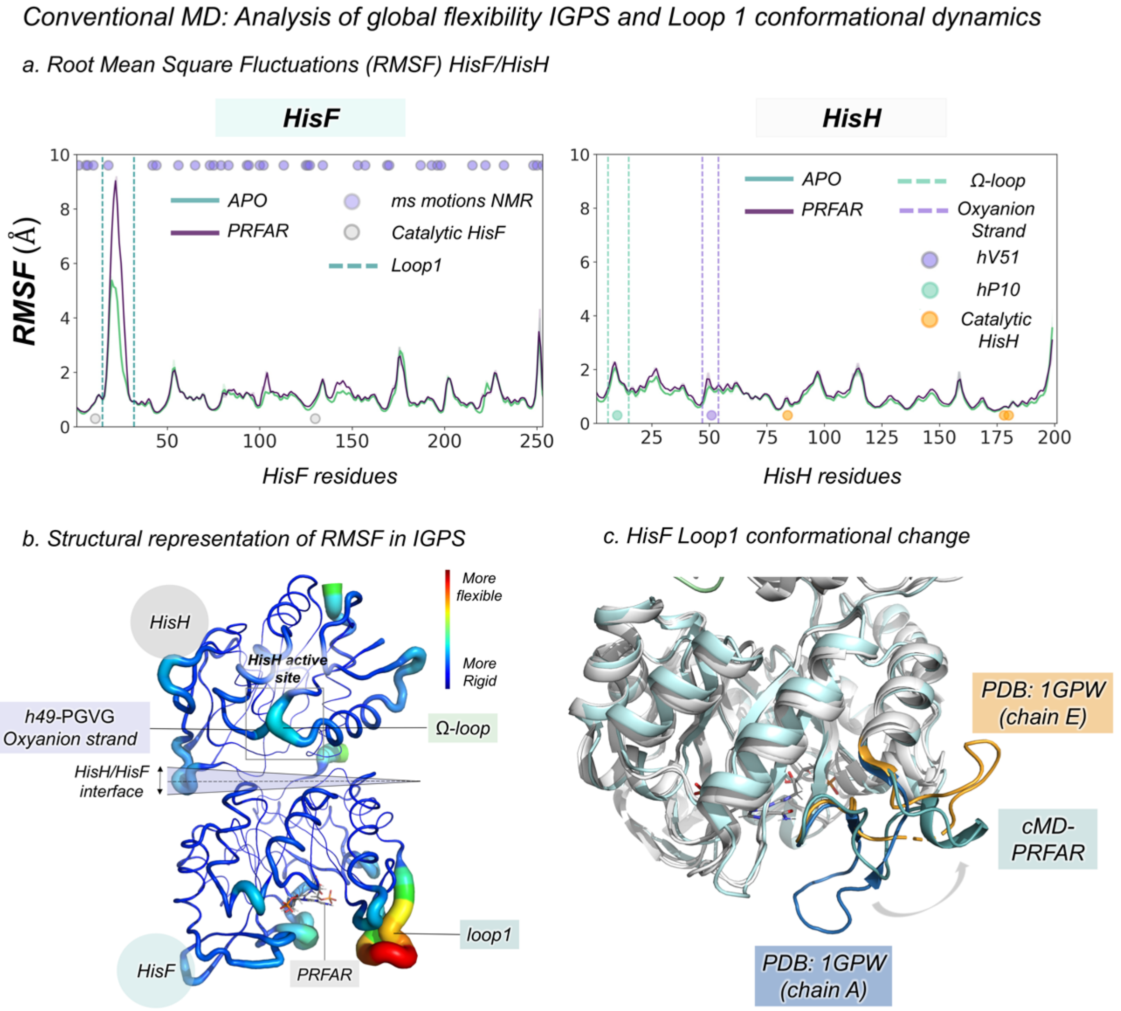


**Fig. S8. Analysis of global flexibility of IGPS and Loop1 conformational dynamics**. (a) Plot of the Root Mean Square Fluctuation (RMSF, in Å) for apo (green) and PRFAR-IGPS (purple) obtained from ten replicas of 1.5 μs cMD simulations. The plot is divided into HisF (left) *f*1-*f*253 residues and HisH (right) *h*1-*h*201 residues. The most relevant catalytic residues are highlighted in gray and in orange for HisF and HisH, respectively. The residues displaying NMR millisecond motions in HisF in the presence of PRFAR reported by Lisi and Loria(20) are highlighted in purple in the top of the plot. In HisH, the positions of *h*P10 and *h*V51 are shown in green and purple respectively. Vertical green and purple dashed lines indicate the position of the Ω-loop and oxyanion strand, respectively. In terms of global flexibility, the most significant differences are in *f*loop1 (see SI Extended text above for a complete description). (b) Structural representation of RMSF in the IGPS structure. The most flexible regions are represented in red (thicker loops), and the least flexible in blue (thinner loops). (c) Representative conformation of loop1 extracted from cMD simulations (teal). Overlay of IGPS structures with a closed loop1 conformation (PDB 1GPW chain A, in blue) and open loop1 conformation (PDB 1GPW chain E, in orange). Loop1 transitions from closed to open conformation in both apo and PRFAR-IGPS cMD simulations.

*Conventional MD: HisF conformational dynamics*

*a. HisF:HisH interface and HisF salt bridge network interactions*

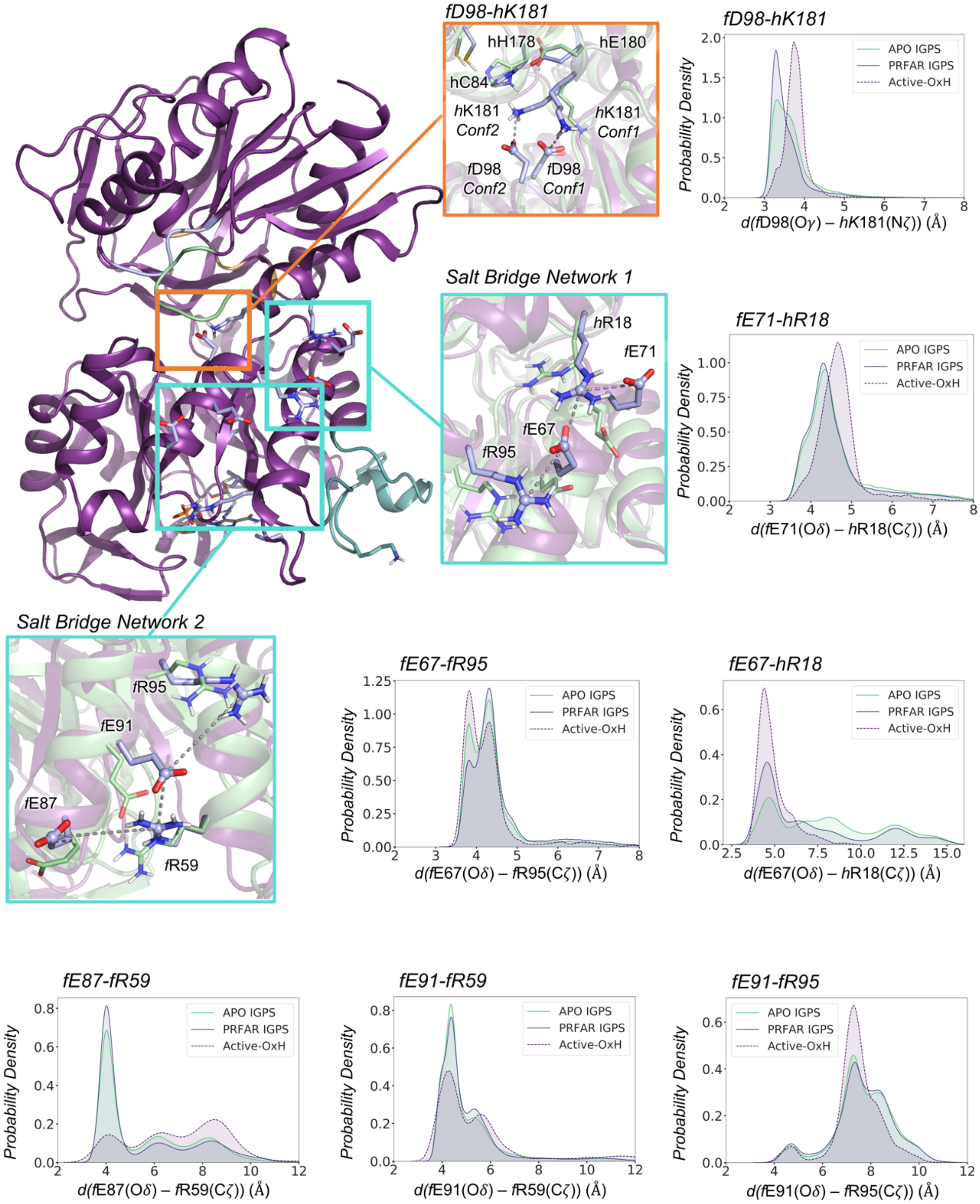

*Conventional MD: HisF conformational dynamics (continuation)*

*b. HisF hydrophobic cluster and fK19-PRFAR interactions*

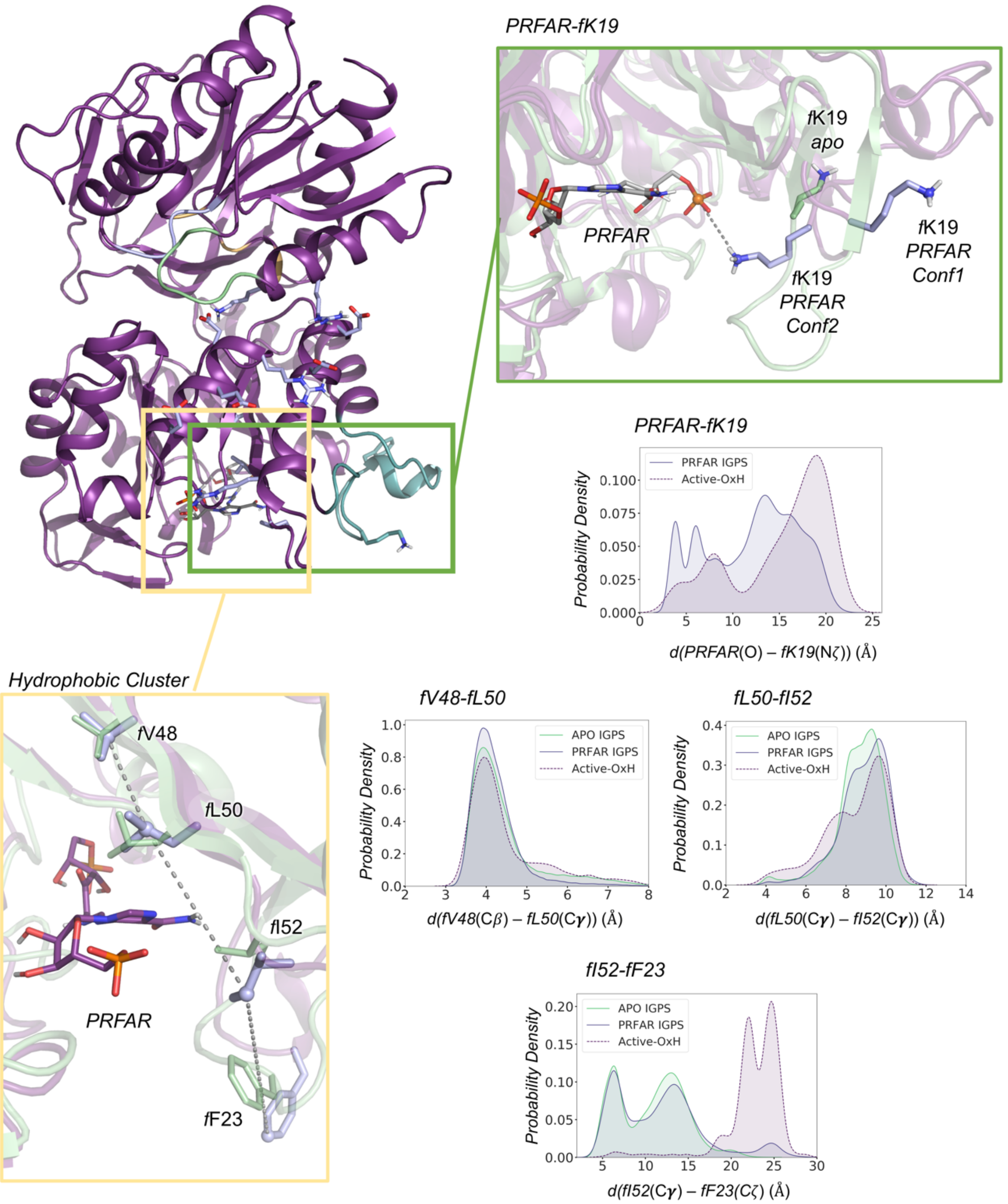


**Fig. S9. HisF conformational dynamics in cMD simulations.** Analysis of the most relevant interactions in HisF subunit in the apo and PRFAR-IGPS states along the cMD simulations. The selected distances were described by Rivalta et al.(4) and shown to be relevant for analysing the effects of PRFAR. The global conformation of PRFAR-IGPS is shown in deep purple. Loop1 is represented in teal, the Ω-loop in green, and the oxyanion strand in light purple. Overlay of the side chains of representative conformationas in the apo and IGPS-PRFAR states are shown in green and light purple, respectively. (a) Molecular representation of HisF salt-bridge network (highlighted in cyan squares) and *f*D98-*h*K181 HisF:HisH interaction (highlighted in an orange square). Probability density distribution of the most relevant distances of the salt-bridge network and *f*D98-*h*K181 distance. The distances of the salt bridge network are calculated between the carbon atom of the carboxylate group of the glutamate side chain and the carbon atom of the guanidinium group of the arginine residues. The distance of the *f*D98-*h*K181 interaction is calculated between the carbon atom of the carboxylate group of *f*D98 side chain and the nitrogen of the side chain of *h*K181. (b) Molecular representation of HisF hydrophobic cluster (highlighted in a yellow square) and PRFAR-*f*K19 HisF:HisH interaction (highlighted in a green square). Probability density distribution of the most relevant distances of the hydrophobic cluster and PRFAR-*f*K19 distance. The distances between the residues forming the hydrophobic cluster are monitored between the beta carbon of *f*V48, the gamma carbon of *f*L50, the gamma carbon of *f*I52, and the zeta carbon of *f*F23. The distance of the PRFAR-*h*K19 interaction is calculated between the phosphorus atom of PRFAR and the nitrogen of the side chain of *h*K19. In the probability density plots the apo, PRFAR-IGPS, and Active-OxH PRFAR-IGPS distances are shown in green, purple, and dashed purple lines, respectively. All distances are in Å. See SI Extended text for a complete description of the results.

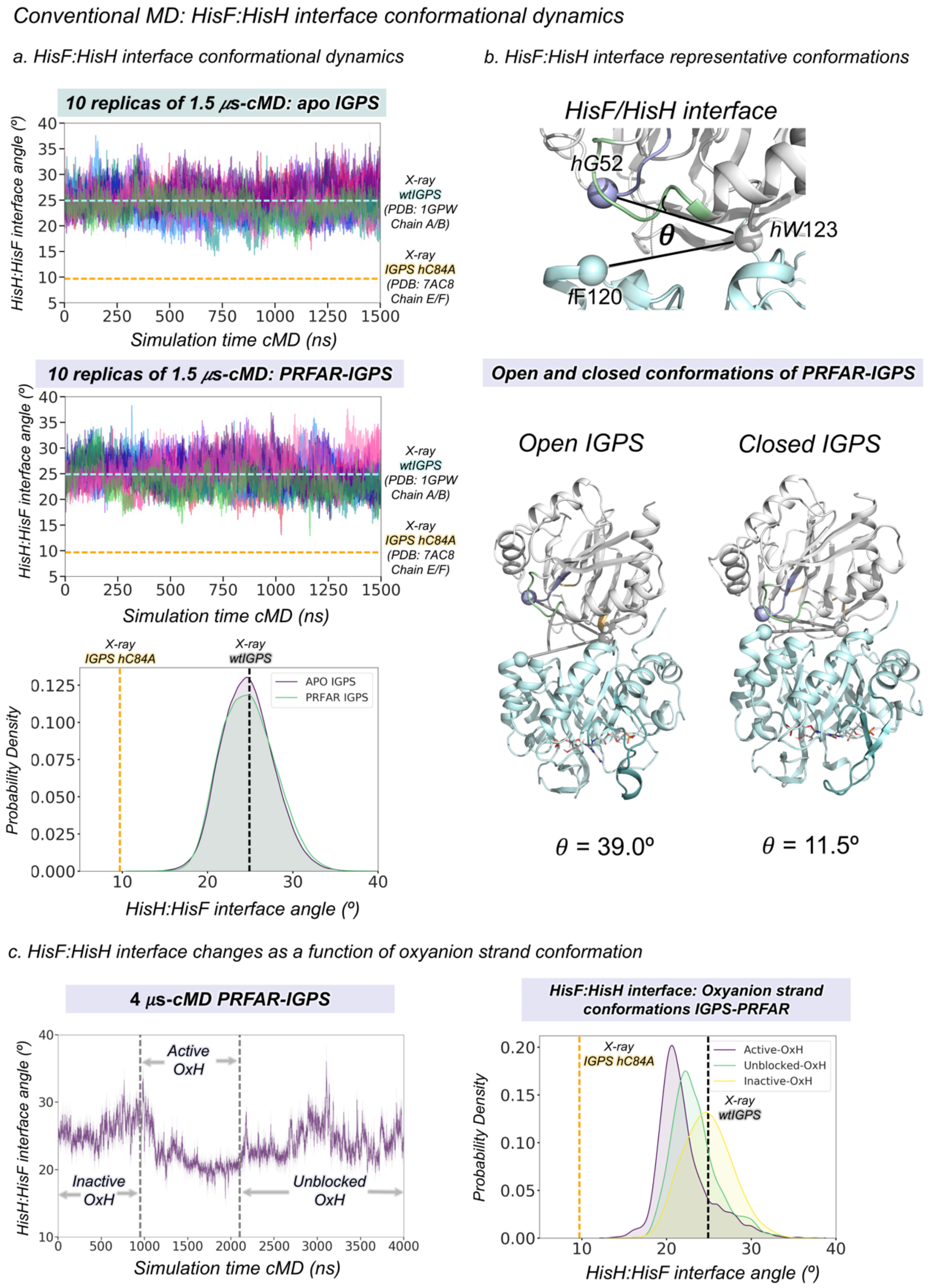


**Fig. S10. HisF:HisH interface conformational dynamics in cMD simulations.** (a) Plot of the HisF:HisH interface angle (θ) for ten replicas of 1.5 µs cMD simulations in the apo-IGPS and

PRFAR-IGPS states. Each replica is depicted in a different color. Horizontal cyan dashed lines indicate the HisF:HisH interface angle ($\theta$ = 24.9 º) found in the x-ray structure (PDB 1GPW chains A/B) used as starting point for cMD simulations. Horizontal orange dashed lines indicate the HisF:HisH interface angle ($\theta$ = 9.7 º) found in the x-ray structure of substrate-bound *h*C84A IGPS (PDB 7AC8 chains E/F) that displays an active conformation of the oxyanion strand. Probability density distribution for the HisF:HisH interface angle in the apo and PRFAR-IGPS states. Vertical gray dashed line corresponds to the 1GPW (chains A/B) x-ray HisF:HisH interface angle and the vertical dashed orange line to the 7AC8 (chains E/F) x-ray HisF:HisH interface angle. (b) Representative conformations of an open and closed conformations of the HisF:HisH interface sampled in PRFAR-IGPS. (c) Plot of the HisF:HisH interace angle along a representative 4 $\mu$s-cMD simulation that displays the formation of the *h*V51 oxyanion hole. Vertical dashed gray lines indicate the range of the Active-OxH state of the oxyanion strand. The formation of the *h*V51 oxyanion hole is correlated with a partial closure of the HisF:HisH interface (see SI Extended text for a complete description) Probability density distribution of the HisF:HisH interface angle for the different states of the oxyanion strand obtained in cMD simulations. The population of the Active-OxH oxyanion strand decreases the HisF:HisH interface angle with respect to the Inactive-OxH and Unblocked-OxH states. However, the values of the HisF:HisH interface angle are still far from the productive closure observed in PDB 7AC8 (chains E/F). The angle of the HisF:HisH interface is calculated from the alpha-carbons of *f*F120, *h*W123 and *h*G52 as indicated by Rivalta and coworkers.(4)

*Accelerated MD: HisH oxyanion strand conformational dynamics*

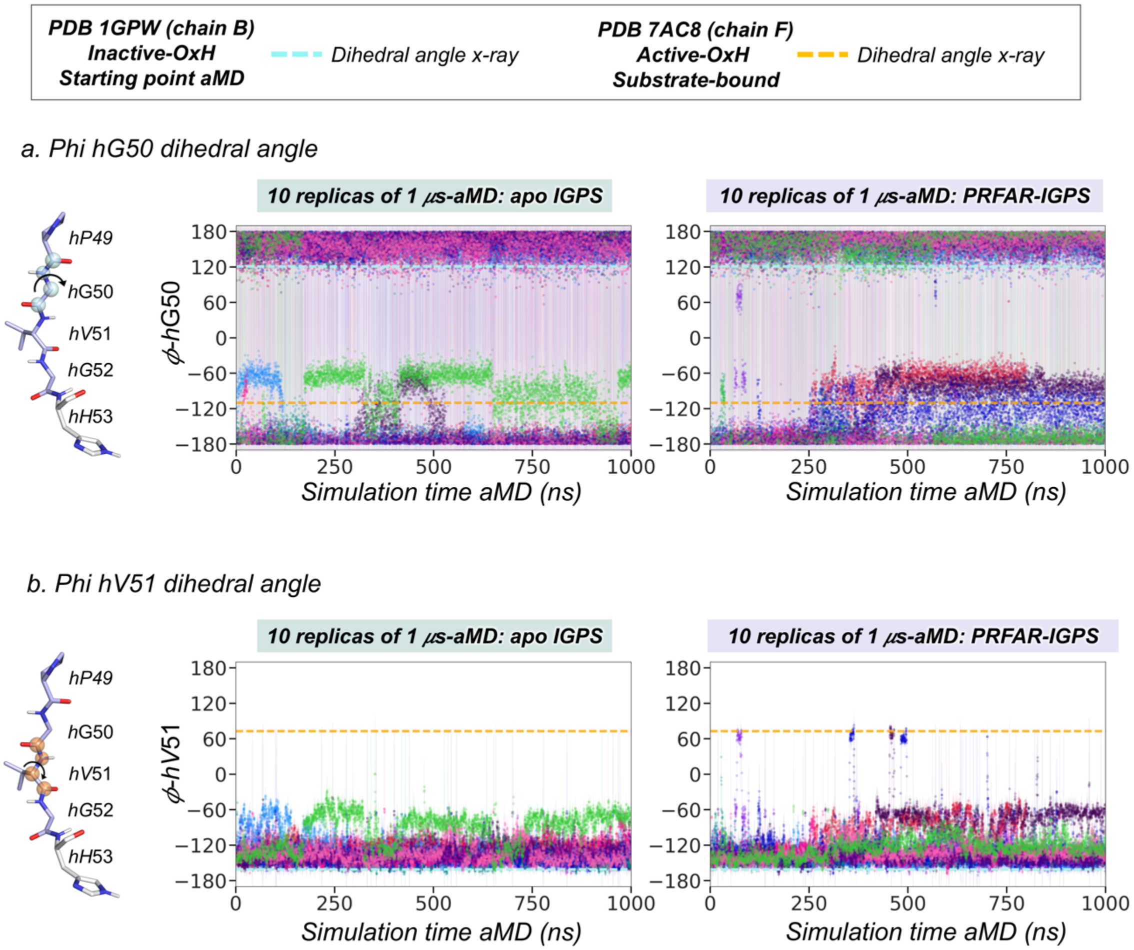


**Fig. S11. HisH *h*49-PVGV conformational dynamics along aMD simulations.** Plot of the most relevant dihedral angles of the *h*49-PGVG oxyanion strand for ten replicas of 1 μs accelerated molecular dynamics (aMD) simulations in the apo-IGPS and PRFAR-IGPS states. Each replica is depicted in a different color. Horizontal cyan dashed lines indicate the value of the dihedral angle corresponding to the x-ray structure (PDB 1GPW (chain B)) used as starting point for aMD simulations. Horizontal orange dashed lines indicate the value of the dihedral angle corresponding to the x-ray structure of substrate-bound *h*C84A IGPS (PDB 7AC8 (chain F)) that displays an active conformation of the h49-PVGV oxyanion strand. The oxyanion strand residues are shown in light purple and the atoms involved in each dihedral angle are represented as spheres of different color. (a) ϕ dihedral angle of *h*G50; (b) ϕ dihedral angle of *h*V51. In aMD, multiple short-lived formations of the *h*V51 oxyanion hole are observed. See Fig. S12 for a molecular representation of the most relevant states.

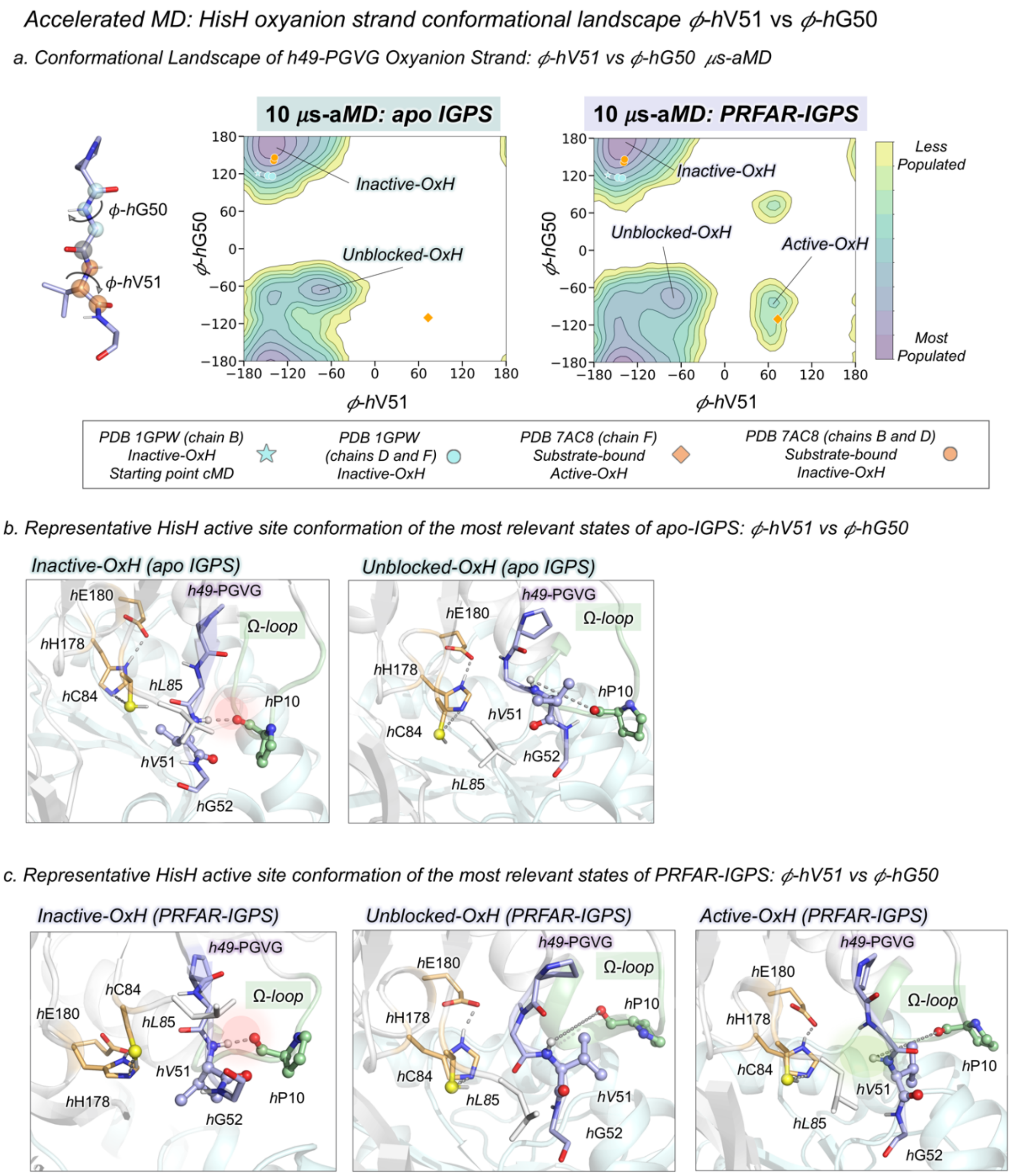


**Fig. S12. Conformational Landscape of *h*49-PGVG Oxyanion Strand obtained from accelerated Molecular Dynamics (aMD) simulations.** The conformational landscape of the *h*49-PGVG oxyanion strand of apo and PRFAR-IGPS is constructed from an accumulated time of 10 μs of aMD simulations (10 replicas of 1 μs). The conformational landscape of each state is clusterized into 20 different clusters. (a) Conformational landscape of apo and PRFAR-IGPS constructed using the $\phi$ dihedral angles of *h*V51 and *h*G50. The values of the $\phi$ dihedral angles of *h*V51 and *h*G50 found in the x-ray structures corresponding to the three chains of PDB 1GPW are depicted in cyan and the three chains of PDB 7AC8 are represented in orange, respectively. The conformation used as starting point for cMD simulations is shown using the cyan star symbol. The conformation corresponding to the active oxyanion strand (Active-OxH) observed in *h*C84A IGPS is depicted using the orange diamond symbol. (b) Representative HisH active site structures of most populated states in apo-IGPS conformational landscape: inactive-OxH and unblocked-OxH. (c) Representative HisH active site structures of most populated states in PRFAR-IGPS conformational landscape: inactive-OxH, unblocked-OxH, and active-OxH.

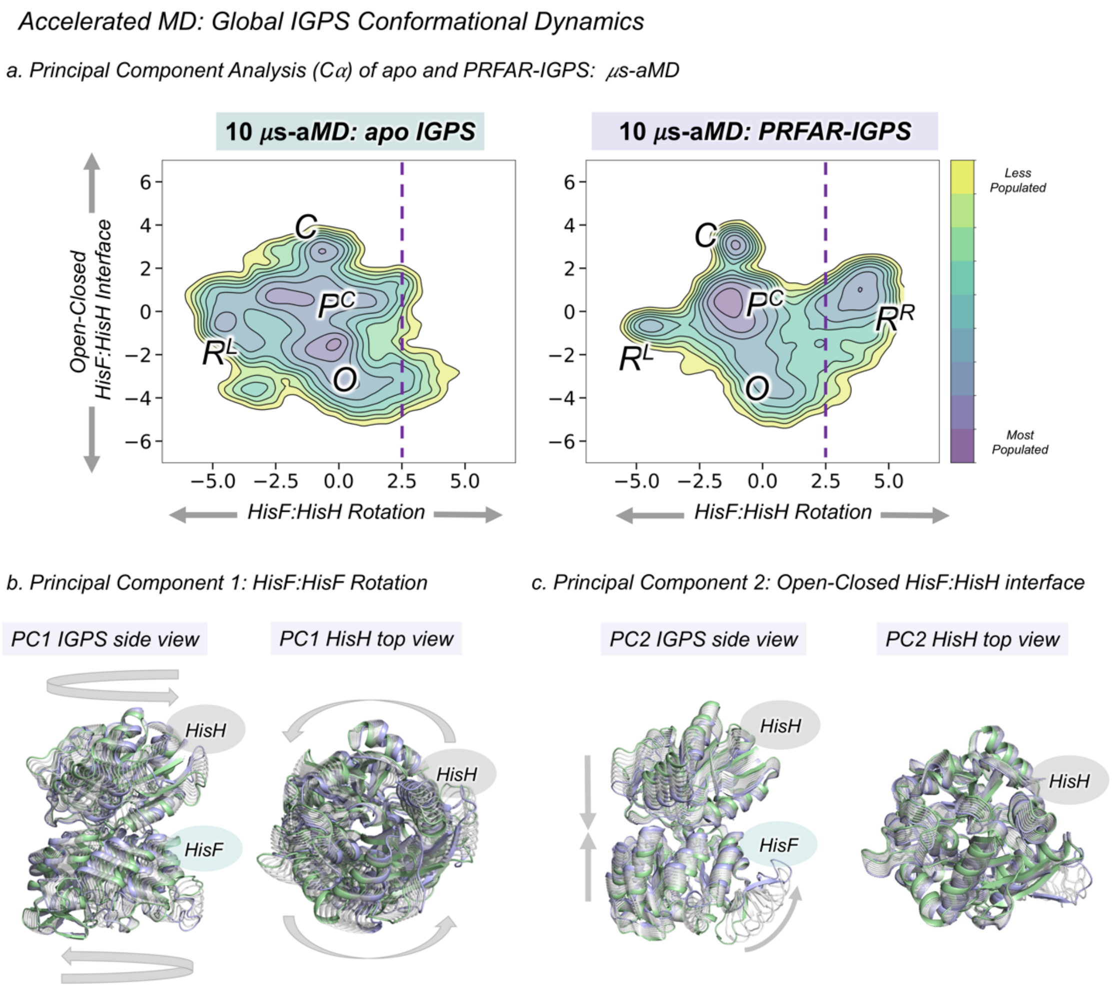

Accelerated MD: Global IGPS Conformational Dynamics
a. Principal Component Analysis (Cα) of apo and PRFAR-IGPS: μs-aMD
10 μs-aMD: apo IGPS
10 μs-aMD: PRFAR-IGPS
Open-Closed HisF:HisH Interface
HisF:HisH Rotation
C
P^C
R^L
O
R^R
Less Populated
Most Populated
6
4
2
0
−2
−4
−6
−5.0
−2.5
0.0
2.5
5.0
b. Principal Component 1: HisF:HisF Rotation
c. Principal Component 2: Open-Closed HisF:HisH interface
PC1 IGPS side view
PC1 HisH top view
PC2 IGPS side view
PC2 HisH top view
HisH
HisF

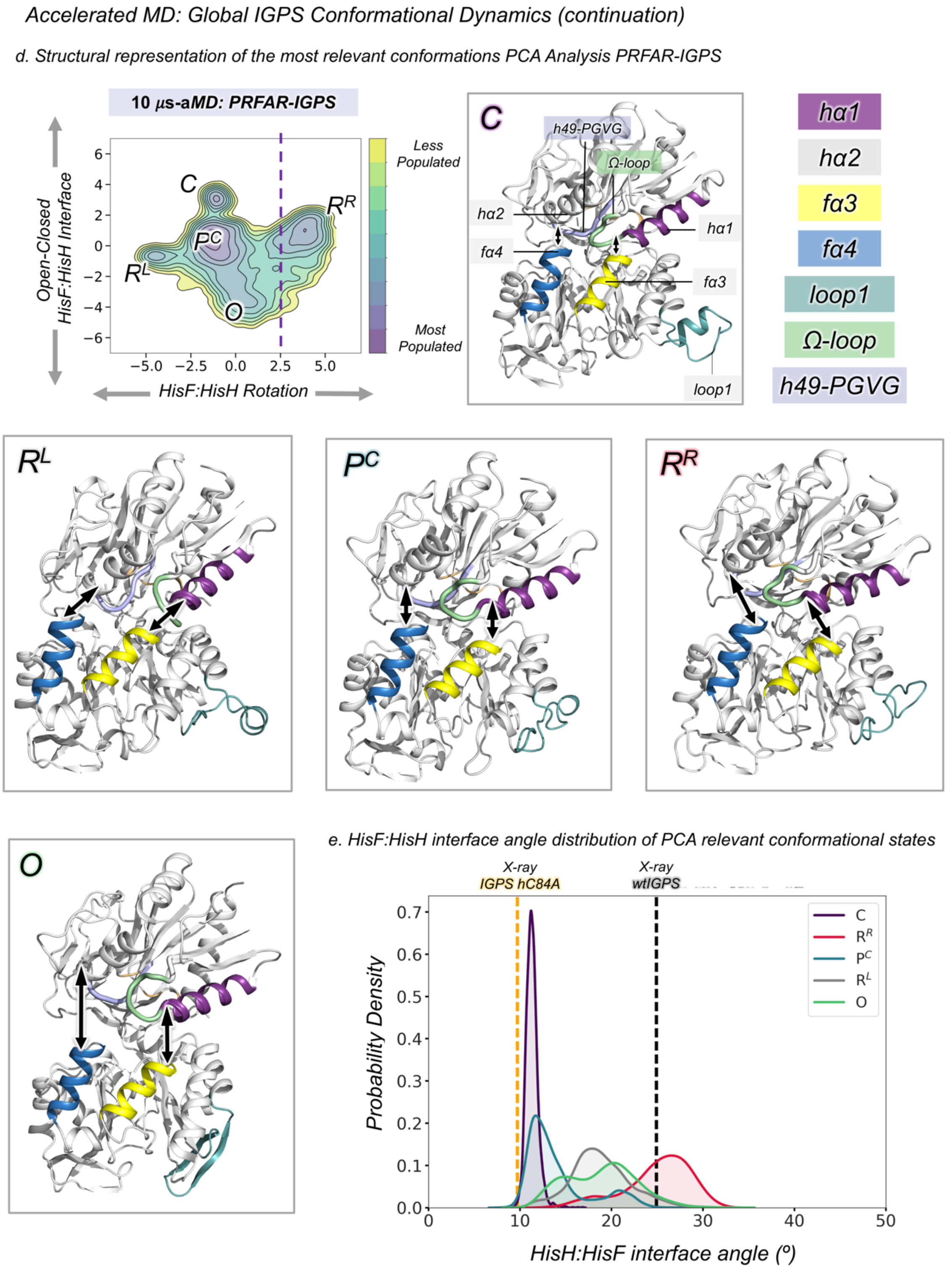


**Fig. S13. Global IGPS conformational dynamics in aMD simulations.** Principal component analysis of aMD simulations. (a) Principal Component (PC) analysis considering all alpha-carbons PRFAR-IGPS states reconstructed from ten replicas of 1 μs aMD simulations. PC1 indicates the counter-clock rotation of HisF and HisH subunits with respect to the HisF:HisH interface while PC2 represents the open-closed transition of the HisF:HisH interface. The apo-IGPS simulations are projected into the PC space of PRFAR-IGPS for a direct comparison. The vertical purple dashed line indicate the region of PC1 space not visited in apo-IGPS simulations. (b) Structural representation of PC1 and PC2 motions from side and top views of IGPS (see Fig. S6). The arrows indicate the direction of motions when going from negative (green cartoon) to positive (purple cartoon) values of the PC space. (c) Structural representation of PC2 motion. The arrows indicate the direction of motions when going from negative (green cartoon) to positive (purple cartoon) values of the PC space. (d) Representative conformations of the most populated states of the PC landscape of PRFAR-IGPS. The *h*α1, *h*α2, *f*α3, and *f*α4 helices are shown in purple, gray, yellow, and blue, respectively. Loop1 is represented in teal, the Ω-loop in green, and the oxyanion strand in light purple. (e) Probability density distribution of the HisF:HisH interface angle in the most relevant states visited in the aMD simulations. Vertical gray dashed line indicate the HisF:HisH interface angle found in the x-ray structure (PDB 1GPW chains A/B) used as starting point for aMD simulations. Vertical orange dashed lines indicate the HisF:HisH interface angle found in the x-ray structure of substrate bound *h*C84A IGPS (7AC8 chain E/F) that displays an active conformation of the oxyanion strand. The angle of the HisF:HisH interface is calculated from the alpha-carbons of *f*F120, *h*W123 and *h*G52. aMD simulations show that IGPS can attain the productive closure (C state) even when the substrate is not present. See SI Extended text for a complete description of these results.

*Accelerated MD: hV51 oxyanion hole formation and productive HisF:HisH closure are not correlated*

*a. Conformational Landscape of HisF:HisH interface angle and ϕ-hV51 dihedral angle*

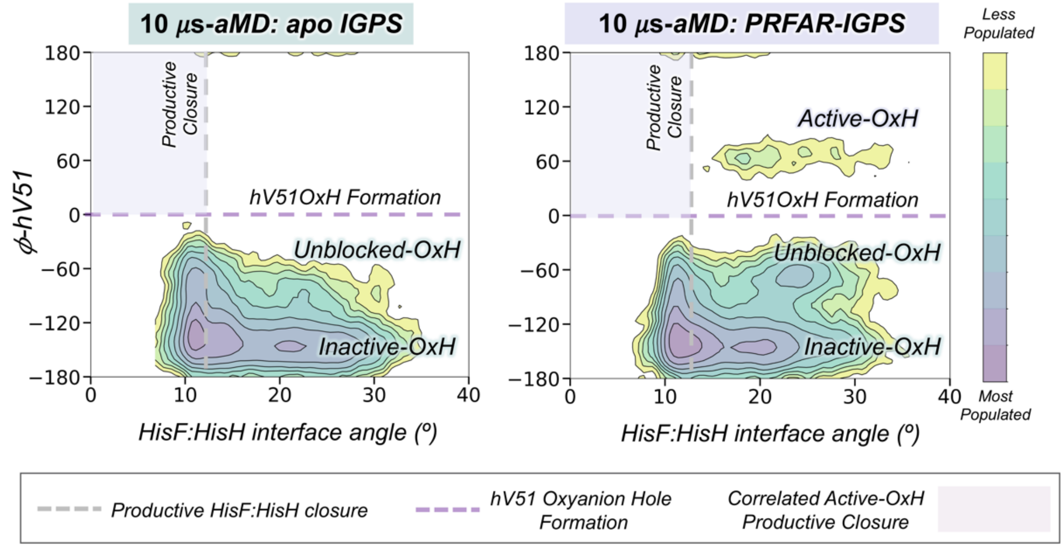


**Fig. S14. Uncorrelated HisF:HisH interface and oxyanion-strand conformational dynamics in aMD simulations.** Conformational landscape constructed using the HisF:HisH interface angle and ϕ dihedral angle of *h*V51 obtained from accelerated Molecular Dynamics (aMD) simulations of apo and PRFAR-IGPS states. Vertical gray dashed line indicates productive closure (HisF:HisH interface angle below 12 º). Horizontal purple dashed line indicate *h*V51 oxyanion hole formation (ϕ-*h*V51 above 0º). The purple area in the plot indicate the region of the conformational landscape with a productively closed HisF:HisH interface and a *h*V51 oxyanion hole formed. As the results show, the two events are not correlated in substrate-free aMD simulations.

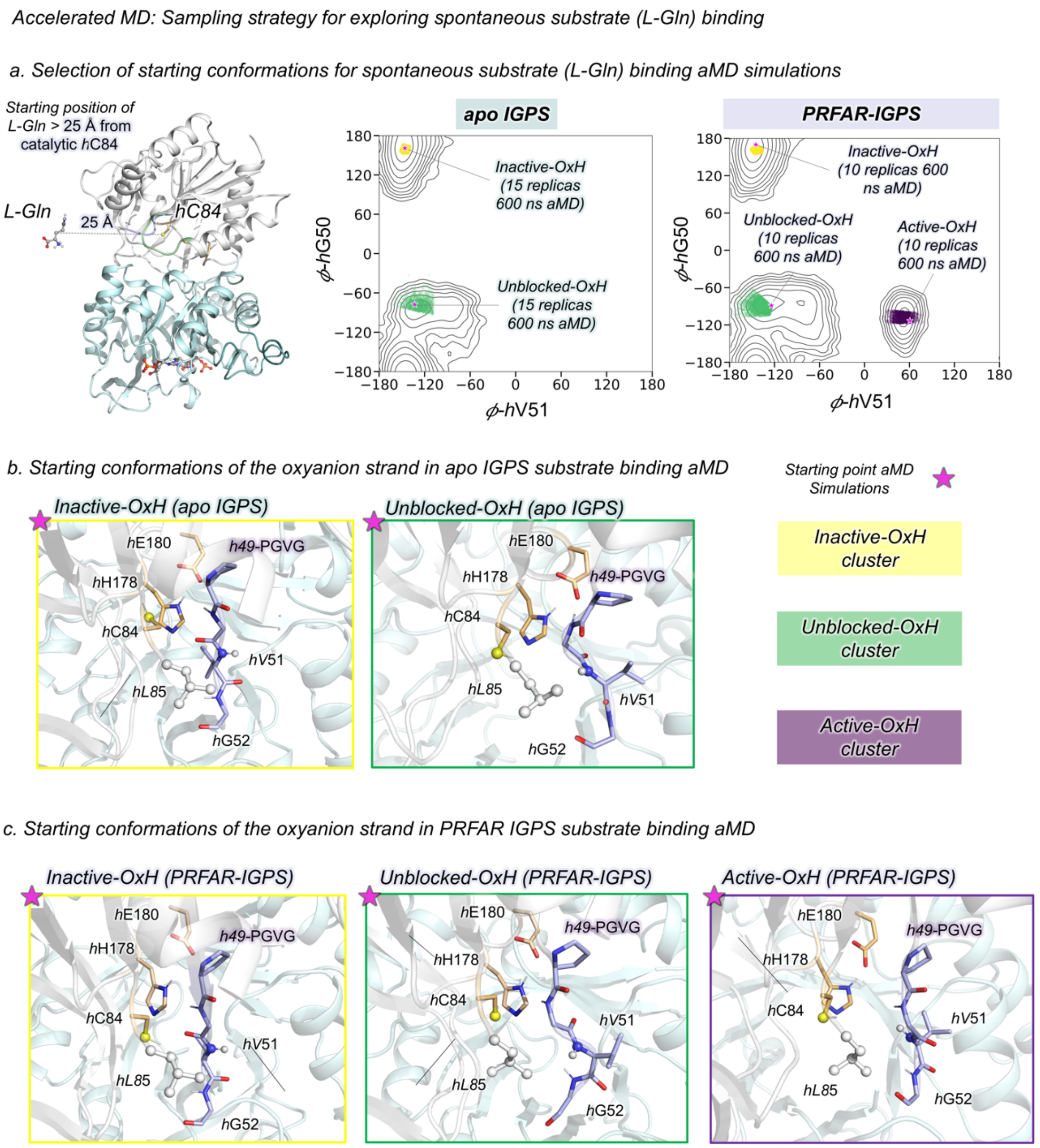


**Fig. S15. Spontaneous substrate binding sampling strategy**. (a) General scheme of spontaneous substrate binding process in the *apo* and PRFAR-IGPS states. The ligand (glutamine, L-Gln) is arbitrarily positioned 25 Å away from the catalytic *h*C84 residue at the HisH active site**.** Conformational landscape of apo and PRFAR-IGPS constructed using the $\phi$ dihedral angles of *h*V51 and *h*G50 with the relevant oxyanion strand conformations highlighted. The pink stars indicate the IGPS structures used as starting point for substrate binding aMD simulations. The cluster of conformations of Inactive-OxH, Unblocked-OxH, and Active-OxH are shown in yellow, green and purple, respectively. In apo-IGPS, 15 replicas of 600 ns are carried out starting from both the inactive-OxH and unblocked-OxH states. In PRFAR-IGPS, 10 replicas of 600 ns are performed starting from the inactive-OxH, unblocked-OxH, and active-OxH states. (b) Structures of IGPS with the corresponding HisH active site conformation selected as starting point for substrate binding aMD simulations in the apo state. (c) Structures of IGPS with the corresponding HisH active site conformation selected as starting point for substrate binding aMD simulations in the PRFAR-IGPS state.

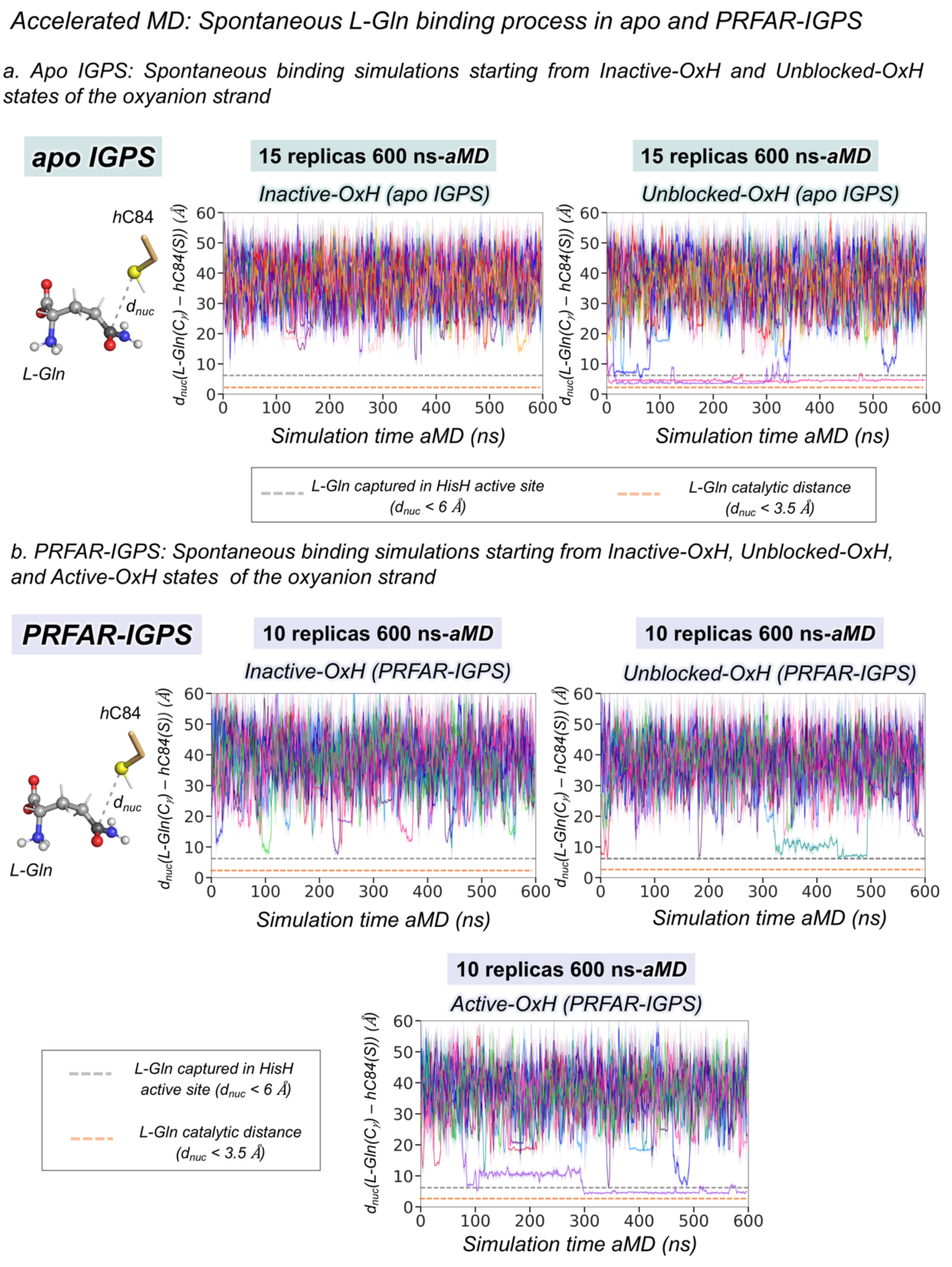


**Fig. S16. Spontaneous L-Gln binding process in apo and PRFAR-IGPS states**. (a) Plot of the distance ($d_{nuc}$) between the gamma carbon of L-Gln and the sulfur of *h*C84 for fifteen replicas of 600 ns aMD simulations starting from the Inactive-OxH and Unblocked-OxH states of the oxyanion strand in apo-IGPS. Each replica is depicted in different color. L-Gln binding occurs in 0/15 and 2/15 (magenta and purple) replicas of Inactive-OxH and Unblocked-OxH states, respectively. (b) Plot of the distance ($d_{nuc}$) between the gamma carbon of L-Gln and the sulfur of *h*C84 for ten replicas of 600 ns aMD simulations simulations starting from the Inactive-OxH, Unblocked-OxH, and Active-OxH states in PRFAR-IGPS. L-Gln binding occurs in 0/10, 0/10, and 1/10 (purple)

replicas of Inactive-OxH, Unblocked-OxH, and Active-OxH states, respectively. Horizontal gray dashed line indicate the distance when L-Gln is captured in the HisH active site ($d_{nuc}$ below 6 Å). Horizontal orange dashed line indicate the distance when L-Gln is at a catalytic distance of *h*C84 ($d_{nuc}$ below 3.5 Å). All distances are in Å.

### Accelerated MD: Analysis of Substrate Binding Simulations (continuation)

#### a. Analysis of productive L-Gln binding in the PRFAR-IGPS (starting from Active-OxH)

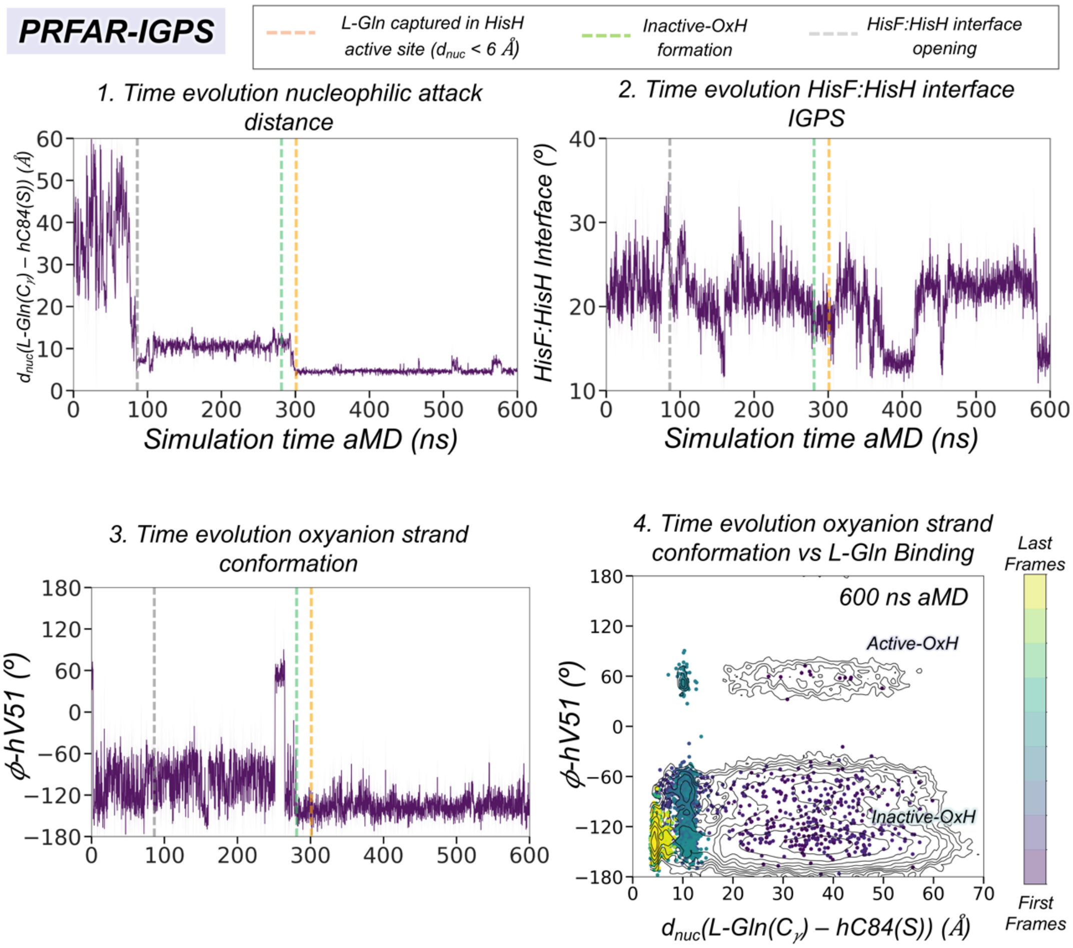


**Fig. S17. Analysis of substrate binding simulations in PRFAR-IGPS**. Analysis of a representative aMD simulation where L-Gln binding in the HisH active site is observed in PRFAR-IGPS. (a) Plot of the most significant distances for ligand binding aMD simulations in the PRFAR-IGPS. Vertical orange dashed line indicates when L-Gln is captured for the first time in the HisH active site. Vertical gray dashed line indicates when HisF:HisH interface expands to capture L-Gln. Vertical green dashed line indicates the deactivation of the oxyanion strand from Active-OxH to Inactive-OxH. 1. Plot of the nucleophilic attack distance between the amide carbon of L-Gln and the sulfur of the side chain of *h*C84. 2. Plot of the HisF:HisH interface angle along the simulation time. 3. Plot of the $\phi$ dihedral angle of *h*V51. 4. Projection of a representative aMD trajectory on the conformational landscape obtained from the nucleophilic attack distance between the thiol group of catalytic *h*C84 and the amide carbon of L-Gln, and the $\phi$ dihedral angle of *h*V51 (see Fig. 4 main text). The time evolution of the ligand binding pathway is represented in a color scale , purple for the first frames and yellow for the last frames.

### Molecular basis of L-Gln binding in PRFAR-IGPS

#### a. Ligand binding pathway in IGPS-PRFAR from different views

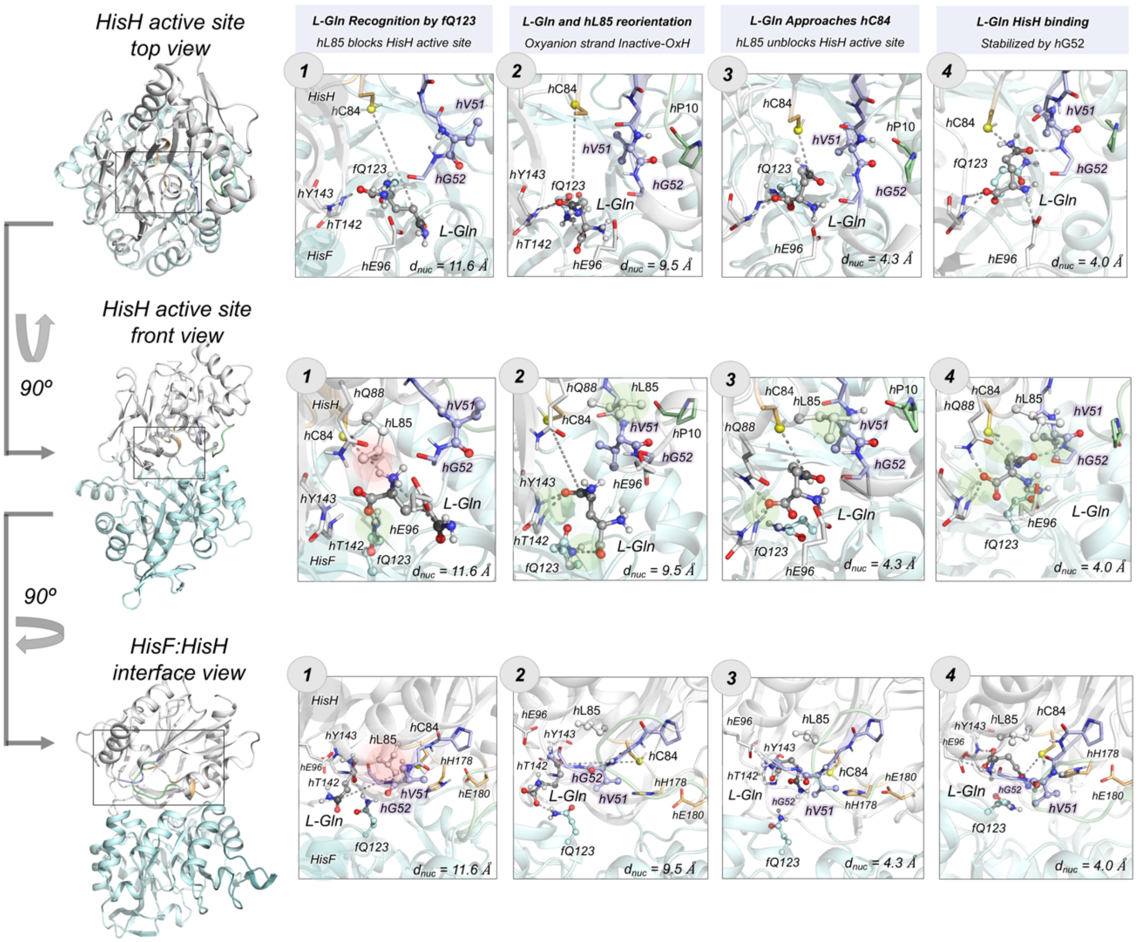


**Fig. S18. Molecular basis of L-Gln binding in PRFAR-IGPS**. (a) Molecular representation of the four most relevant steps in the binding pathway of L-Gln into the HisH active site obtained from aMD trajectories. The selected snapshots are depicted from different views: top, front and HisF:HisH interface views (see Fig. S6 for a complete description of the different points of view). The HisH catalytic residues are highlighted in orange, Ω-loop residues in green, and the residues of the *h*49-PGVG oxyanion strand in purple. Other relevant HisF and HisH residues are shown in cyan and white, respectively. The atoms of *h*L85, *f*Q123, *h*V51, and *h*P10 are shown as spheres. *h*L85 is not shown in the top view to facilitate the visual analysis of L-Gln conformation along the binding pathway. The green surfaces indicate the establishment of non-covalent interactions. The red surface indicates when *h*L85 blocks the access to the nucleophilic *h*C84. The nucleophilic attack distance ($d_{nuc}$) between the amide carbon of L-Gln and the sulfur of the side chain of *h*C84 is specified for each snapshot.

*Accelerated MD: Evolution of non-covalent interactions (NCI) along the L-Gln binding pathway*

*a. NCI and NCI volumes for the relevant steps of the binding pathway*

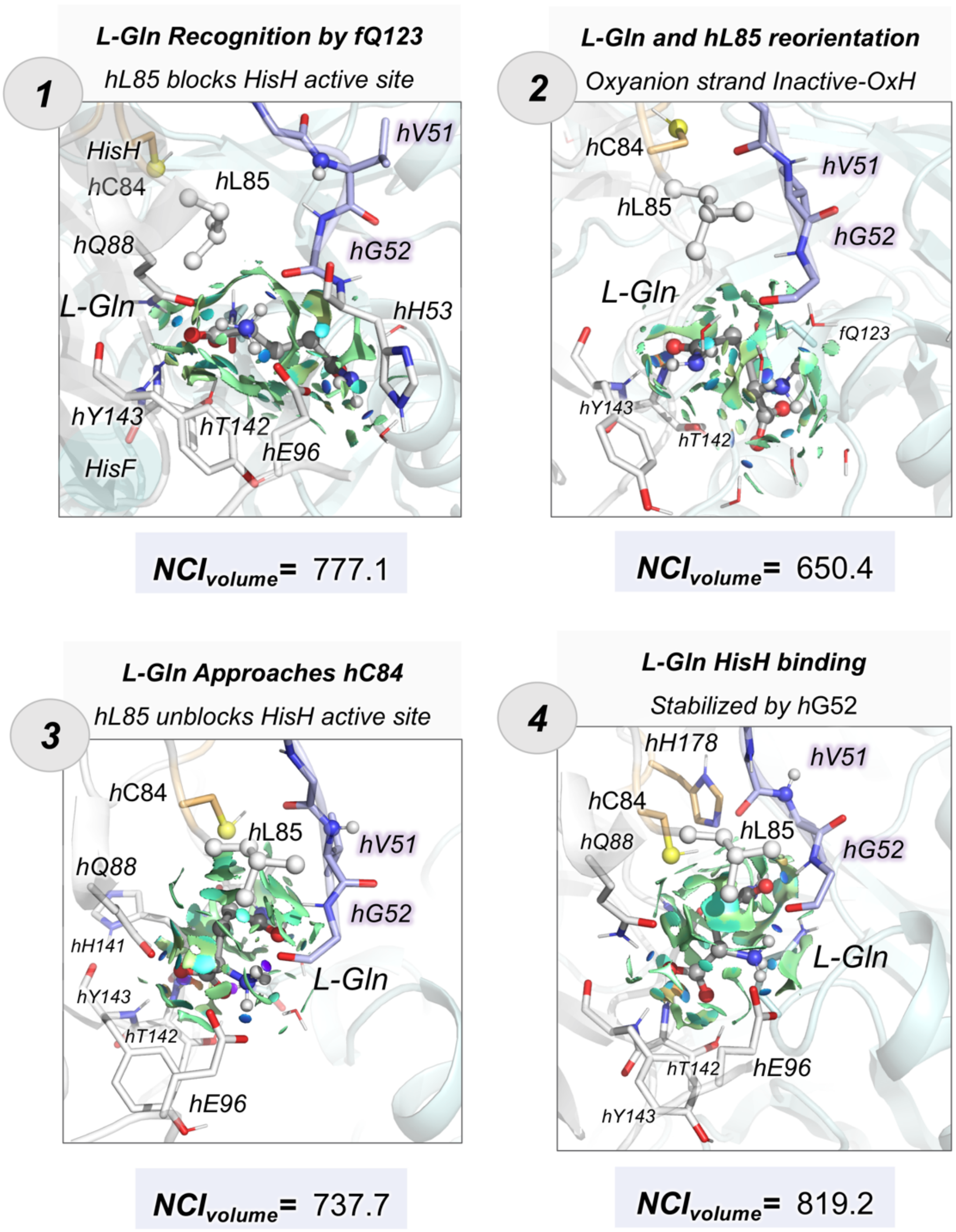


**Fig. S19. Evolution of non-covalent interactions (NCI) along the ligand binding pathway.** Schematic representation of non-covalent interactions for the four most relevant steps of the L-Gln binding process into the HisH active site in PRFAR-IGPS calculated with the NCI plot.(21) Blue, green, and red surfaces indicate strong, weak, and repulsive non-covalent interactions, respectively. The integrated volumes of non-covalent interactions are provided for each step. Higher volumes indicate overall stronger NCI interactions.

### Accelerated MD: Analysis of Substrate Binding Simulations

#### a. Analysis of productive L-Gln binding in the PRFAR-free IGPS (starting from Unblocked-OxH)

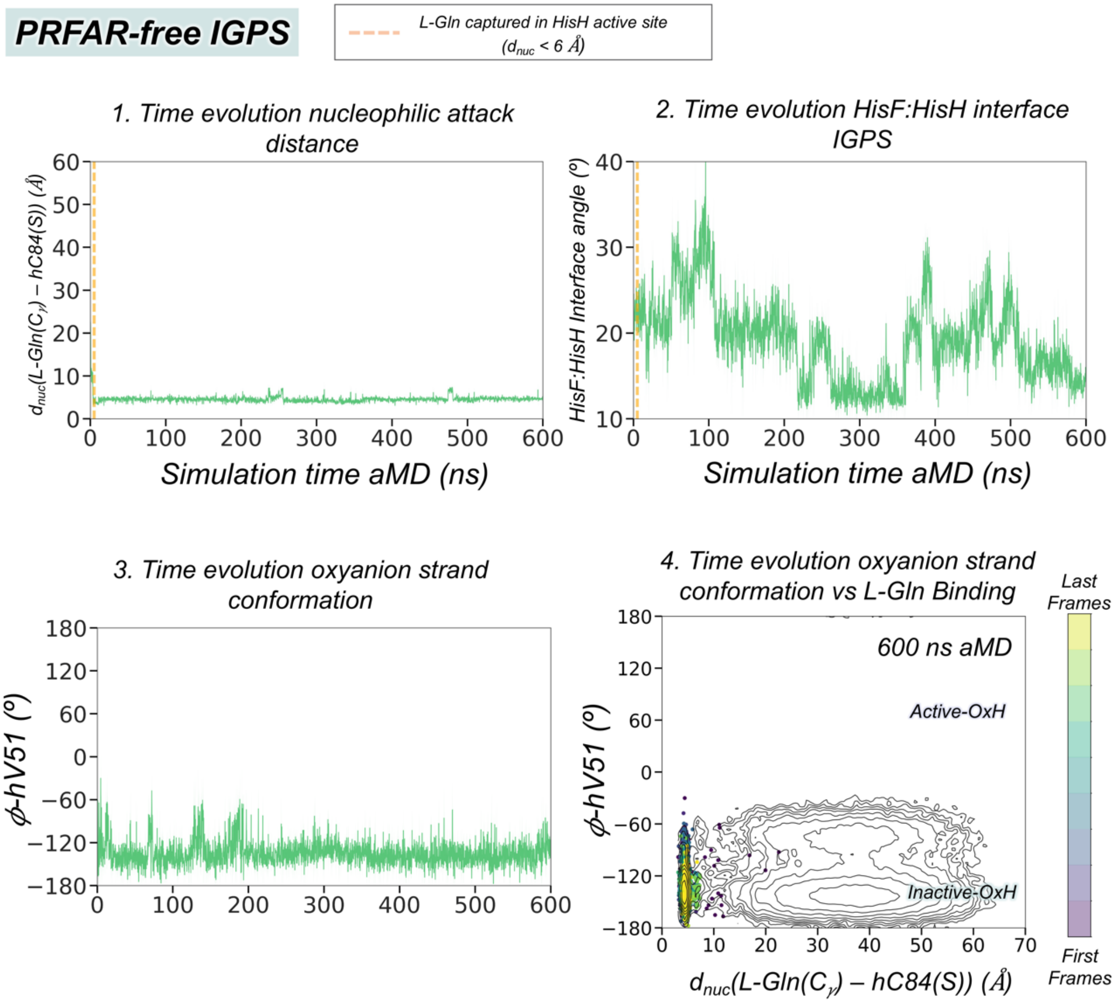


**Fig. S20. Analysis of substrate binding simulations in PRFAR-free IGPS**. Analysis of a representative aMD simulation where L-Gln binding in the HisH active site is observed in PRFAR-free IGPS. (a) Plot of the most significant distances for ligand binding aMD simulations in the PRFAR-free IGPS. Vertical orange dashed line indicates when L-Gln is captured for the first time in the HisH active site. 1. Plot of the nucleophilic attack distance between the amide carbon of L-Gln and the sulfur of the side chain of *h*C84. 2. Plot of the HisF:HisH interface angle along the simulation time. 3. Plot of the $\phi$ dihedral angle of *h*V51. 4. Projection of a representative aMD trajectory on the conformational landscape obtained from the nucleophilic attack distance between the thiol group of catalytic *h*C84 and the amide carbon of L-Gln, and the $\phi$ dihedral angle of *h*V51 (see Fig. 4 main text). The time evolution of the ligand binding pathway is represented in a color scale, as purple for the first frames and yellow for the last frames.

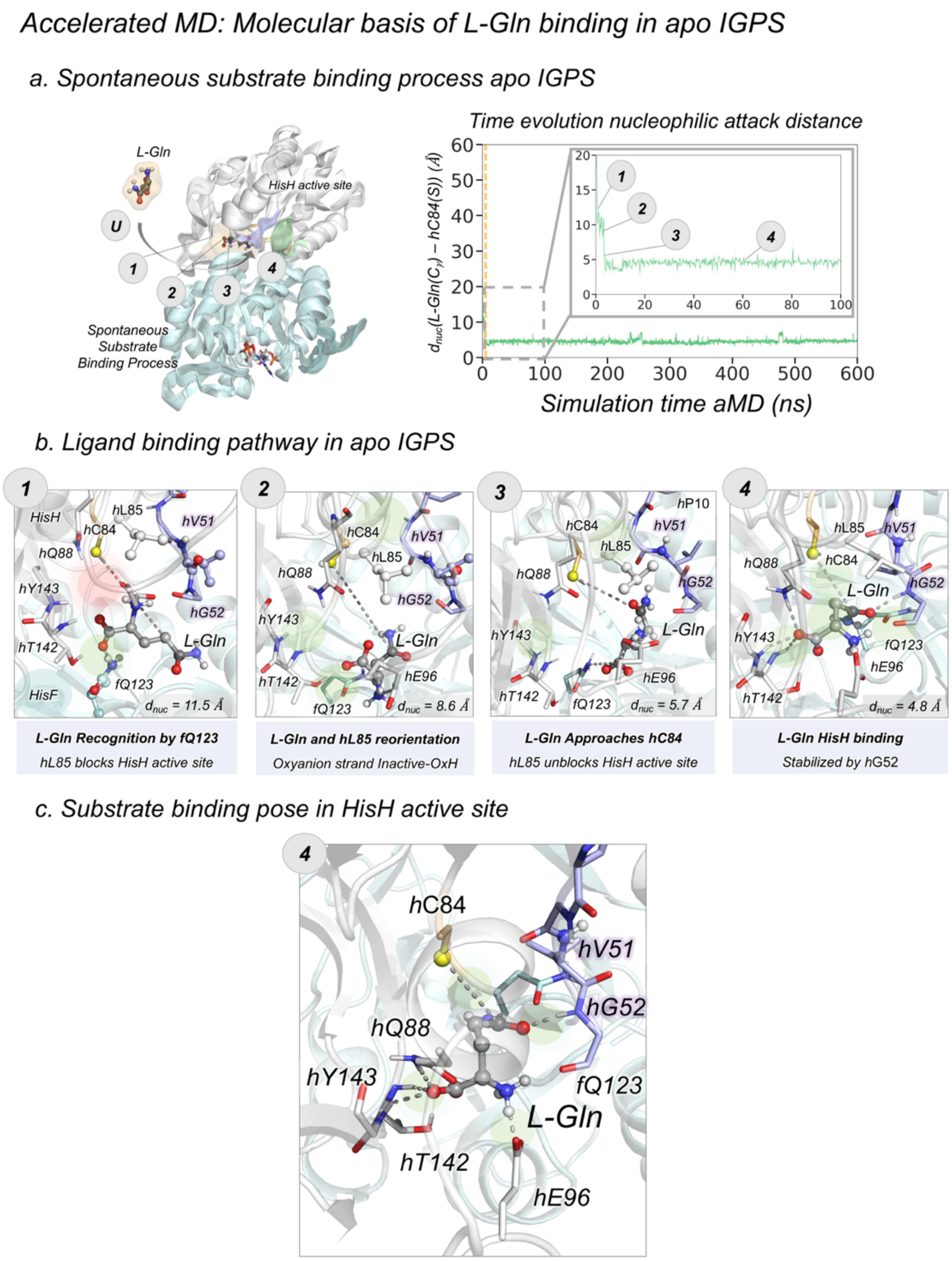


**Fig. S21. Molecular basis of L-Gln binding in PRFAR-free IGPS.** (a) General scheme of spontaneous substrate binding process in PRFAR-free IGPS. The numbers indicate the different steps of the substrate binding process. Plot of the distance corresponding to the nucleophilic attack along the 600 ns of aMD simulation for a representative replicas. (b) Structural representation of selected key conformational states of the L-Gln binding pathway in apo IGPS. The substrate is shown in gray, the oxyanion strand residues in purple, the catalytic residues in orange, the Ω-loop in green and other relevant HisH and HisF residues in white and cyan, respectively. (c) Structural representation of the L-Gln binding pose in the HisH active site of PRFAR-free IGPS.

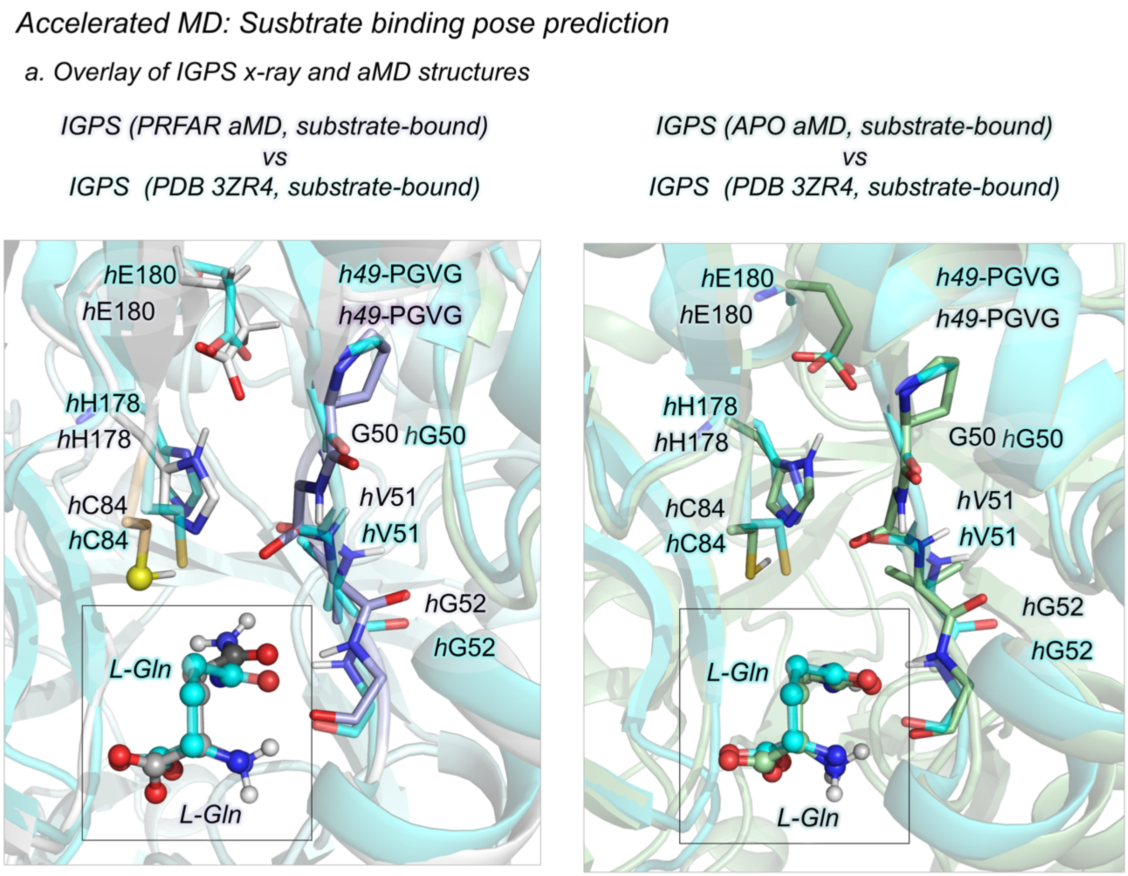


**Fig. S22. Substrate binding pose prediction**. (a) Overlay of a representative substrate-bound inactive-OxH PRFAR-IGPS (in purple) structure extracted from the aMD simulations with the substrate-bound IGPS x-ray structure (PDB: 3ZR4, in cyan). (b) Overlay of a representative substrate-bound inactive-OxH PRFAR-free IGPS (in green) structure extracted from the aMD simulations with the substrate-bound IGPS x-ray structure (PDB: 3ZR4, in cyan).

*Accelerated MD: HisH oxyanion strand conformational dynamics in the IGPS ternary complex*

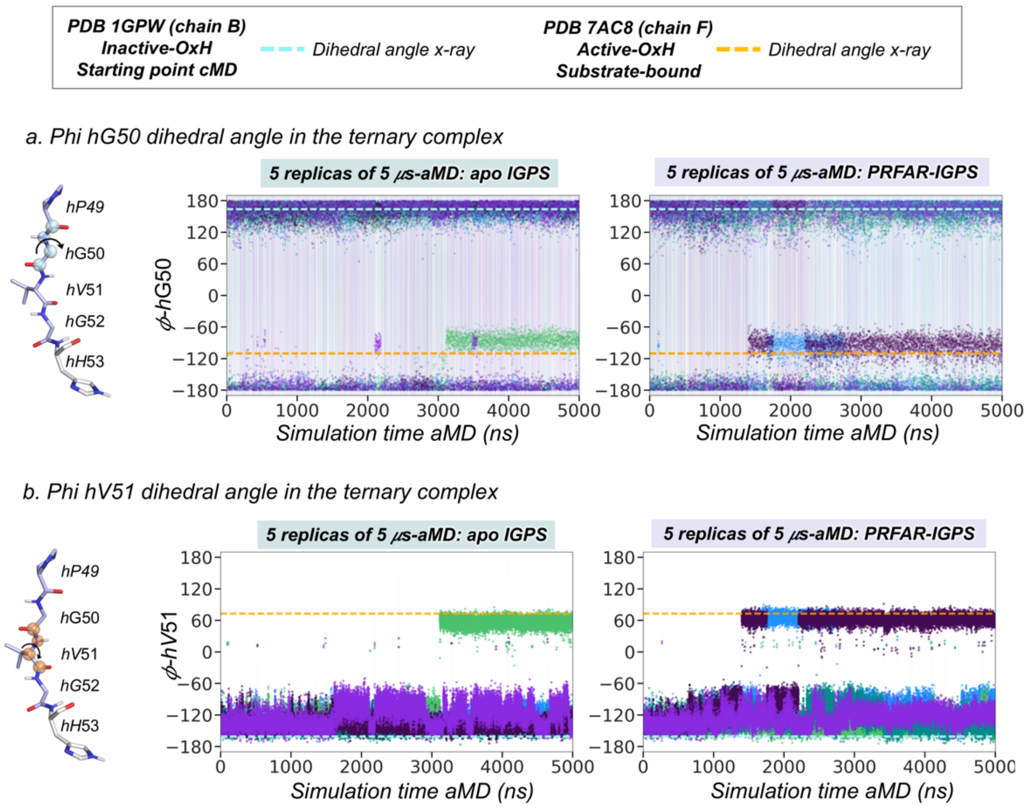


**Fig. S23. HisH *h*49-PVGV conformational dynamics in the IGPS ternary complex.** Plot of the most relevant dihedral angles of the *h*49-PGVG oxyanion strand for five replicas of 5 μs accelerated MD simulations in the PRFAR-free IGPS and PRFAR-IGPS states. Each replica is depicted in a different color. Horizontal cyan dashed lines indicate the dihedral angle found in the x-ray structure (1GPW chain B) used as starting point for cMD simulations. Horizontal orange dashed lines indicate the dihedral angle found in the x-ray structure of *h*C84A IGPS (7AC8 chain F) that displays an active conformation of the oxyanion strand. (a) ϕ dihedral angle of *h*G50; (b) ϕ dihedral angle of *h*V51. See Fig. S24 for a molecular representation of the most relevant states.

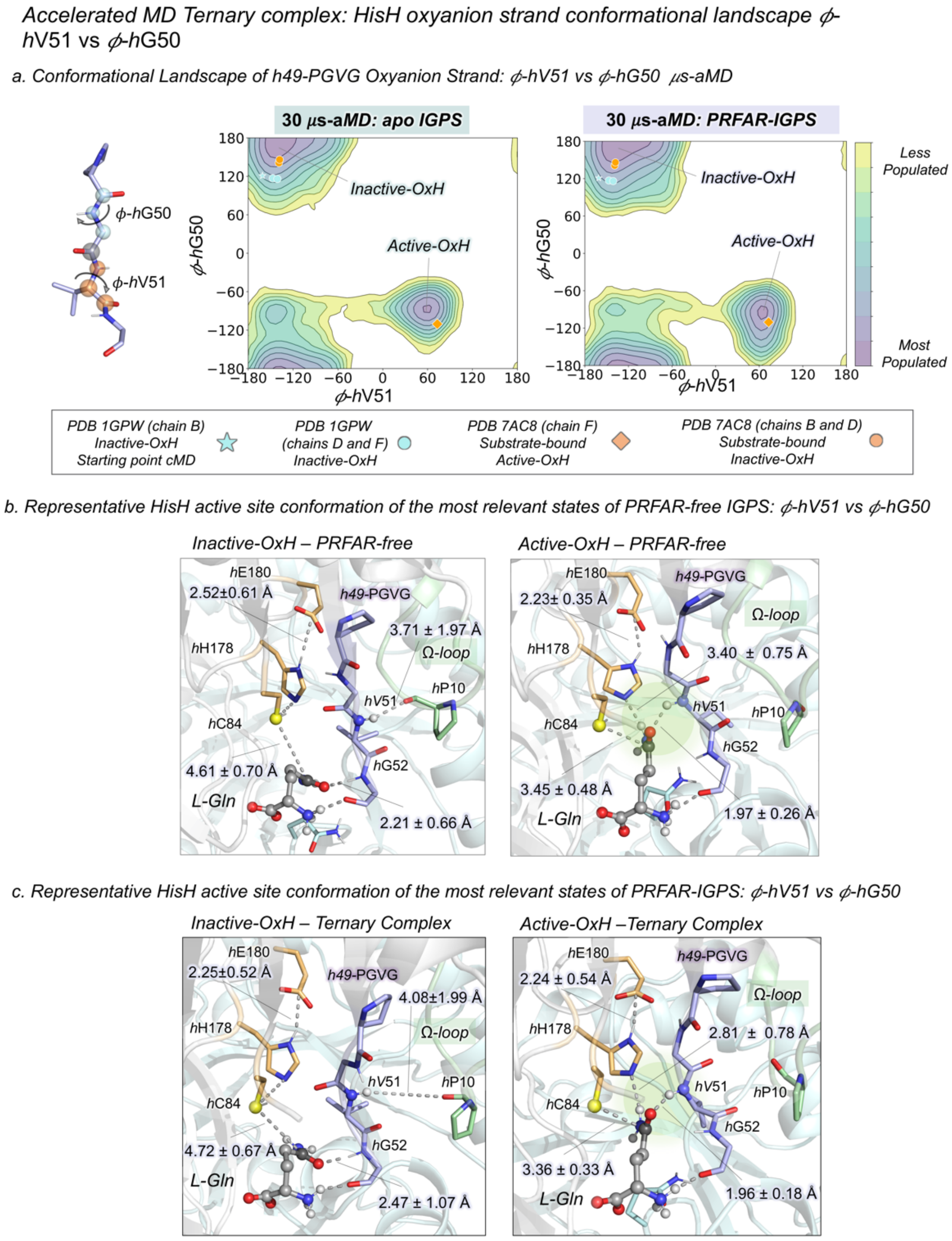


**Fig. S24. Conformational Landscape of *h*49-PGVG Oxyanion Strand in the ternary complex.** (a) The PRFAR-free IGPS and PRFAR-IGPS conformational landscapes are constructed from a total of 30 μs of aMD simulations in each case. (a) Conformational landscape of apo and PRFAR-IGPS constructed using the $\phi$ dihedral angles of *h*V51 and *h*G50. The values of the $\phi$ dihedral angles of *h*V51 and *h*G50 corresponding to the three chains of PDB 1GPW are depicted in cyan and the three chains of PDB 7AC8 are represented in orange. The conformation used as starting point for cMD simulations is shown using the star symbol. The conformation corresponding to the active oxyanion strand observed in *h*C84A IGPS is depicted using the diamond symbol. (b) Representative HisH active site structures of most populated states in PRFAR-free IGPS conformational landscape. (c) Representative HisH active site structures of most populated states in PRFAR-IGPS conformational landscape. The HisH catalytic residues are highlighted in orange,

Ω-loop residues in green, and the residues of the *h*49-PGVG oxyanion strand in purple. Other relevant HisF and HisH residues are shown in cyan and white respectively. The atoms of *L-Gln* are shown as spheres.

*Accelerated MD: Non-covalent interactions (NCI) in the HisH active site of the Ternary Complex*

*a. NCI and NCI volumes for the Active-OxH ternary complex*

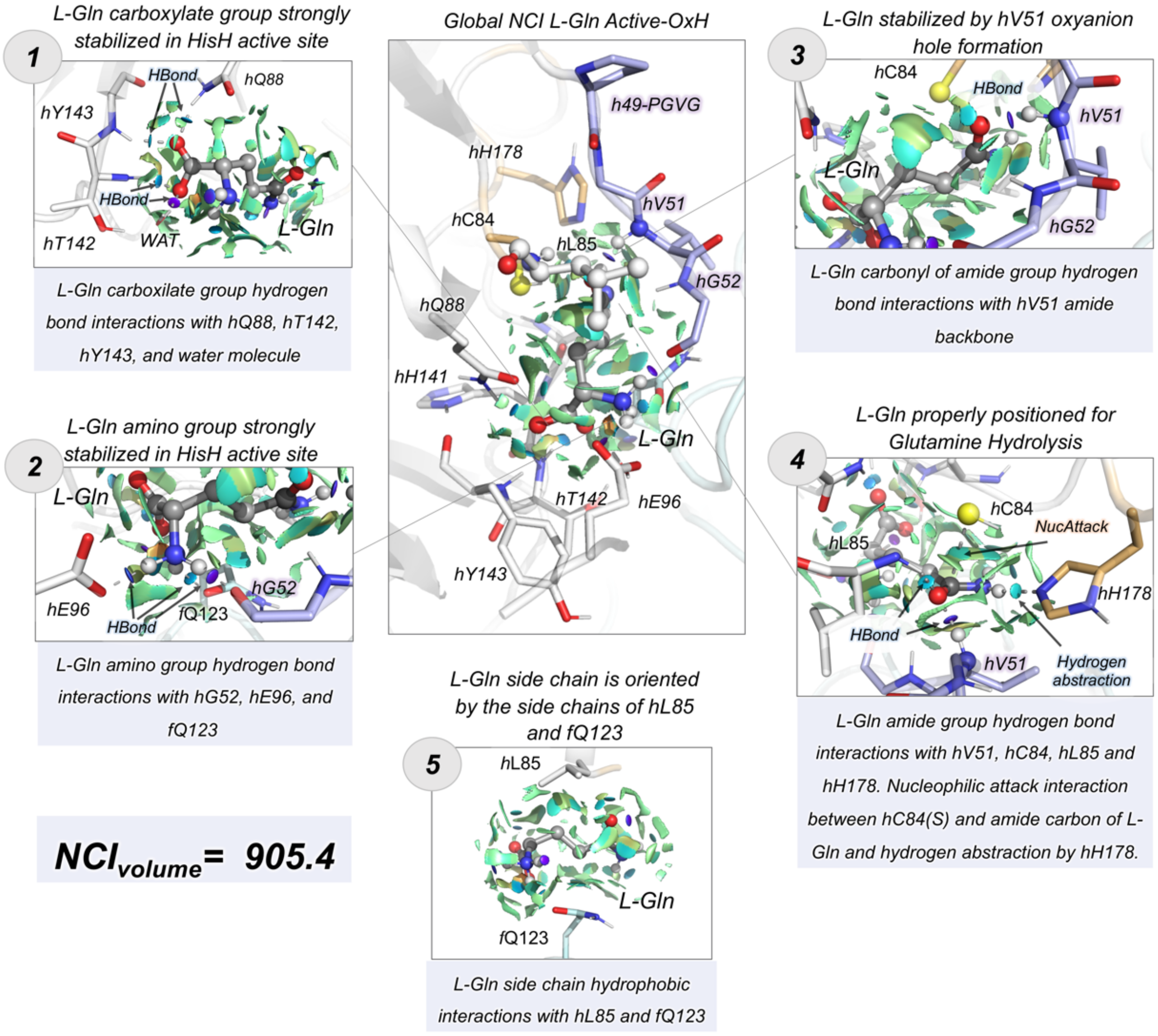

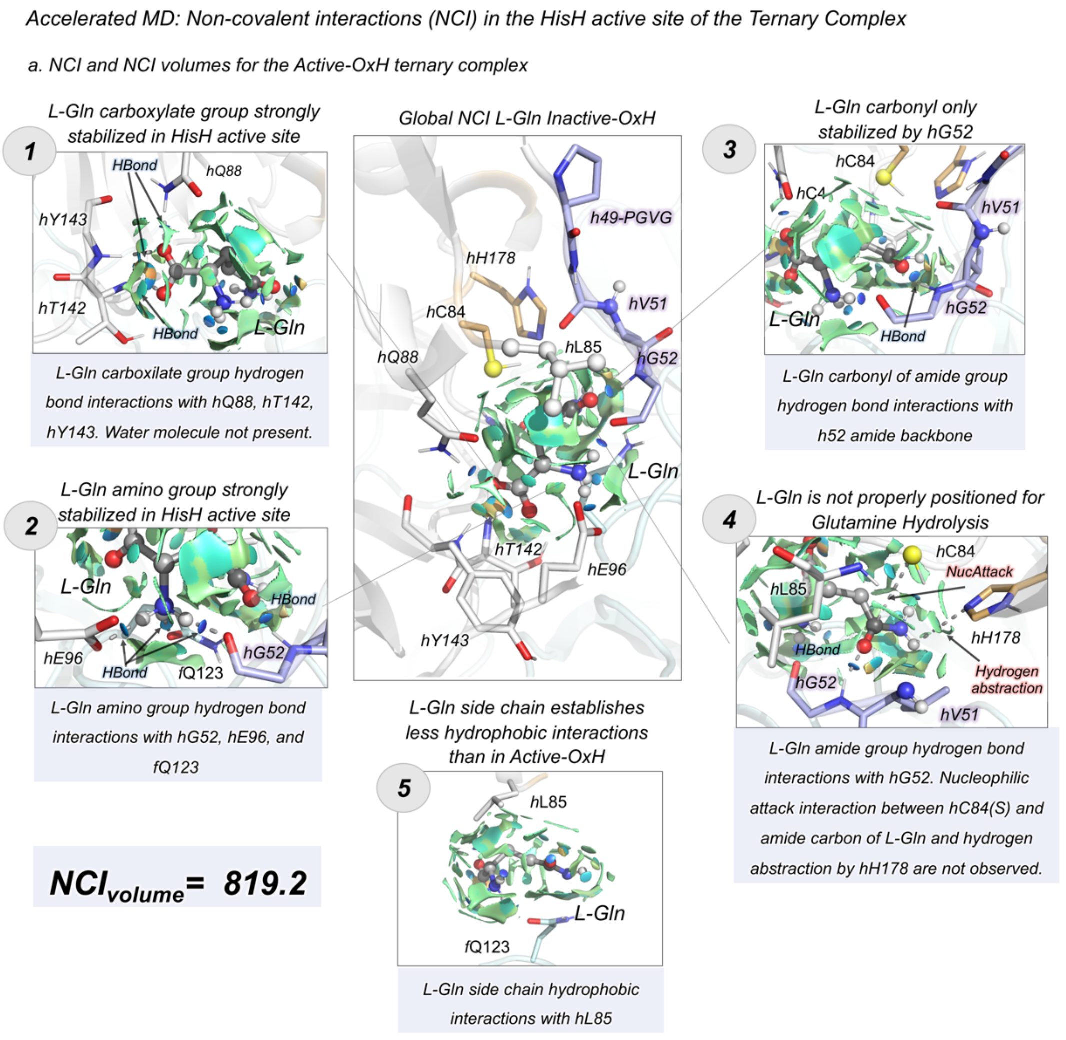


**Fig. S25. Non-covalent interactions (NCI) in the HisH active site of the IGPS Ternary Complex**. Schematic representation of non-covalent interactions for the L-Gln bound to Active-OxH (a) and Inactive-OxH (b) states of PRFAR-IGPS calculated with the NCI plot.(21) Blue, green, and red surfaces indicate strong, weak, and repulsive non-covalent interactions, respectively. The integrated volumes of non-covalent interactions are provided for each step. Higher volumes indicate stronger NCI interactions.

*Accelerated MD: HisH oxyanion strand conformational dynamics in the IGPS ternary complex*

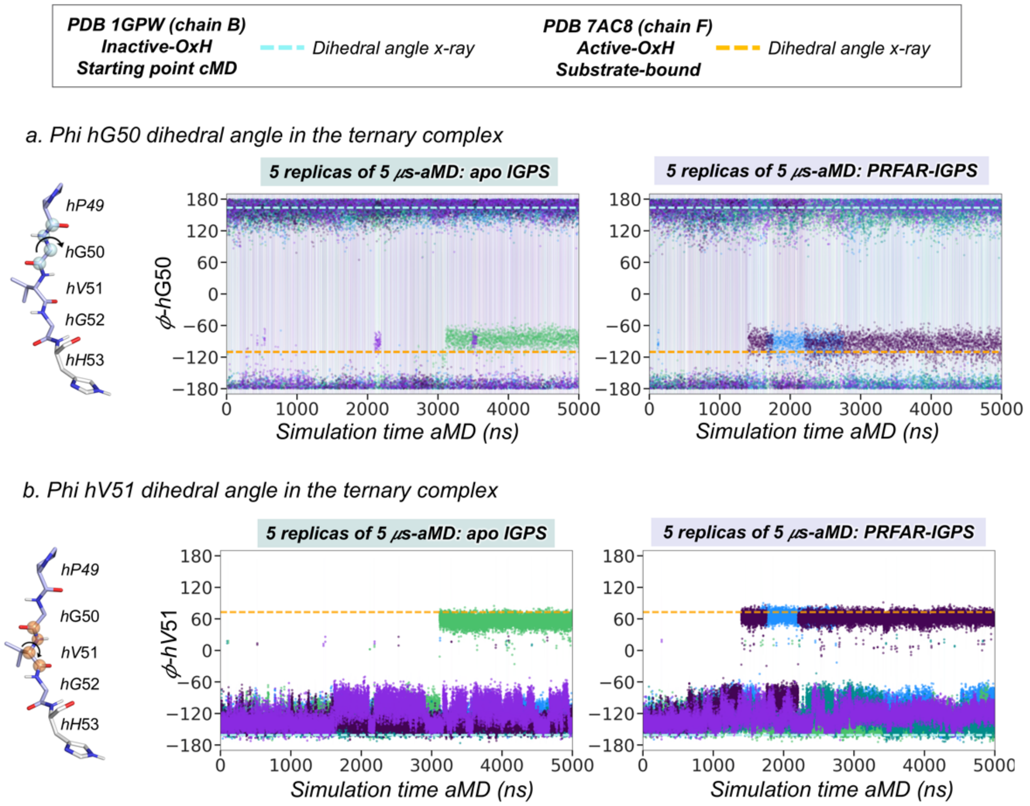


**Fig. S26. HisH *h*49-PVGV conformational dynamics in the IGPS ternary complex.** Plot of the nucleophilic attack distance (a) and HisF:HisH interface angle (b) for five replicas of 5 μs accelerated MD simulations in the PRFAR-free IGPS and PRFAR-IGPS states. Each replica is depicted in a different color. (a) Horizontal orange dashed line indicate that indicate nucleophilic attack distance at catalytic distance. (b) Horizontal black dashed lines indicate the HisF:HisH angle is below 12º indicative of productive interface closure. Horizontal orange dashed lines indicate the HisF:HisH interface angle found in the x-ray structure of *h*C84A IGPS (7AC8 chains E/F) that displays an active conformation of the oxyanion strand.

### *WT-Metadynamics: sampling strategy*

#### *a. aMD simulations and selected representative conformations for WT-Metadynamics*

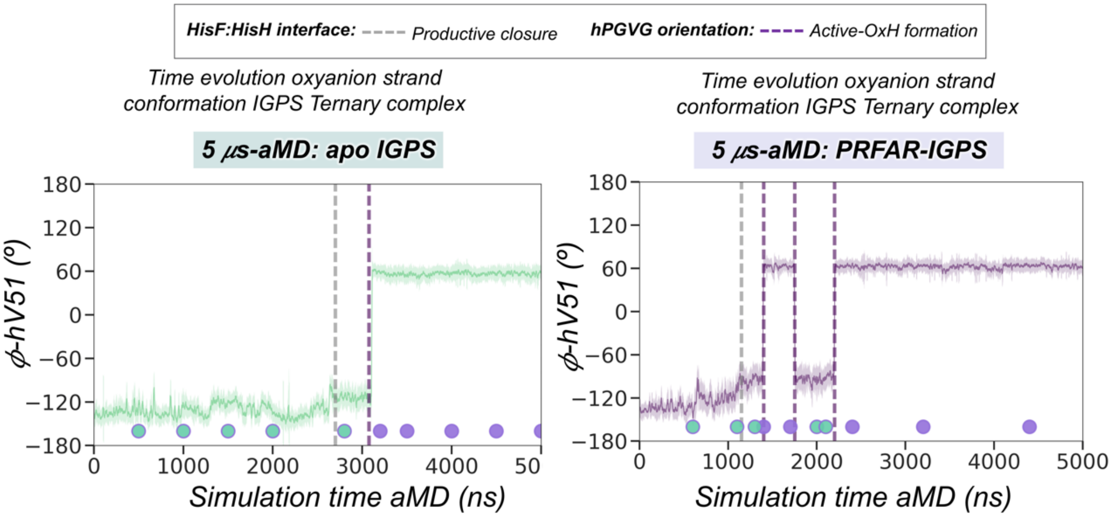


#### *b. Free Energy Landscape metadynamics simulations*

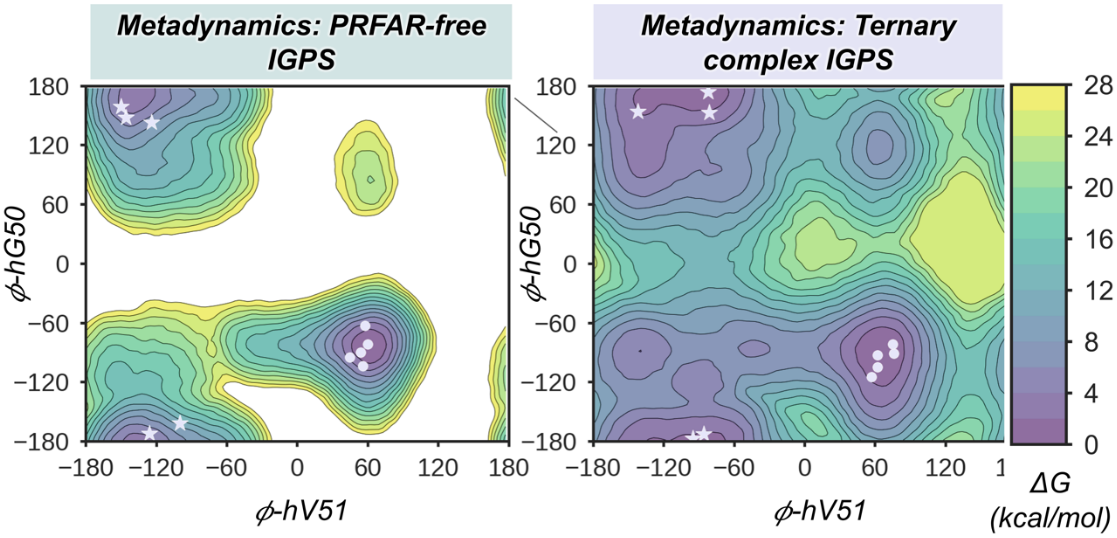


**Fig. S27. WT-Metadynamics sampling strategy**. (a) Plot of the *h*V51 dihedral angle along the 5 μs-aMD simulations in PRFAR-free (green) and PRFAR-IGPS (purple). The five representative aMD structures for the Inactive-OxH and Active-OxH conformations used starting points for the WT-metadynamics simulations are shown as green and purple circles respectively. (b) Free energy landscape of the *h*49-PGVG in the *apo* and PRFAR-IGPS states obtained from WT-tempered metadynamics simulations. Stars indicate the coordinates of the five starting points corresponding to Inactive-OxH walker replicas used for the WT-metadynamics while circles indicate the coordinates of the five starting points corresponding to the Active-OxH walker replicas. Note that the green and purple circles shown in (a) corresponds to the stars and circles in (b), respectively.

*Accelerated MD Ternary Complex: Active Ternary Complex prediction*

*a. Overlay of IGPS x-ray and aMD structures*

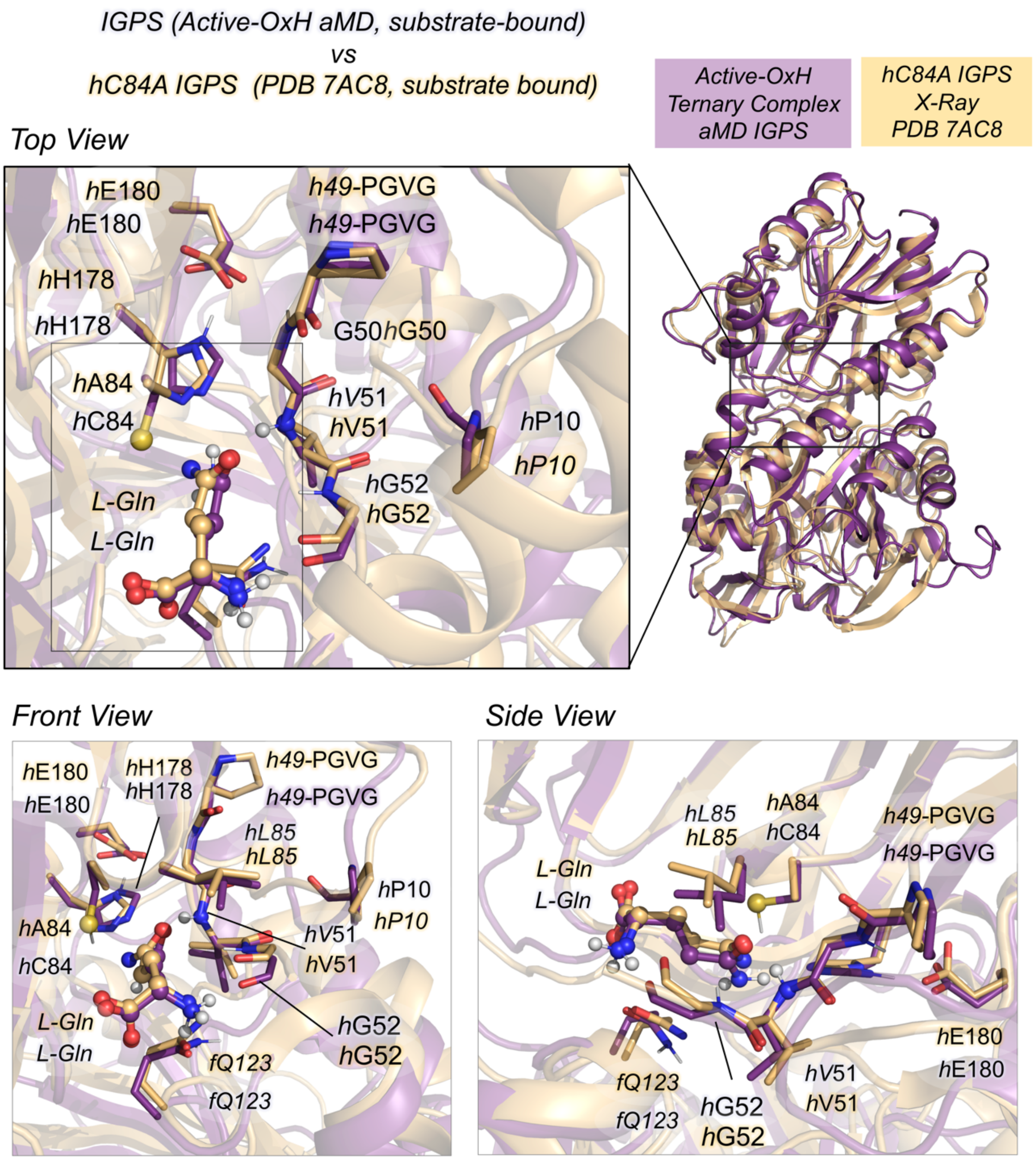


**Fig. S28. Active ternary complex predicted from aMD simulations**. (a) Overlay of a representative substrate-bound active-OxH PRFAR-IGPS (in purple) structure extracted from the PRFAR-IGPS conformational landscape with the substrate-bound *h*C84A IGPS (PDB: 7AC8 (chain F), in orange) from different views.

*Accelerated MD ternary complex: Analysis time evolution beyond 10 µs*

*a. aMD beyond 10 µs*

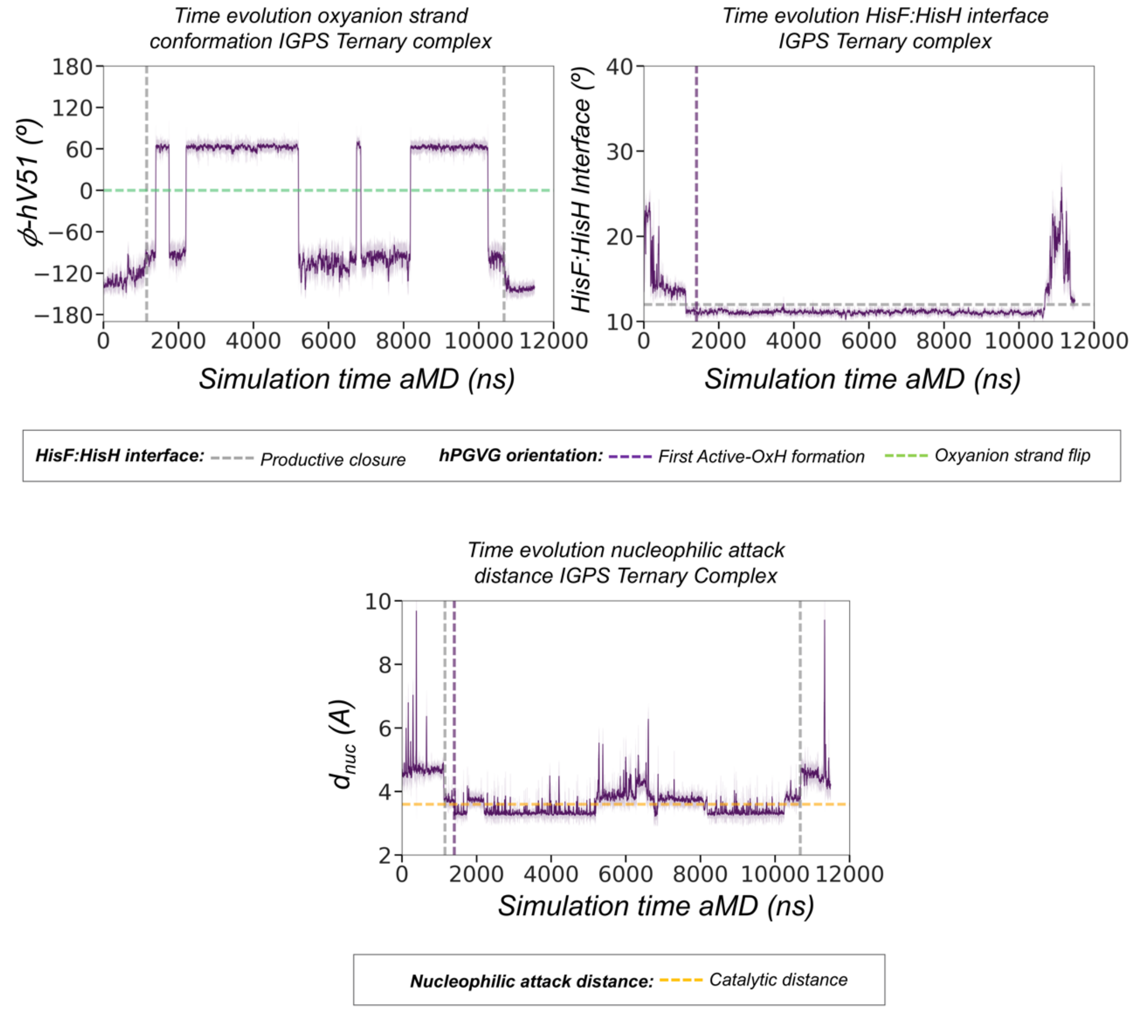


**Fig. S29. Ternary complex conformational dynamics beyond 10 µs.** (a) Plot of the HisF:HisH interface angle along 11.5 µs-aMD simulations. Plot of the *h*V51 dihedral angle along the 11.5 µs-aMD simulations. Plot of the distance corresponding to the nucleophilic attack along the 15 µs-aMD simulations. Gray dashed line indicates the range of HisF:HisH productive closure. Purple dashed line indicates the moment when the first *h*V51 oxyanion hole formation occurs. Green dashed line indicates the oxyanion hole transition. Orange dashed line indicate catalytic distance.

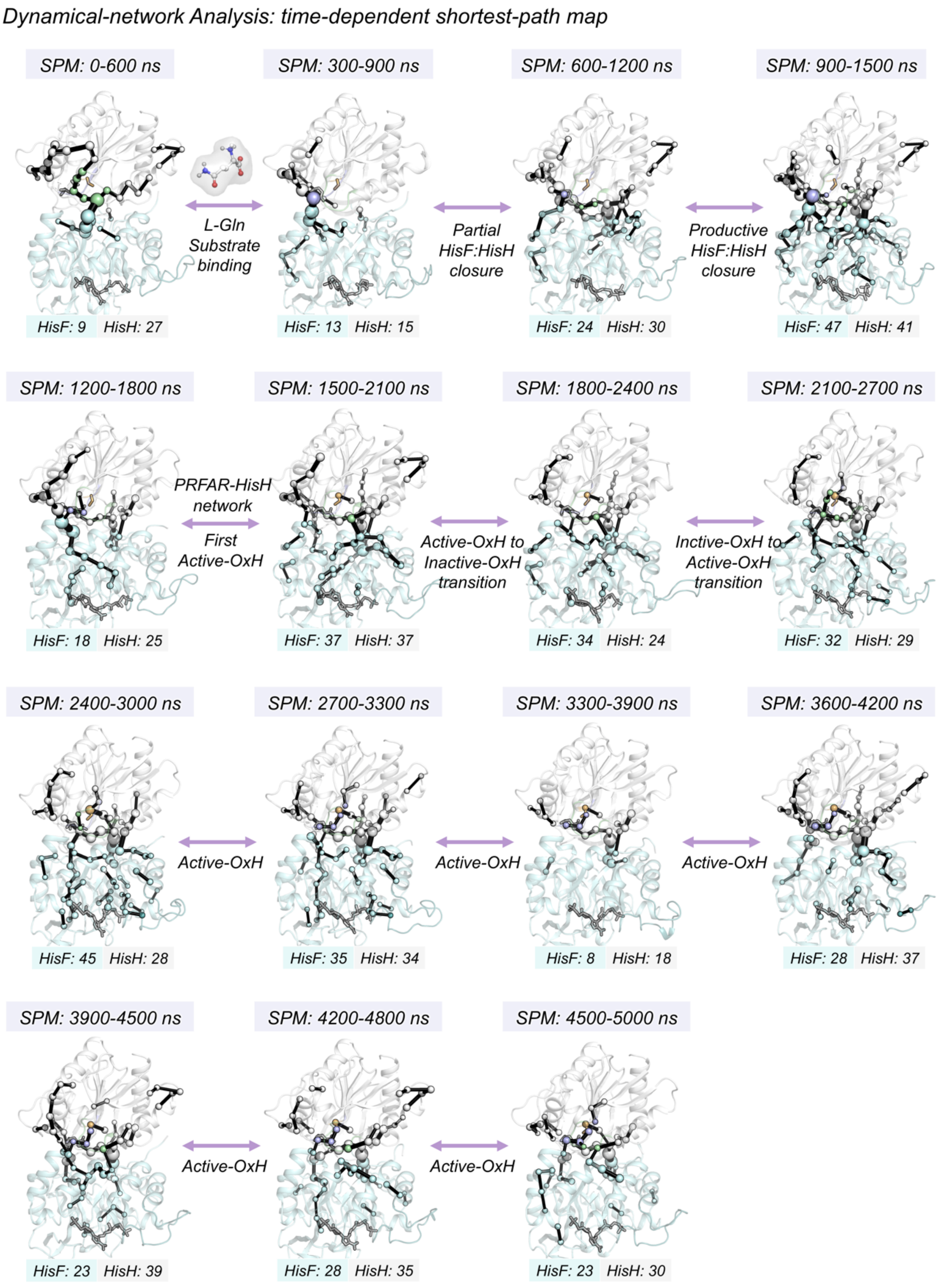


**Fig. S30. Dynamical Network Analysis: time-evolution Shortest-Path Map analysis**. Identification of the amino-acids that contribute to the propagation of the allosteric activation in IGPS.

*Dynamical Network Analysis: Shortest Path Map*

*a. SPM 0-600 ns*

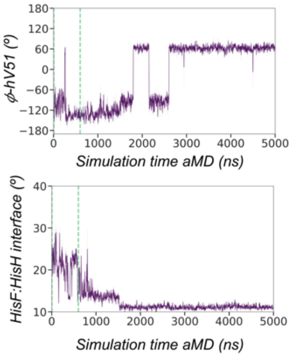


*L-Gln binding 400 ns aMD*

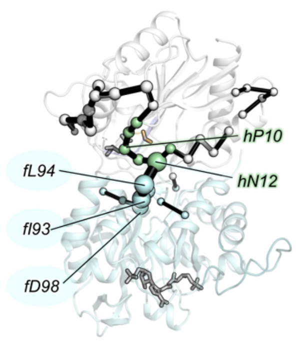


| ***HisF (9/253)*** | ***HisH (27/201)*** | | |
|---|---|---|---|
| *fA3* | *fV8* | *hV30* | *hW123* |
| *fV66* | *fG9* | *hS31* | |
| *fI93* | *fP10* | *hE36* | |
| *fL94* | *fG11* | *hS37* | |
| *fA97* | *fN12* | *hG55* | |
| *fD98* | *fI13* | *hE56* | |
| *fK99* | *fH14* | *hR59* | |
| *fS122* | *fY17* | *hR62* | |
| *fA124* | *fR18* | *hE63* | |
| | *fK21* | *hN64* | |
| | *fS24* | *hE95* | |
| | *fF27* | *hE96* | |
| | *fG28* | *hA97* | |

*b. SPM 300-900 ns*

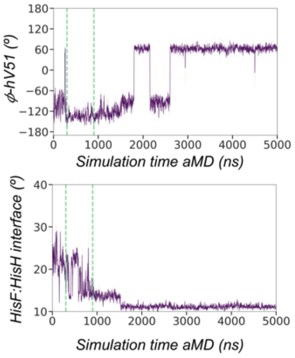


*Partial Closure 900 ns*

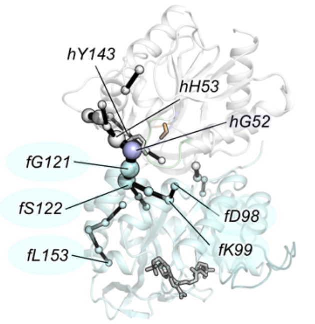


| ***HisF (15/253)*** | | ***HisH (14/201)*** | |
|---|---|---|---|
| *fA3* | *fV246* | *hG52* | *fY143* |
| *fD98* | *fN247* | *hH53* | |
| *fK99* | | *hG55* | |
| *fG121* | | *hE56* | |
| *fS122* | | *hR59* | |
| *fA124* | | *hL61* | |
| *fL153* | | *hL66* | |
| *fW156* | | *hE95* | |
| *fE159* | | *hA97* | |
| *fV160* | | *fV111* | |
| *fA165* | | *fW123* | |
| *fG166* | | *fV140* | |
| *fE167* | | *fV141* | |

*c. SPM 600-1200 ns*

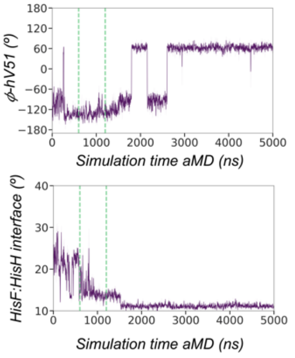


*Partial Closure 900 ns*

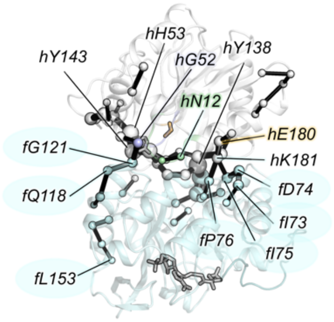


| ***HisF (24/253)*** | | ***HisH (34/201)*** | | |
|---|---|---|---|---|
| *fA3* | *fL153* | *fG11* | *fL61* | *fV140* |
| *fK4* | *fW156* | *fN12* | *fL66* | *fH141* |
| *fR5* | *fE159* | *fG19* | *fE95* | *fT142* |
| *fD45* | *fV160* | *fA23* | *fA97* | *fY143* |
| *fV69* | *fR163* | *fS24* | *fV110* | *fE180* |
| *fI73* | *fT195* | *fF27* | *fV111* | *fK181* |
| *fD74* | *fI199* | *fE28* | *fR117* | *fS182* |
| *fI75* | *fD219* | *fV30* | *fH120* | *fG186* |
| *fP76* | *fA220* | *fG52* | *fM121* | |
| *fF77* | *fV246* | *fH53* | *fG122* | |
| *fT114* | *fN247* | *fG55* | *fW123* | |
| *fQ118* | | *fE56* | *fY138* | |
| *fG121* | | *fR59* | *fF139* | |

Dynamical Network Analysis: Shortest Path Map

d. SPM 900-1500 ns

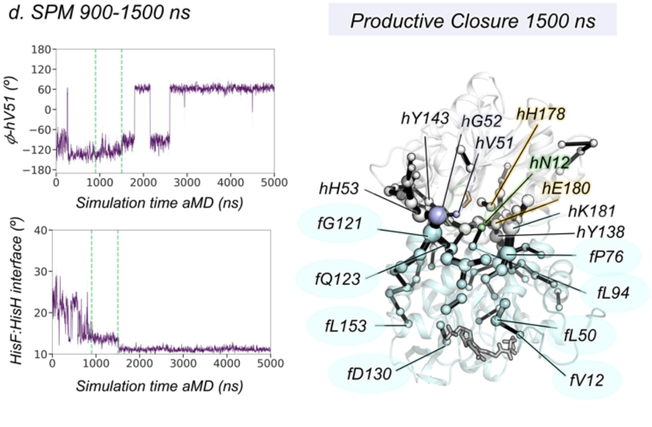


| HisF (48/253) | | | HisH (41/201) | | |
|---|---|---|---|---|---|
| fA3 | fA97 | fR163 | hN12 | hE95 | hV140 |
| fK4 | fK99 | fL193 | hS24 | hA97 | hH141 |
| fR5 | fV100 | fT194 | hF27 | hV111 | hY143 |
| fV12 | fS101 | fT195 | hE28 | hK112 | hF177 |
| fY39 | fI102 | fL196 | hV30 | hL113 | hH178 |
| fI44 | fT114 | fS201 | hE36 | hS115 | hE180 |
| fD45 | fA117 | fA218 | hS37 | hR117 | hK181 |
| fE46 | fG121 | fD219 | hV51 | hW123 | hS182 |
| fF49 | fQ123 | fA220 | hG52 | hN124 | hG186 |
| fL50 | fA124 | fA221 | hH53 | hE125 | |
| fI73 | fI129 | fL222 | hG55 | hV126 | |
| fI75 | fD130 | fD233 | hE56 | hI127 | |
| fP76 | fL153 | fE236 | hM58 | hF128 | |
| fF77 | fW156 | fL241 | hR59 | hY137 | |
| fT78 | fE159 | fV246 | hR62 | hY138 | |
| fL94 | fV160 | fN247 | hE63 | hF139 | |

e. SPM 1200-1800 ns

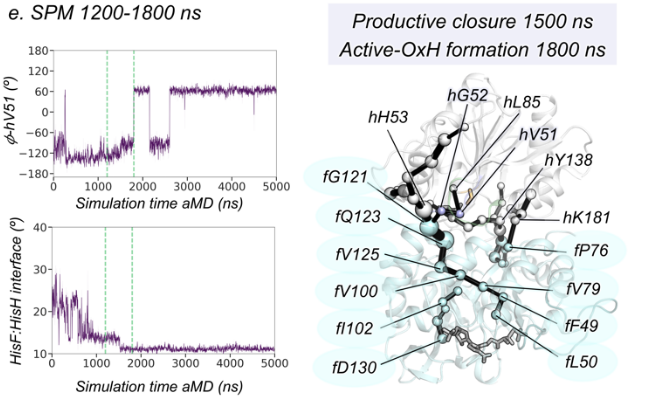


| HisF (18/253) | | HisH (25/201) | |
|---|---|---|---|
| fA3 | fA128 | hP38 | hW123 |
| fK4 | fI129 | hV51 | hN124 |
| fR5 | fD130 | hG52 | hE125 |
| fF49 | fV246 | hH53 | hV126 |
| fL50 | fN247 | hG55 | hY137 |
| fP76 | | hE56 | hY138 |
| fV79 | | hR59 | hF139 |
| fV100 | | hL61 | hH141 |
| fS101 | | hL66 | hY143 |
| fI102 | | hL85 | hK181 |
| fG121 | | hE95 | hS182 |
| fQ123 | | hA97 | hG186 |
| fV125 | | hV111 | |

f. SPM 1500-2100 ns

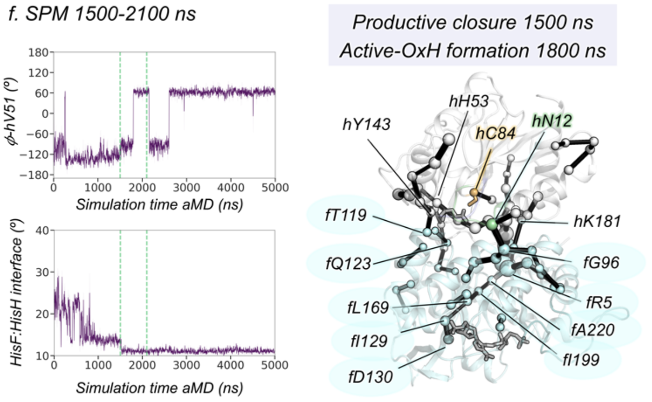


| HisF (37/253) | | | HisH (37/201) | | |
|---|---|---|---|---|---|
| fA3 | fG96 | fW156 | hN12 | hC84 | hF128 |
| fK4 | fA97 | fE159 | hM14 | hE95 | hY137 |
| fR5 | fK99 | fV160 | hS24 | hA97 | hY138 |
| fV33 | fV100 | fR163 | hF27 | hV111 | hF139 |
| fG36 | fS101 | fE167 | hE28 | hK112 | hV140 |
| fY39 | fI102 | fL169 | hV30 | hL113 | hH141 |
| fI44 | fT114 | fL196 | hS31 | hS115 | hY143 |
| fD45 | fA117 | fI199 | hH53 | hR117 | hF177 |
| fL50 | fT119 | fA220 | hG55 | hW123 | hK181 |
| fD51 | fG121 | fV246 | hE56 | hN124 | hS182 |
| fV69 | fQ123 | fN247 | hR59 | hE125 | hG186 |
| fA70 | fI129 | | hL61 | hV126 | |
| fP76 | fD130 | | hL66 | hI127 | |

*Dynamical Network Analysis: Shortest Path Map*

*g. SPM 1800-2400 ns*

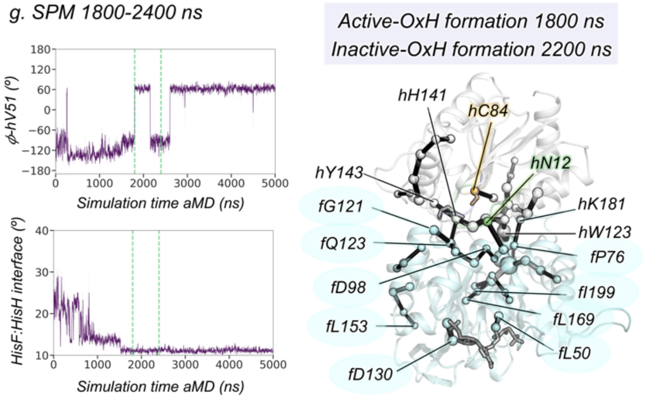


| HisF (34/253) | | | HisH (24/201) | |
|---|---|---|---|---|
| fA3 | fA97 | fV160 | hN12 | hI127 |
| fK4 | fD98 | fR163 | hG55 | hF128 |
| fR5 | fK99 | fL169 | hE56 | hY137 |
| fY39 | fT114 | fL199 | hR59 | hY138 |
| fI44 | fA117 | fA218 | hR62 | hF139 |
| fD45 | fG121 | fA220 | hD65 | hV140 |
| fV48 | fQ123 | fV246 | hE66 | hH141 |
| fF49 | fA124 | fN247 | hC84 | hY143 |
| fL50 | fI129 | | hA97 | hK181 |
| fD51 | fD130 | | hW123 | hS182 |
| fP76 | fL153 | | hN124 | hG186 |
| fT78 | fW156 | | hE125 | |
| fG96 | fE159 | | hV126 | |

*h. SPM 2100-2700 ns*

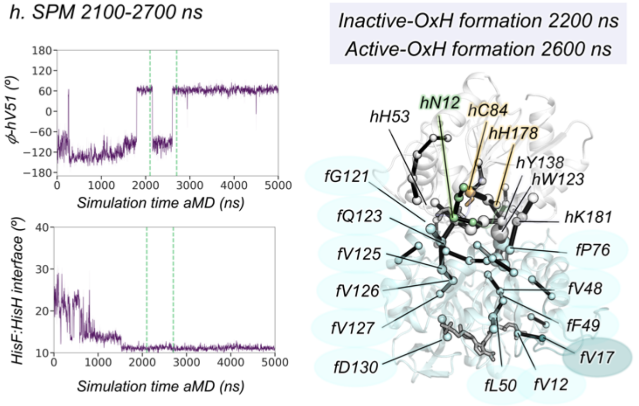


| HisF (32/253) | | | HisH (29/201) | | |
|---|---|---|---|---|---|
| fA3 | fT78 | fI129 | hV8 | hR117 | hK181 |
| fK4 | fL94 | fD130 | hG9 | hW123 | hS182 |
| fR5 | fR95 | fA224 | hP10 | hN124 | hG186 |
| fV12 | fA97 | fV226 | hG11 | hF125 | |
| fV17 | fK99 | fV246 | hE56 | hV126 | |
| fV33 | fT114 | fN247 | hR59 | hY137 | |
| fV48 | fA117 | | hR62 | hY138 | |
| fF49 | fG121 | | hD65 | hF139 | |
| fL50 | fQ123 | | hL66 | hV140 | |
| fD51 | fA124 | | hG82 | hH141 | |
| fE67 | fV125 | | hV83 | hY143 | |
| fV69 | fV126 | | hC84 | hF177 | |
| fP76 | fV127 | | hS115 | hH178 | |

*i. SPM 2400-3000 ns*

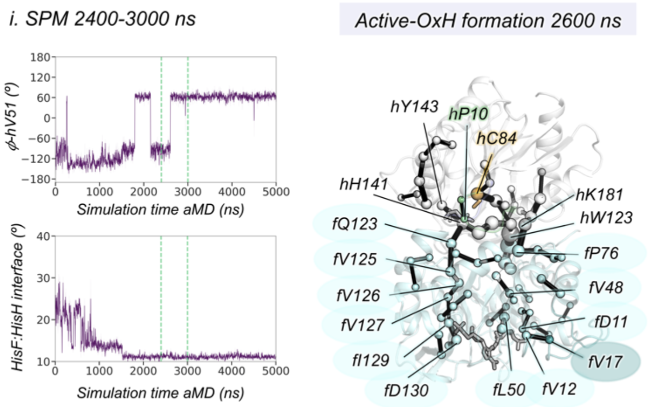


| HisF (45/253) | | | HisH (28/201) | |
|---|---|---|---|---|
| fA3 | fQ72 | fV125 | hG9 | hY137 |
| fK4 | fI73 | fV126 | hP10 | hY138 |
| fR5 | fI75 | fV127 | hG55 | hF139 |
| fI7 | fP76 | fI129 | hE56 | hV140 |
| fA8 | fT78 | fD130 | hR59 | hH141 |
| fC9 | fA97 | fA131 | hR62 | hY143 |
| fL10 | fK99 | fT142 | hD65 | hF177 |
| fD11 | fS101 | fS201 | hL66 | hH178 |
| fV12 | fI102 | fL222 | hG82 | hK181 |
| fV17 | fN103 | fA224 | hV83 | hS182 |
| fV33 | fT104 | fV226 | hC84 | hG186 |
| fV48 | fL112 | fI232 | hA97 | hL189 |
| fF49 | fQ115 | fD233 | hW123 | |
| fL50 | fI116 | | hN124 | |
| fD41 | fQ123 | | hE125 | |
| fV69 | fA124 | | hV126 | |

*Dynamical Network Analysis: Shortest Path Map*

*j. SPM 2700-3300 ns*

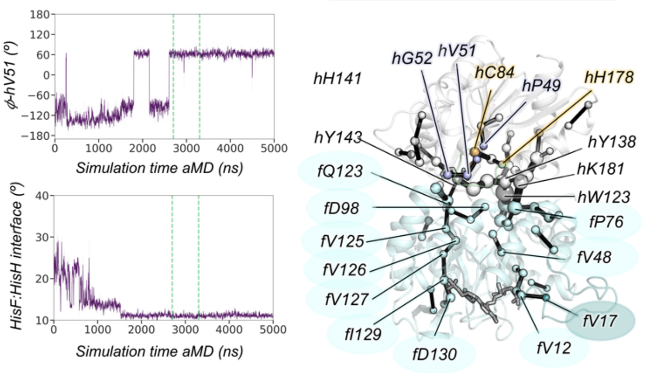


| HisF (36/253) | | | HisH (35/201) | | |
|---|---|---|---|---|---|
| fA3 | fI75 | fI129 | hS24 | hA97 | hH141 |
| fK4 | fP76 | fD130 | hF27 | hV111 | hY143 |
| fR5 | fF77 | fA131 | hF47 | hS115 | hF177 |
| fC9 | fT78 | fT142 | hI48 | hR117 | hH178 |
| fL10 | fD98 | fT194 | hP49 | hW123 | hK181 |
| fD11 | fK99 | fT195 | hG50 | hN124 | hS182 |
| fV12 | fT104 | fA224 | hV51 | hE125 | hG186 |
| fV17 | fG121 | fV226 | hG52 | hV126 | hR187 |
| fV33 | fQ123 | fV246 | hG55 | hI127 | hL189 |
| fV48 | fA124 | fN247 | hE56 | hY137 | |
| fV69 | fV125 | | hR59 | hY138 | |
| fQ72 | fV126 | | hR62 | hF139 | |
| fI73 | fV127 | | hC84 | hV140 | |

*k. SPM 3300-3900 ns*

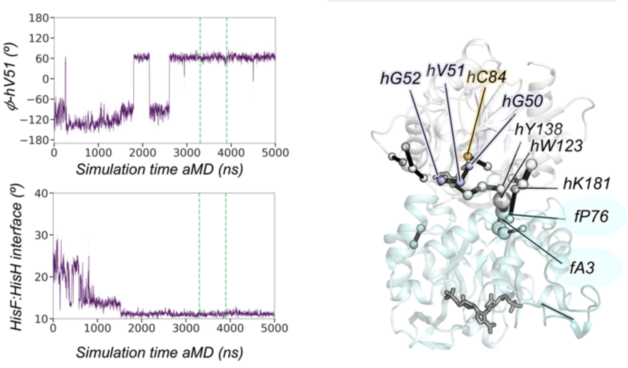


| HisF (8/253) | HisH (19/201) | |
|---|---|---|
| fA3 | hG50 | hV140 |
| fK4 | hV51 | hH141 |
| fR5 | hG52 | hF177 |
| fD45 | hH53 | hK181 |
| fP76 | hG55 | hS182 |
| fT194 | hE56 | hG186 |
| fT195 | hR59 | |
| fN247 | hC84 | |
| | hA97 | |
| | hW123 | |
| | hY137 | |
| | hY138 | |
| | hF139 | |

*l. SPM 3600-4200 ns*

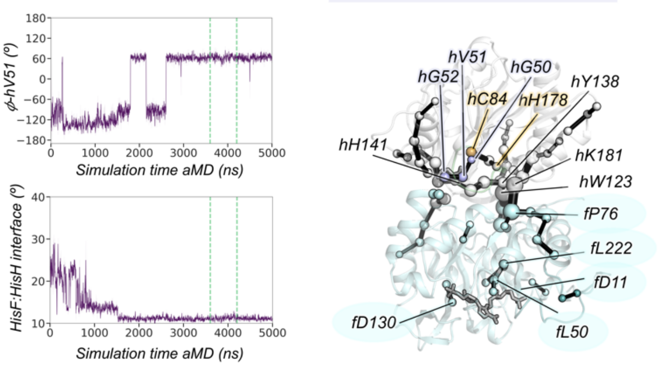


| HisF (28/253) | | | HisH (37/201) | | |
|---|---|---|---|---|---|
| fA3 | fF77 | FV226 | hA23 | hC84 | hF139 |
| fK4 | fK99 | fN247 | hS24 | hA97 | hV140 |
| fR5 | fV100 | | hF27 | hV111 | hH141 |
| fS29 | fF120 | | hE28 | hL118 | hT142 |
| fG30 | fG121 | | hG50 | hP119 | hY143 |
| fV33 | fI129 | | hV51 | hW123 | hF177 |
| fF49 | fD130 | | hG52 | hN124 | hH178 |
| fL50 | fV158 | | hH53 | hE125 | hK181 |
| fD51 | fT194 | | hG55 | hV126 | hS182 |
| fV69 | fT195 | | hE56 | hI127 | hG186 |
| fI73 | fS201 | | hR59 | hF128 | hR187 |
| fI75 | fL222 | | hR62 | hY137 | |
| fP76 | fA224 | | hD65 | hY138 | |

**Fig. S31. Detailed time-evolution Shortest-Path Map (SPM) analysis along a representative 5-μs aMD trajectory.** Plot of the HisF:HisH interface angle along the simulation time. Plot of the $\phi$ dihedral angle of *h*V51. Vertical dashed green lines indicate the range of simulation time where the SPM was calculated. Structural representation of the PRFAR-IGPS SPM analysis with the most relevant residues highlighted. Residues belonging to HisF and HisH subunits are highlighted in cyan and white, respectively. oxyanion strand, Ω-loop, catalytic residues, and loop1 are colored in purple, green, orange, and teal, respectively. List of all residues included in the SPM.

**References**


1. A. Douangamath, *et al.*, Structural evidence for ammonia tunneling across the (βα)8 barrel of the imidazole glycerol phosphate synthase bienzyme complex. *Structure* **10**, 185–193 (2002).
2. A. T. Vanwart, J. Eargle, Z. Luthey-Schulten, R. E. Amaro, Exploring residue component contributions to dynamical network models of allostery. *J. Chem. Theory Comput.* **8**, 2949–2961 (2012).
3. A. T. Van Wart, J. Durrant, L. Votapka, R. E. Amaro, Weighted implementation of suboptimal paths (WISP): An optimized algorithm and tool for dynamical network analysis. *J. Chem. Theory Comput.* **10**, 511–517 (2014).
4. I. Rivalta, *et al.*, Allosteric pathways in imidazole glycerol phosphate synthase. *Proc. Natl. Acad. Sci. U. S. A.* **109**, 8366 (2012).
5. H. G. D.A. Case, R.M. Betz, D.S. Cerutti, T.E. Cheatham, III, T.A. Darden, R.E. Duke, T.J. Giese, C. A.W. Goetz, N. Homeyer, S. Izadi, P. Janowski, J. Kaus, A. Kovalenko, T.S. Lee, S. LeGrand, P. Li, I. Lin, T. Luchko, R. Luo, B. Madej, D. Mermelstein, K.M. Merz, G. Monard, H. Nguyen, H.T. Nguyen, J. S. Omelyan, A. Onufriev, D.R. Roe, A. Roitberg, C. Sagui, C.L. Simmerling, W.M. Botello-Smith, L. X. and P. A. K. R.C. Walker, J. Wang, R.M. Wolf, X. Wu, AMBER 16 (2016).
6. J. Wang, R. M. Wolf, J. W. Caldwell, P. A. Kollman, D. A. Case, Development and testing of a general amber force field. *J. Comput. Chem.* **25**, 1157–1174 (2004).
7. B. H. Besler, K. M. Merz, P. A. Kollman, Atomic charges derived from semiempirical methods. *J. Comput. Chem.* **11**, 431–439 (1990).
8. and D. M. J. Frisch, G. W. Trucks, H. B. Schlegel, G. E. Scuseria, M. A. Robb, J. R. Cheeseman, G. Scalmani, V. Barone, G. A. Petersson, H. Nakatsuji, X. Li, M. Caricato, A. Marenich, J. Bloino, B. G. Janesko, R. Gomperts, B. Mennucci, H. P. Hratchian, J. V. Ort, Gaussian 09, Revision A.02 (2016).
9. J. A. Maier, *et al.*, ff14SB: Improving the Accuracy of Protein Side Chain and Backbone Parameters from ff99SB. *J. Chem. Theory Comput.* **11**, 3696–3713 (2015).
10. W. L. Jorgensen, J. Chandrasekhar, J. D. Madura, R. W. Impey, M. L. Klein, Comparison of simple potential functions for simulating liquid water. *J. Chem. Phys.* **79**, 926 (1998).
11. T. Darden, D. York, L. Pedersen, Particle mesh Ewald: An N·log(N) method for Ewald sums in large systems. *J. Chem. Phys.* **98**, 10089 (1998).
12. D. Hamelberg, J. Mongan, J. A. McCammon, Accelerated molecular dynamics: A promising and efficient simulation method for biomolecules. *J. Chem. Phys.* **120**, 11919–11929 (2004).
13. D. Hamelberg, C. A. F. De Oliveira, J. A. McCammon, Sampling of slow diffusive conformational transitions with accelerated molecular dynamics. *J. Chem. Phys.* **127**, 155102 (2007).
14. G. A. Tribello, M. Bonomi, D. Branduardi, C. Camilloni, G. Bussi, PLUMED 2: New feathers for an old bird. *Comput. Phys. Commun.* **185**, 604–613 (2014).
15. A. Barducci, G. Bussi, M. Parrinello, Well-Tempered Metadynamics: A Smoothly Converging and Tunable Free-Energy Method. *Phys. Rev. Lett.* **100**, 020603 (2008).
16. *,‡ Paolo Raiteri, ‡ Alessandro Laio, ‡ Francesco Luigi Gervasio, § and Cristian Micheletti, M. Parrinello‡, Efficient Reconstruction of Complex Free Energy Landscapes by Multiple Walkers Metadynamics†. *J. Phys. Chem. B* **110**, 3533–3539 (2005).
17. A. Romero-Rivera, M. Garcia-Borràs, S. Osuna, Role of Conformational Dynamics in the Evolution of Retro-Aldolase Activity. *ACS Catal.* **7**, 8524–8532 (2017).
18. A. C. Kneuttinger, *et al.*, Significance of the Protein Interface Configuration for Allostery in Imidazole Glycerol Phosphate Synthase. *Biochemistry* **59**, 2729–2742 (2020).
19. S. Beismann-Driemeyer, R. Sterner, Imidazole glycerol phosphate synthase from Thermotoga maritima. Quaternary structure, steady-state kinetics, and reaction mechanism of the bienzyme complex. *J. Biol. Chem.* **276**, 20387–20396 (2001).
20. J. M. Lipchock, J. P. Loria, Nanometer propagation of millisecond motions in V-type allostery. *Structure* **18**, 1596–1607 (2010).
21. R. A. Boto, *et al.*, NCIPLOT4: Fast, Robust, and Quantitative Analysis of Noncovalent

Interactions. *J. Chem. Theory Comput.* **16**, 4150–4158 (2020).
